# Multi-omics approach to identify bacterial polyynes and unveil their antifungal mechanism against *Candida albicans*

**DOI:** 10.1101/2021.03.30.437786

**Authors:** Ching-Chih Lin, Sin Yong Hoo, Chih Lin, Kai-Fa Huang, Ying-Ning Ho, Chi-Hui Sun, Han-Jung Lee, Pi-Yu Chen, Lin-Jie Shu, Bo-Wei Wang, Wei-Chen Hsu, Yu-Liang Yang

**Affiliations:** Agricultural Biotechnology Research Center, Academia Sinica, Taipei 11529, Taiwan; Institute of Biological Chemistry, Academia Sinica, Taipei 11529, Taiwan; Institute of Marine Biology and Center of Excellence for the Oceans, National Taiwan Ocean University, Keelung 20224, Taiwan

## Abstract

Bacterial polyynes are highly active natural products with a broad-spectrum of antimicrobial activities. However, their detailed mechanism of action remains unclear. Through integrating comparative genomics, transcriptomics, functional genetics, and metabolomics analysis, we identified a unique polyyne resistance gene, *masL* (encoding acetyl-CoA acetyltransferase), from the biosynthesis gene cluster (BGC) dominant for the production of antifungal polyynes (massilin A, massilin B, collimonin C, and collimonin D) in *Massilia* sp. YMA4. Phylogenic and chemotaxonomic analyses characterized the core architecture of bacterial polyyne BGC. The crystallographic analysis of the MasL-collimonin C complex indicated that bacterial polyynes serve as a covalent inhibitor of acetyl-CoA acetyltransferase. Moreover, we confirmed that the bacterial polyynes disrupted cell membrane integrity and inhibited cell viability of *Candida albicans* by targeting ERG10 (homolog of MasL). Overall, understanding of the antifungal mechanism of bacterial polyynes presented herein will be useful for the development of polyynes for fungal infections.

## Introduction

Invasive fungal infections caused by *Candida, Aspergillus, Pneumocystis*, and *Cryptococcus* spp. in humans result in approximately 1.4 million deaths per year worldwide^1^. *Candida albicans* is the most prevalent pathogen among the *Candida* spp., causing an invasive fungal infection called Invasive Candidiasis (IC)^2^. The clinical guidelines for the management of Candidiasis offered by the Infectious Diseases Society of America recommend echinocandin and azole-type drugs as initial therapy for Candidiasis^3^. Echinocandin inhibits fungal cell wall synthesis by targeting 1,3-β-glucan synthase and the azoles interfere with fungal cell membrane formation by inhibiting lanosterol 14α-demethylase^4, 5^. However, more and more azole-resistant *Candida* spp. are being isolated from hospital IC patients due to drug abuse of azoles^6^. Because of the increasing severity of drug resistance and the limited number of clinical drugs currently available for treatment, new types of antifungal agents are urgently required^5, 7^.

Polyynes or polyacetylenes, a substantial class of compounds derived from polyunsaturated fatty acids, contain a conformationally rigid rod-like architecture and an electron-rich consecutive acetylene moiety. Hundreds of polyynes have been discovered, out of which compounds have mostly been isolated from terrestrial plants such as (*3R*)-falcarinol and ichthyothereol^8^. In contrast to polyynes from plant sources, bacterial polyynes contain a distinguished terminal alkyne with conjugated systems, which causes bacterial polyynes to be more unstable. This instability has discouraged surveys of bacterial polyynes using the bioactivity-guided isolation approach. To date, only 12 bacterial polyynes have been recorded in a few species. However, these polyynes have been reported to have a broad spectrum of antimicrobial effects. For instance, cepacin, isolated from *Pseudomonas cepacia* (taxonomically reclassified as a *Burkholderia diffusa*), was reported to have anti-bacterial activity against the majority Gram-negative bacteria, staphylococci, and anti-oomycetal activity against *Pythium ultimum*^9, 10^; collimonins isolated from *Collimonas fungivorans* Ter331^11, 12^ and Sch 31828 isolated from *Microbispora* sp. SCC1438^13^ were reported to have antifungal activity against *Aspergillus niger* and *Candida* spp., respectively. Despite the apparent antibiotic effect of these compounds, the active target and mechanism(s) remain unclear.

Here, we delineated the antifungal mechanism of bacterial polyynes. We used a multi-omic approach to identify the bioactive polyynes of *Massilia* sp. YMA4 and characterized their biosynthesis gene cluster (BGC). By comparing bacterial polyyne BGCs via genome mining, we revealed that bacterial polyynes are antifungal agents that act by targeting the first enzyme of ergosterol biosynthesis, acetyl-CoA acetyltransferase. Crystallographic analysis unveiled the detailed binding model of polyynes to the acetyl-CoA acetyltransferase. This information will be useful in new antifungal drug screening and ligand-based drug design.

## Results

### Transcriptomics analysis reveals polyynes as antifungal agents and their encoding BGC in *Massilia* sp. YMA4

Based on a previous survey, *Massilia* sp. YMA4 has antimicrobial effects against *Staphylococcus aureus, Staphylococcus epidermidis, Paenibacillus larvae*, and the pathogenic fungus, *C. albicans*^14^. In antagonism assay of *Massilia* sp. YMA4 against *C. albicans*, a distinct phenotype showed that the antifungal agent was produced in potato dextrose agar (PDA) medium but not in yeast malt agar (YMA) medium (**Fig. 1a**) and this was further confirmed by disc diffusion assay. Notably, we found that the antifungal metabolites were unstable in the extract and hard to scale up for bioassay using the classic bioactivity-guided isolation approach. Therefore, to mine the antifungal metabolites, a combined transcriptomics and metabolomics approach was used to identify the compounds produced in the two different media (PDA and YMA). First, the circular genome of *Massilia* sp. YMA4 was assembled as 6.33 megabase pairs (Mbp) with 5315 coding sequences (CDSs) by the PacBio sequencing system (**Fig. 1b**). Then transcriptomics analysis of *Massilia* sp. YMA4 cultured in the two media, processed using the Illumina platform was conducted. It showed differential expression with 192 upregulated genes and 226 downregulated genes in PDA compared to YMA (with *P* < 0.05 and |fold-change| > 2, **Supplementary Fig. 1a and Data 1**). Then, we assigned these 418 differentially expressed genes (DEGs) into 77 pathways for pathway analysis using the Kyoto Encyclopedia of Genes and Genomes (KEGG)^15^. The results identified a total of eight significantly enriched pathways involved in different culture conditions (FDR-adjusted *P* < 0.05, **Supplementary Fig. 1b and Data 2**). Compared to YMA, the enriched pathways in PDA were conspicuously associated with small-molecule metabolism, especially fatty acid-related metabolism.

**Fig. 1.**
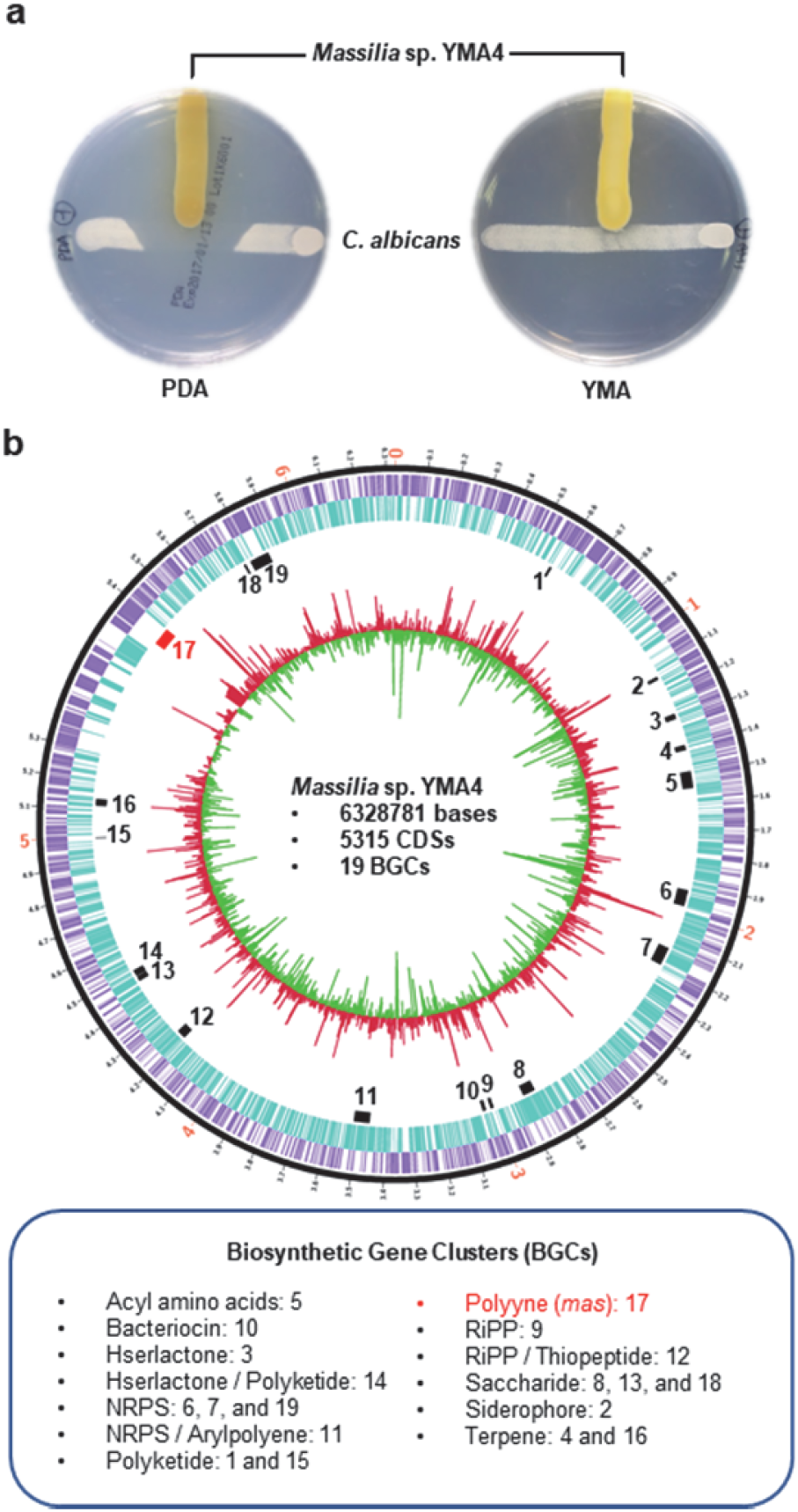
Differentiation of antifungal phenotype and differential expression of biosynthetic gene clusters of *Massilia* sp. YMA4. (**a**) Antagonism assay of *Massilia* sp. YMA4 against *C. albicans* on PDA (active) and YMA (inactive) media. Whole-genome sequence and RNA-seq analysis of *Massilia* sp. YMA4 on PDA (active) and YMA (inactive) media. Megabases are labeled as red on the outer black track; smaller ticks correspond to 100 kbp segments. The circular tracks from outside to inside represent: (1) coding sequences (CDSs) on the forward strand (purple); (2) CDSs on reverse strand (blue); (3) predicted biosynthetic gene clusters (BGCs, black and red) and polyyne BGC (red); (4) fold change histogram of CDSs of *Massilia* sp. YMA4 on PDA versus YMA; red indicates upregulation, and green indicates downregulation.

To mine the biosynthesis genes encoding the unstable antifungal metabolites, an *in-silico* BGC identification combined the results of rule-based antiSMASH (bacterial version, v.5)^16^ and deep learning annotation DeepBGC^17^ to characterize 19 BGCs in *Massilia* sp. YMA4 (annotation list shown in **Supplementary Data 3**). Combining the BGC mining and transcriptomics results, we found that the predicted gene cluster 17 is the only BGC highly and consistently expressed with most genes in PDA compared to YMA. We named the predicted gene cluster 17 as massilin (*mas*) BGC with 12 transcribed genes (*masA* to *masL*). The unique features of *mas* BGC are genes encoding fatty acyl-AMP ligase (*masD*) and acyl carrier protein (*masG*) for fatty acid substrate loading, and modification encoding genes fatty acid desaturases (*masA, masE, masF* and *masH*) and hydrolases (*masI* and *masK*) (**Supplementary Table 1**).

### Characterization of *mas* BGC producing polyynes by *Massilia* sp. YMA4

To identify the metabolites of *mas* BGC, we constructed a biosynthesis-deficient mutant strain (Δ*masH*) through insertion mutation at the *masH* gene locus in *Massilia* sp. YMA4. Δ*masH* lost antifungal activity against type strain ATCC18804 of *C. albicans* and clinically isolated fluconazole-resistant *C. albicans* and *C*.*tropicali* (**Supplementary Fig. 2**). Next, we conducted target isolation using the differential features identified in the UPLC-DAD-HRMS/MS analysis of wild-type and Δ*masH* (**Supplementary Fig. 3**). Then, we purified four major polyynes from ethyl acetate extract. Their structures were elucidated by high-resolution mass spectrometry (**Supplementary Fig. 3 and 4**) and nuclear magnetic resonance (NMR). Of the four, collimonin C **1** and collimonin D **2** were reported in a previous study isolated from *C. fungivorans* Ter331^11^. A new compound with an ene-triyne moiety was named massilin A **3**, which was identified as a racemate with a hydroxyl group at the C6 position of the unsaturated hexadecanoic acid backbone. Another new compound with an ene-diyne-ene moiety named massilin B **4** was supposed to be the precursor of collimonin C **1** or collimonin D **2**. Notably, massilin B **4** is more chemically stable than other polyynes with a terminal alkyne. The four polyynes were biosynthesized by a *mas* BGC putatively derived from palmitic acid with multiple cycles of desaturation and oxidation modification.

For antifungal activity assay, polyynes with a terminal alkyne moiety showed potent inhibition of *C. albicans* with minimum inhibitory concentrations (MIC): 69.73 μM (collimonin C **1**), 35.24 μM (collimonin D **2**) and 2.40 μM (massilin A **3**). However, massilin B **4** with a terminal alkene moiety had no antifungal activity with MIC > 500 μM (**Table 1** and **Supplementary Fig. 5**). These results imply that the terminal alkyne is an essential functional moiety for the antifungal activity of the polyynes.

**Table 1.** Antifungal activity of polyynes and their inhibition kinetics to MasL of *Massilia* sp. YMA4.

| Polyynes | MIC ($\mu\text{M}$)<sup>a</sup> | K<sub>i</sub> ($\mu\text{M}$)<sup>b</sup> | $k_{inact}$ ($\text{min}^{-1}$)<sup>b</sup> | $k_{inact} / K_i$ ($\mu\text{M}^{-1} \text{min}^{-1}$)<sup>b</sup> |
| --- | --- | --- | --- | --- |
| Collimonin C **1** | 69.73 | 297.10 | 0.09798 | 0.000330 |
| Collimonin D **2** | 35.24 | 42.84 | 0.05208 | 0.001216 |
| Massilin A **3** | 2.40 | 132.10 | 0.03449 | 0.000261 |
| Massilin B **4** | >500 | - | - | - |

<sup>a</sup>Minimum inhibitory concentration (MIC) for *C. albicans* (**Supplementary Fig. 5**).
<sup>b</sup>Experimental details and statistics are provided in the **Supplementary Notes**.

### Phylogenetic analysis of polyyne BGCs and their genetic-chemotaxonomy relationship

A phylogenetic analysis was performed on *mas* BGC and its homologous BGCs from finished sequenced bacterial genomes. We used fully transcribed genes of *mas* BGC as a template to process multiple sequence alignments for mining the polyyne BGCs by using MultiGeneBlast^18^ in the bacterial genome database (BCT, 2020 November, NCBI) and additional genomes of polyyne producing bacteria (**Supplementary Data 4**). The results revealed that polyyne BGCs were discontinued in bacterial phylogeny and appeared in certain genera in bacteria. Among the homologous polyyne BGCs in 56 bacteria genomes, we recognized a consensus region of polyyne BGC with a unique gene cluster architecture: fatty acyl-AMP ligase (FAAL) - 2x fatty acid desaturase (FAD) - acyl carrier protein (ACP) - fatty acid desaturase (FAD) – hydrolases/thioesterase (H/TE).

In view of the conservation of the gene cluster architecture, the concatenated amino acid sequence of the consensus region was used to build a phylogenetic tree, and the bacterial species could be intergraded into 11 leaves (**Fig. 2a** and **Supplementary Data 5**). Based on the reported polyyne structures (**Fig. 2b**), we configured polyyne BGCs into three monophyla: the palmitate-derived polyynes family (**C16**) containing the *Massilia* group (this study, compounds **1**-**4**), the *Collimonas* group (*C. fungivorans* Ter331, compounds **1, 2, 7** and **8**^11^) and the *Burkholderia* group 2 (*B. ambifaria* BCC019, compounds **5** and **6**^9^) with an outgroup of *Streptomyces* group and *Amycolatopsis orientalis*; the Stearate-derived polyynes family (**C18**) contains the *Trinickia* group (*T. caryophylli*, compound **9**^19, 20^), *Burkholderia* group 1 (*B. gladioli* BSR3, compound **9**^20^), *Pseudomonas* group (*P. protegens* Cab57, compounds **10** and **11**^21^) and *Gynuella sunshinyii* (ergoynes A, polyyne derivate^22^); and the uncharacterized monophylum with *Mycobacterium* group and *Nocardia brasiliensis*. The phylogenic and chemotaxonomy relationship suggests that polyyne BGCs might first have evolved with an adaptive mutation for a different substrate-specific family before spreading within the family.

**Fig. 2.**
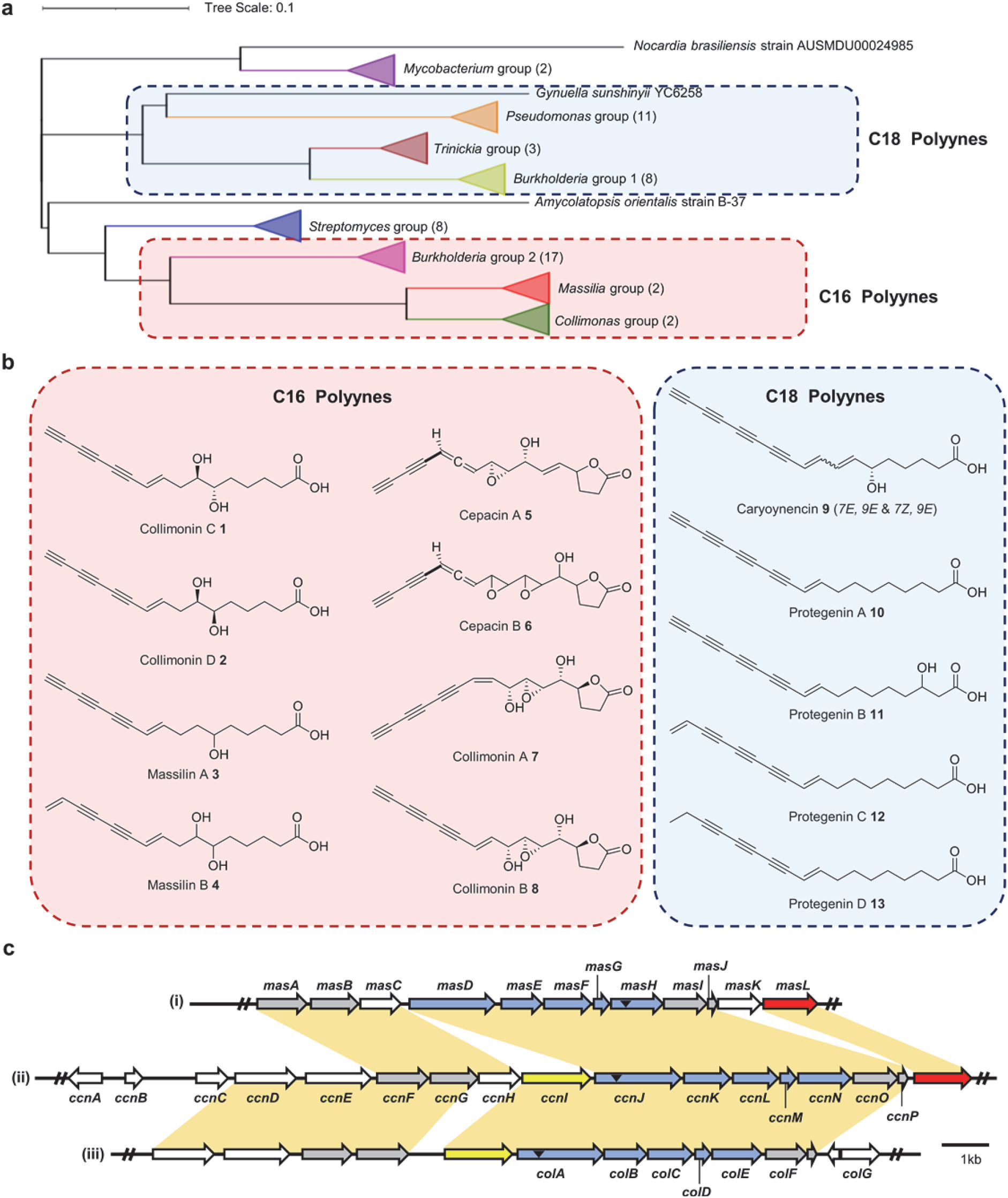
Comparative analysis of polyyne biosynthetic gene cluster (BGC) and structures of bacterial polyynes. (**a**) Phylogenetic analysis of polyyne BGCs in 56 bacteria genomes. The polyyne BGCs were mined in the NCBI BCT database (2020 version) through protein sequence homology using polyyne BGC of *Massilia* sp. YMA4 (*masA* to *masL*) as a query by MultiGeneBlast^18^. Species in the blue boxes have been reported to produce C18 polyynes and species in the red boxes have been reported to produce C16 polyynes. The phylogenetic tree was built with concatenated protein sequences of the gene cluster’s conserved region (*masD* to *masI*) using MUSCLE alignment algorithm and distance estimated with 5000 bootstraps of UPGMA method in MEGA 10^45^. (**b**) The chemical structures of C16 and C18 polyynes ^9, 10, 11, 19, 21^, Collimonin C **1**, collimonin D **2**, and new compounds massilin A **3**, massilin B **4** were found in *Massilia* sp. YMA4. (**c**) Comparison of the polyyne BGC architectures of *Massilia* sp. YMA4 massilins (**i**), *B. ambifaria* BCC0191 cepacins ^9^ (**ii**), and *C. fungivorans* Ter331 collimonins ^11^ (**iii**). Genes conserved in polyyne BGCs across the phylogenetic tree are colored blue and those conserved in C16 polyyne monophylum are colored gray. The potential protective genes in BGC are colored red for acetyl-CoA acetyltransferase and yellow for MFS transporter. The corresponding homolog (over 40% identity) in BGCs between the two species are shown in the green area. Black triangles indicate the mutation sites in previous research and this study.

### MasL serves as a polyyne direct target and has a protective function

We further analyzed the palmitate-derived polyyne (**C16**) monophylum, including *mas* BGC in *Massilia* sp. YMA4 (**Fig. 2c-i**), *ccn* BGC in *B. ambifaria* BCC019 ^9^(**Fig. 2c-ii**), and *col* BGC in *C. fungivorans* Ter331^12^(**Fig. 2c-iii**). The phylogenetic analysis showed that *ccn* BGC branched out before the most recent common ancestor of *mas/col* BGCs, which implies that *ccn* BGC is the evolutionary ancestor of BGC dividing into *mas* and *col* BGCs with a deletion event, independently. Interestingly, the gene encoding major facilitator superfamily (MFS) transporter, which are implicated in multidrug resistance and transport small molecules and xenobiotics^23^, is preserved in *col* BGC but lost in *mas* BGC. In contrast, the *masL*, the acetyl-CoA acetyltransferase gene, remains in *mas* BGC but not in *col* BGC. Antibiotic producers often harbor resistance genes within the antibiotic BGCs for self-protection^24, 25^. On the other hand, drug resistance is also achieved by amplification/overexpression of the drug target^26^. For instance, many fluconazole-resistant strains of *Candida* spp. were reported to have overexpression of the drug target ERG11/CYP51^6^. To evaluate the protective effect of *masL*, we first constructed heterologous expression of *masL* in polyyne-sensitive *C. albicans* (P_*tet*_*- masL*). The expression of *masL* rescued fungal cell viability from polyyne inhibition with MIC (**Fig. 3a**). Furthermore, an *in vitro* MasL inhibition assay showed that polyynes (compounds **1**-**3**) inhibited the MasL enzyme activity (**Table 1**). These results suggest that *masL* serves as a self-resistance gene (SRG) in the *mas* BGC, and in addition that MasL could serve as a direct target of bacterial polyynes for further antifungal mechanistic studies.

**Fig. 3.**
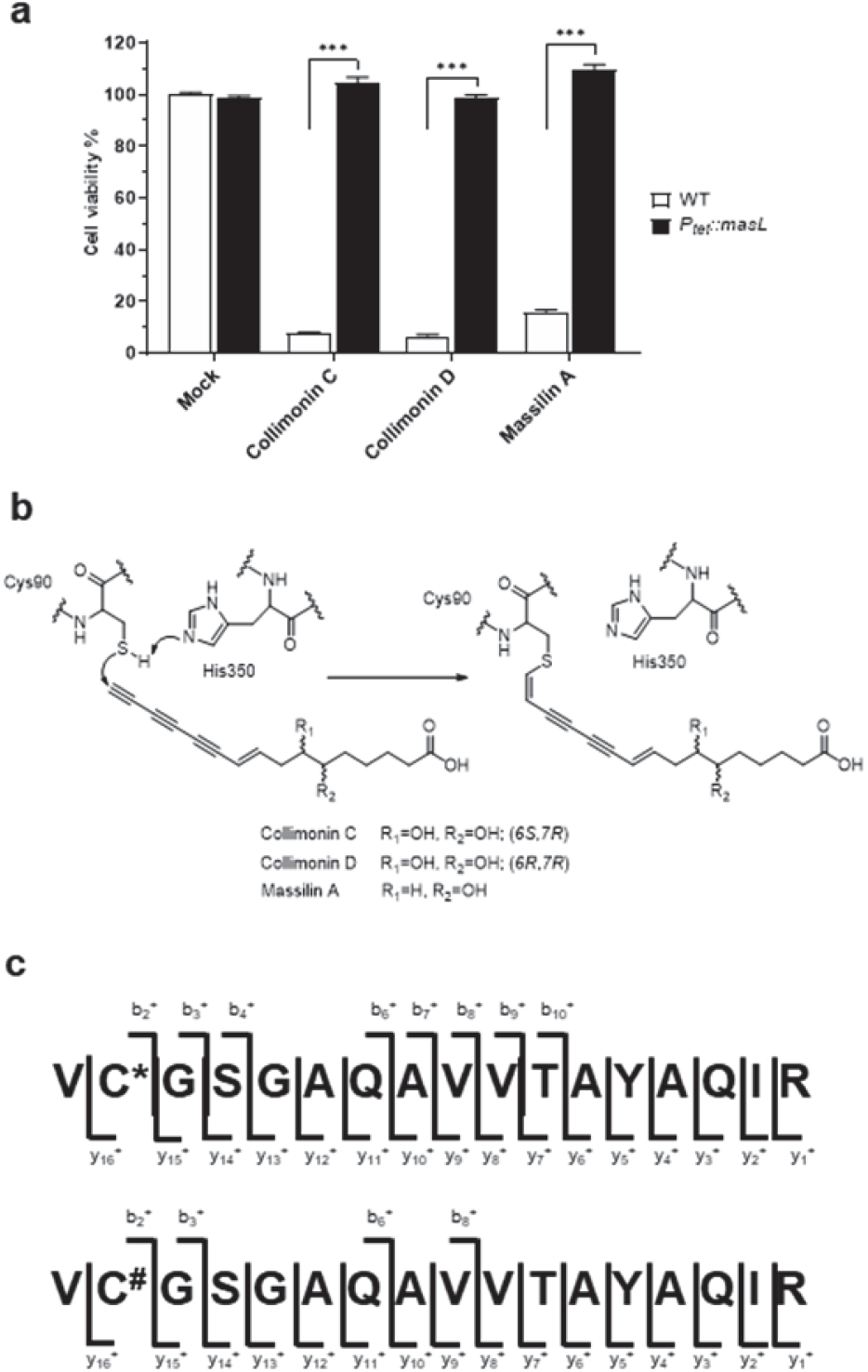
Polyynes as electrophiles for thiol–alkyne addition target MasL active cysteine residue for irreversible covalent inhibition. (**a**) *C. albicans* were rescued by overexpression of *Massilia* sp. YMA4 *masL* from the minimum inhibitory concentration of polyyne treatment. The standard deviation was calculated based on three replicates and the Student t-test was used for statistical analysis. ***, *P* < 0.001. (**b**) The proposed nucleophilic addition mechanism of polyynes (with terminal alkyne) and MasL via S-alkylation of Cys90. (**c**) Mass spectrometry analysis of the polyynes-derived covalent modification on MasL Cys90 (as indicated by Δ mass +258 (asterisks) Da for massilin A **3** and +274 (hash mark) Da for collimonin C/D **1, 2**). (see details in **Supplementary Fig. 8**).

### Polyynes are covalent inhibitors of acetyl-CoA acetyltransferase

The nucleophilic addition by covalent inhibitors targeting of the sulfhydryl group of cysteine residues is the most widely utilized reaction for achieving irreversible binding^27^. For instance, arecoline was reported to be an acetyl-CoA acetyltransferase inhibitor using α,β-unsaturated carbonyl moieties as an electrophile for the sulfhydryl group of reactive Cys126 in ACAT1 protein^28^. Moreover, (*3R*)-falcarinol, which contained an internal diyne moiety isolated from the plant *Daucus carota*, was reported to modify the chemopreventive agent-sensor Keap1 protein at Cys151 covalently^29^. Therefore, we proposed a nucleophilic addition mechanism on polyynes as electrophiles for the reactive cysteine sulfhydryl moiety of MasL (**Fig. 3b**) and confirmed it by mass spectrometry analysis (**Supplementary Data 6**). After incubating collimonin C **1**, collimonin D **2** and massilin A **3** with MasL, respectively, two peptides were observed to have monoisotopic masses (within 5 ppm error) consistent with Cys90 modified by Δmass +258 Da (+C_16_H_18_O_3_) and 274 Da (+C_16_H_18_O_4_) (**Fig. 3c** and **Supplementary Fig. 6**). This observation of collimonin C/D (+C_16_H_18_O_4_)- and massilin A (+C_16_H_18_O_3_)-derived adducts of MasL Cys90 provides convincing evidence of protein S-alkylation via nucleophilic addition to the conjugated terminal alkyne. Consequently, these polyyne inhibitors (compounds **1-3**) represent targeted covalent inhibitors (TCIs) that selectively covalently modify an essential catalytic residue in MasL, leading to irreversible inhibition.

The selectivity of TCI is described reasonably well by the general equation (**Table 1** and **Supplementary Note**). The kinetic study showed that collimonin D **2** has a lower K_I_ (42.84 μM) than massilin A **3** (132.10 μM) and collimonin C **1** (297.10 μM). This suggests that the stereochemistry of the hydroxyl group on polyynes is vital for initial non-covalent complex affinity. We assume that the stereochemistry also affects the reactivity of covalent complex formation for collimonin C **1** with a faster *k*_*inact*_ (0.09798 min^-1^) than collimonin D **2** (0.05208 min^-1^) and massilin A **3** (0.03449 min^-1^). In addition, enzyme inhibition assays for acetyl-CoA acetyltransferase homolog from *C. albicans* ATCC18804 (ERG10_L127S_) and human transition peptide-truncated ACAT1 showed that collimonin C **1**, collimonin D **2**, and massilin A **3** would inhibit the enzyme activity of recombinant ERG10_L127S_ and ACAT1 (**Supplementary Fig. 7**). The mass spectrometry analysis of collimonin C/D- and massilin A-derived adducts of ERG10_L127S_ and ACAT1 also showed the polyynes to be TCIs (**Supplementary Data 6**). The results showed that polyynes would modify the reactive cysteine residues of acetyl-CoA acetyltransferase (Cys90/Cys382 in ERG10_L127S_ and Cys126/Cys413 in ACAT1). Nevertheless, polyynes would also modify other cysteines with a highly nucleophilic sulfhydryl group (Cys166 in ERG10_L127S_ and Cys119/Cys196 in ACAT1) but not every cysteine in protein (**Supplementary Fig. 8 and 9**). Taken together, polyynes as a lead structure are able target the reactive cysteine residues in acetyl-CoA acetyltransferase with certain selectivity.

### The MasL-collimonin C complex shares a similar interaction in the substrate/inhibitor to enzyme binding model

We solved the crystal structures of MasL in its apo and collimonin C-bound forms at 1.78 Å and 1.66 Å resolution, respectively. The asymmetric unit (space group *P*1 for apo MasL and *P*2_1_ for complex) of both structures contains a tetramer of the protein (**Supplementary Fig.10**), as observed in solution (20 mM Tris-HCl pH8.5, 100 mM NaCl). The monomer of MasL shares the general fold architecture reported in the type II biosynthetic thiolase family^30^. MasL consists of three domains: an N-terminal α/β domain (N-domain, residues 1–121 and 251–271), a loop domain (L-domain, residues 122–250), and a C-terminal α/β domain (C-domain, residues 272–394) (**Supplementary Fig.11)**. The N- and C-domains form a typical five-layered fold (α-β-α-β-α) as observed in the structures of other type II biosynthetic thiolases including *Zoogloea ramigera* PhaA^30^, *Clostridium acetobutylicum* CEA_G2880^31^, *Aspergillus fumigatus* ERG10A^32^, and human ACAT1^33^. The L-domain displays an α/β fold with a tetramerization loop associated with the C-domain (**Supplementary Fig.12)**.

Many high-resolution atomic structural models of acetyl-CoA acetyltransferases/type II biosynthetic thiolases have been reported to date. The structures of thiolases from many organisms are similar despite the lack of sequence similarity and acyl-Co A substrate diversity. Moreover, many structural models of the substrate-binding complex revealed the Claisen condensation reaction and binding model within the reaction pocket. In our MasL and its complex model, the substrate-binding pocket was located on the surface of the enzyme facing the opposite dimer of the tetrameric assembly. The pocket was a tunnel shape of ∼10 Å depth with ∼6-8 Å diameter for the linear pantothenic moiety of coenzyme A (CoA) extending through the reactive center. The Claisen condensation reactive center in MasL contained reactive cysteine residues Cys90 and nucleophilic activation residues His350 and Cys380 in the C-domain. In the MasL-collimonin C complex, the conjugated polyyne tail extended into the MasL substrate binding site and formed a covalent bond between the terminal carbon (C16) and the reactive cysteine sulfhydryl moiety of Cys90 (**Fig. 4 and Supplementary Fig. 13**). The observation is consistent with mass spectrometry analysis indicating the irreversible covalent inhibition of polyynes on MasL or acetyl-CoA acetyltransferase/type II biosynthetic thiolases via nucleophilic addition.

**Fig. 4.**
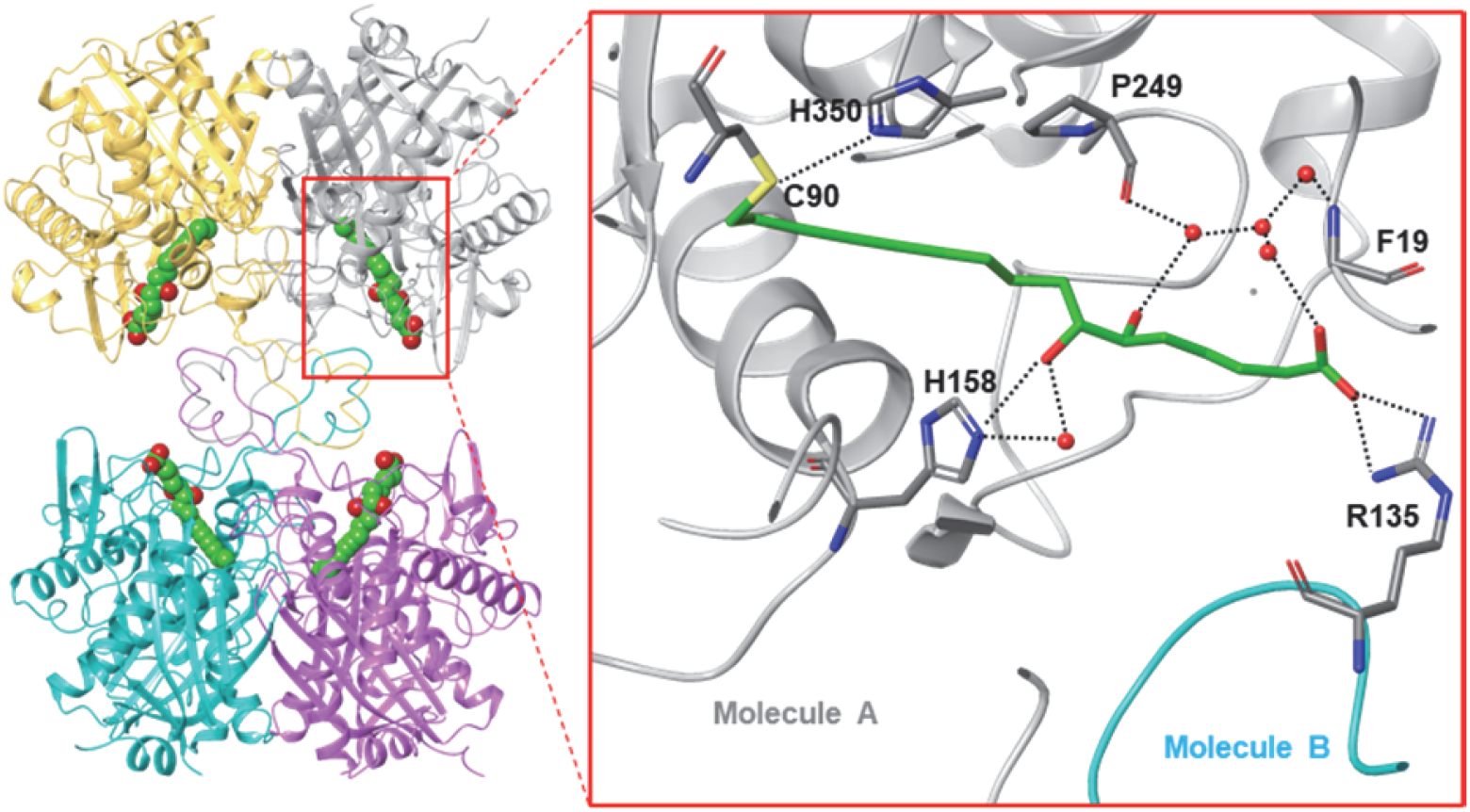
Modeled structures of MasL-collimonin C complex and polar interaction within the binding site. Representative views of the crystal structures of MasL in complex of collimonin C **1**. The residues involved in collimonin C **1** interactions are shown as sticks with sequence identities indicated in the main chain molecule in gray and Arg135 in another molecule in cyan color. The dotted lines indicate the hydrogen bonds and salt bridge involved in collimonin C **1** interactions within the binding pocket.

In the further analysis of the MasL-collimonin C complex, C7-OH of collimonin C **1**, His158 of MasL, and a water molecule formed a strong polar interaction network, including a direct hydrogen bond (3.00-3.16 Å) and a water-mediated hydrogen bond between C7-OH and His158 (**Fig. 4**). The superimposition of four monomers of the MasL-collimonin C complex showed that C6-OH of collimonin C **1** had more flexibility on the spatial direction (with a dihedral angle to C7-OH from 109° to 170°) and built a sophisticated polar interaction network with the amide of Pro249 in the panthetheine loop and multiple water molecules (**Supplementary Fig. 14)**. In the substrate-binding model of the thiolases, the conserved histidine residue on the covering loop formed a water-mediated hydrogen bond to the carbonyl moiety in the pantothenic part of CoA. Also, one or more water molecules mediated the hydrogen-bonding network between the hydroxyl moiety in CoA and backbone amide moieties in the panthetheine loop in thiolase^34^. The polar interacting residues for the collimonin C **1** (inhibitor) binding were similar to CoA (substrate) in other thiolase models therefore the collimonin C **1** competitively bound into the reaction pocket.

The superimposition of the inhibitor/substrate binding models, including *A. fumigatus* ERG10A (pdb code **6L2C**^32^, chain A; identity 37.8%), Human ACAT1 (pdb code **2IBU**^33^, chain A; identity 36.8%), *C. acetobutylicum* CEA_G2880 (pdb code **4XL4**^31^, chain A; identity 48.9%) and *Z. ramigera* PhaA (pdb code **1QFL**^30^, chain A; identity 44.6%), showed that collimonin C **1** could align well with the phosphate-pantothenic part of CoA (**Fig. 5**). The hydrogen bond between C7-OH of collimonin C **1** and His158 of MasL was well superimposed with the polar interaction of carbonyl moiety in CoA. Even though the superimposition between C6-OH of collimonin C **1** and the α-hydroxy pantoic acid moiety showed a slightly different polar network orientation due to hydroxyl moiety flexibility, the polar interaction was still conserved in the substrate/inhibitor binding model. The crystallographic analysis and *in vitro* thiolase activity assay demonstrated that the configuration of hydroxyl moiety of polyynes is vital for enzymatic affinity.

**Fig. 5.**
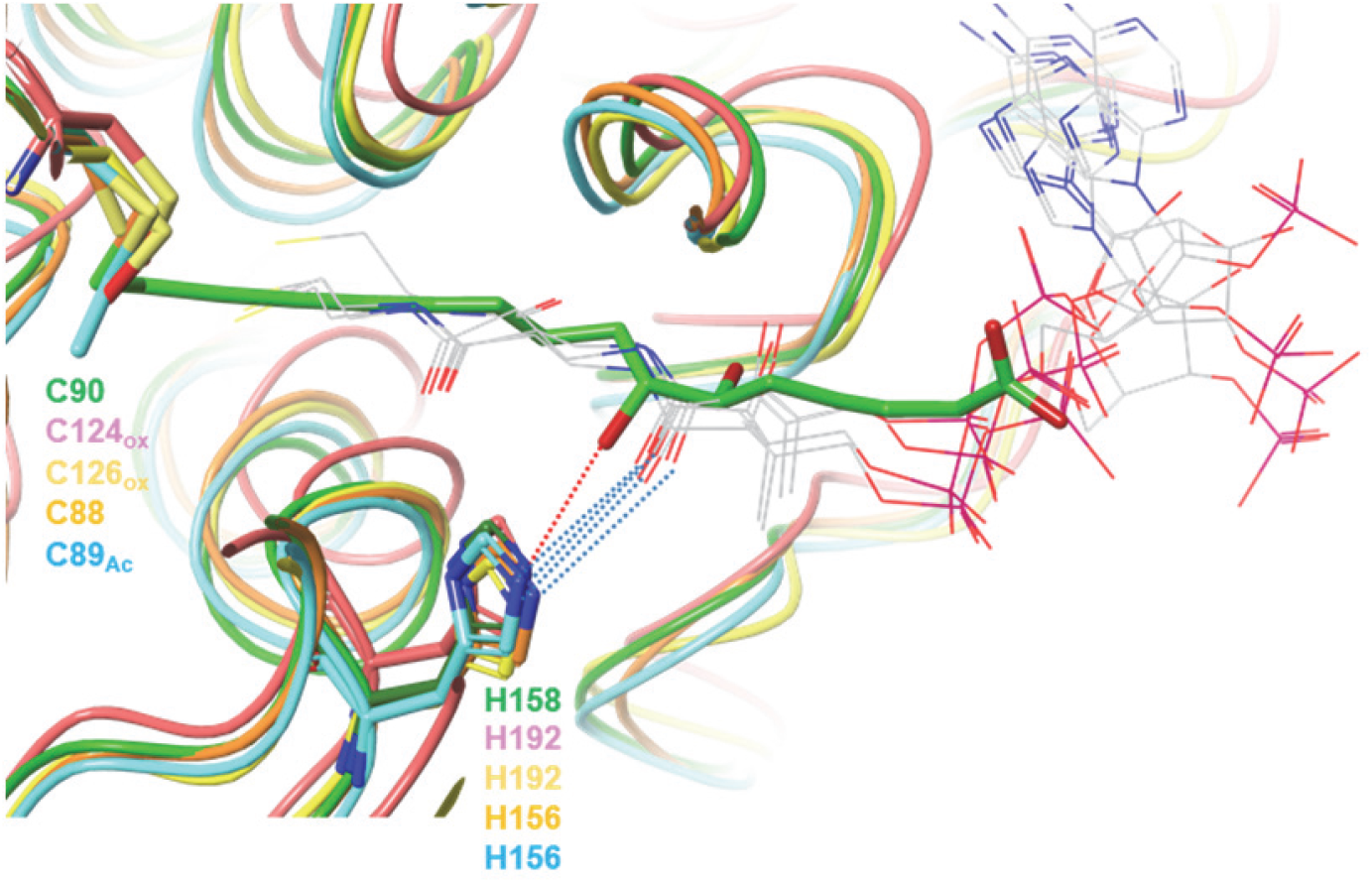
Structure comparison with ligand-bound acetyl-CoA acetyltransferase. Superposition of enzyme monomers of *Massilia* sp. YMA4 MasL bound to Collimonin C **8** (pdb code **7EI4**, chain A; green) and *A. fumigatus* ERG10A (pdb code **6L2C**, chain A^32^; salmon), Human ACAT1 (pdb code **2IBU**, chain A^33^; yellow), *C. acetobutylicum* CEA_G2880 (pdb code **4XL4**, chain A^31^; orange), *Z. ramigera* PhaA (pdb code **1QFL**, chain A^30^; cyan) bound to CoA. The conserved histidine residues involved in polar interactions are shown as sticks, with sequence identities colored the same as the backbone shown as a ribbon. All CoA ligands are shown with carbon atoms in gray as lines; while collimonin C **1** is shown with carbon atoms green as sticks. The red dotted line indicates the hydrogen bonds in the MasL-collimonin C complex. The blue dotted lines indicate the water-mediated polar interactions between CoA and selected histidine residues. Abbreviations for active cysteine modification: Ox, oxidized form of the cysteine thiol group (sulfenic acid type); Ac, acetylation of the cysteine residue.

Surprisingly, although there was no significant induced-fit within the pocket, the collimonin C **1** caused the Arg135 on the tetramerization loop to swap to form a salt bridge across the two subunits within the binding site (**Supplementary Fig. 15)**. This finding was similar to the CoA-bound thiolase in *C. acetobutylicum* CEA_G2880 (pdb code **4XL4** ^31^), in which the Arg133 in CEA_G2880 formed a hydrogen bond to the phosphate moiety of CoA. The salt bridge/hydrogen bond formation on the arginine in thiolase would increase the binding affinity and suggests that collimonin C **1** would stabilize the tetramer of MasL rather than disrupting the tetramerization of ACAT1 as arecoline inhibition^28^.

### Acetyl-CoA acetyltransferase as an antifungal target

In this study, we confirmed that collimonin C **1**, collimonin D **2**, and massilin A **3** would inhibit the enzyme activity of acetyl-CoA acetyltransferase homolog ERG10_L127S_ from *C. albicans* through covalent competition on the substrate binding site and the reactive cysteine residue. As acetyl-CoA acetyltransferase is the first enzyme to catalyze acetoacetyl-CoA formation for mevalonate biosynthesis, inhibition of ERG10_L127S_ would block the mevalonate production and subsequently disrupt the squalene and ergosterol biosynthesis. Among the clinical antifungal drugs, azole drugs inhibit ergosterol biosynthesis by targeting the critical biosynthesis enzyme ERG11 and causing the dysfunction of maintenance of fluidity, permeability, and structural integrity of fungal cell membrane^4^. Moreover, the reduced expression of acetyl-CoA acetyltransferase homolog ERG10A in *A. fumigatus* led to severe morphological defects and increased susceptibility to oxidative and cell wall stresses^32^. Therefore, we carried out a transmission electron microscopy experiment and observed that polyynes disrupt the cell membrane structure (**Fig. 6a**).

**Fig. 6.**
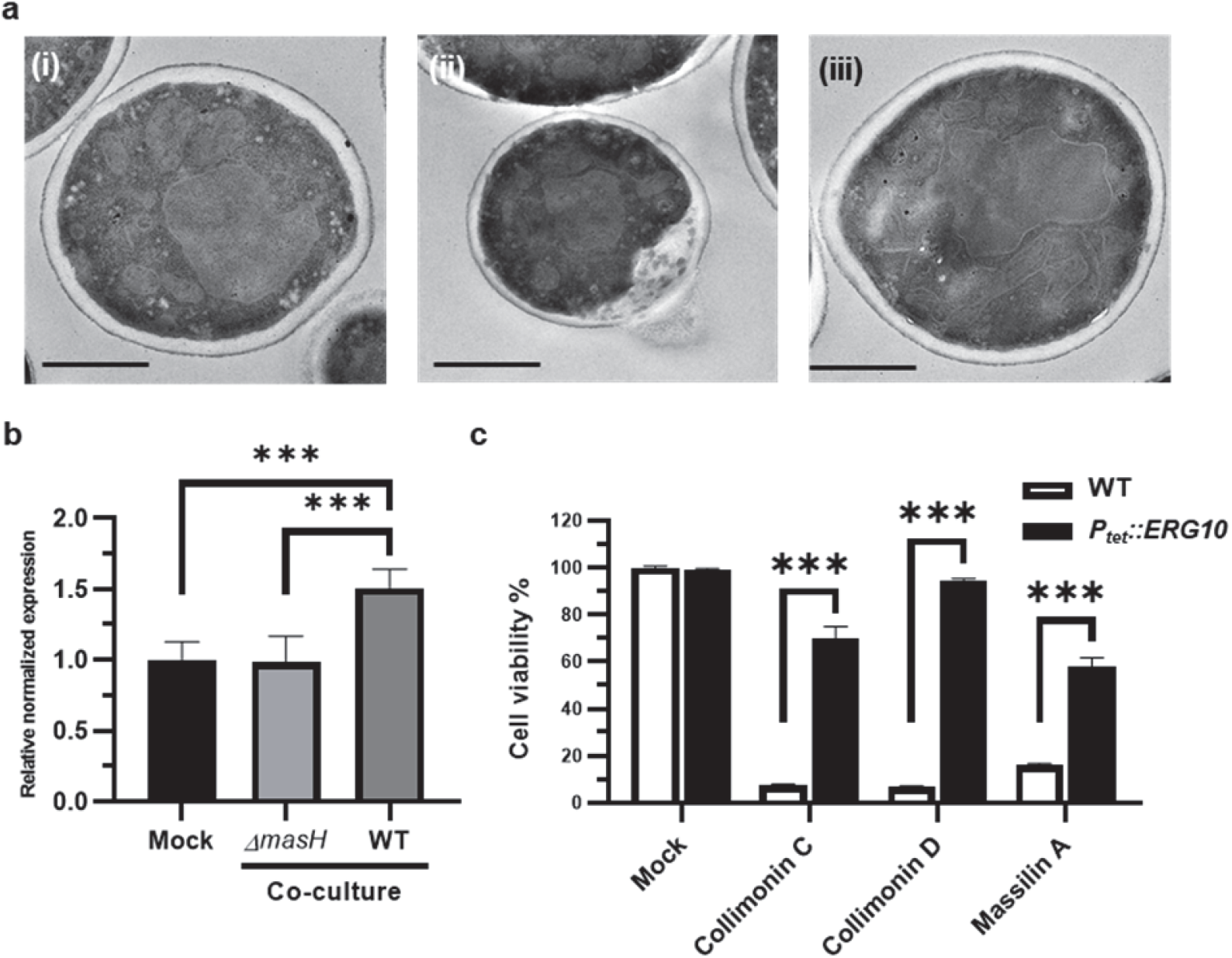
Polyynes inhibit *C. albicans* through disruption of cell membrane stability. (**a**) Transmission electron microscopy images of *C. albicans* cells treated with Mock (**i**) and 1 mg/mL *Massilia* sp. YMA4 ethyl acetate crude extract (**ii**). Cells treated with 1 mg/mL *ΔmasH* ethyl acetate crude extract (**iii**) were used as a negative control. Scale bar, 1 µm. (**b**) Gene expression of *ERG10* in *C. albicans* co-cultured with *ΔmasH* and *Massilia* sp. YMA for two days revealed by real-time qPCR. Mock represents *C. albicans* growth alone. *C. albicans Act1* was used for internal normalization, and *ERG10* expression levels were further normalized to the Mock condition. The standard error of the mean (SEM) was calculated based on at least three replicates and the Student t-test was used for statistical analysis. ***, *P* < 0.001. *C. albicans* are rescued by overexpression of *C. albicans ERG10* from the minimum inhibitory concentration of polyyne treatment (collimonin C/D **1, 2**, and massilin A **3**). The standard deviation was calculated based on three replicates and the Student t-test was used for statistical analysis. ***, *P* < 0.001.

Meanwhile, we detected that *ERG10* gene expression was upregulated during co-culture of *Massilia* sp. YMA4 with *C. albicans* (**Fig. 6b**). Furthermore, we constructed the *ERG10* overexpression strain of *C. albicans* (P_*tet*_*-ERG10*). We found that the overexpression of *ERG10* could rescue the fungal cell viability from polyyne inhibition with MIC as a protective effect of heterologous expression of *Massilia* sp. YMA4 *masL* in *C. albicans* (**Fig. 6c**). Taken together, these results revealed the antifungal mechanism of the polyynes (compounds **1-3**) through targeting the acetyl-CoA acetyltransferase ERG10, resulting in *C. albicans* inducing expression of *ERG10* for tolerance to *Massilia* sp. YMA4 attack during fungal-bacterial interaction.

### Discussion

After the Waksman platform was first introduced in the 1940s^35^, many natural antibiotics were systematically discovered in the chemical crosstalk of microbe-microbe interaction. Even though recent technology can rapidly explore the metabolites hidden in the interaction between host and effector, environmental factors have a significant impact on antibiotic production resulting in various bioactive spectra, which should be considered in practical surveys. *Massilia* sp. YMA4 showed differential antifungal activity in response to different culture conditions. However, the antifungal activity of *Massilia* sp. YMA4 extract was too unstable to identify the active metabolites using the general bioactivity-guide isolation approach. Because of this, we instead combined transcriptomics, functional genetics, and metabolomics analyses to reveal the *mas* BGC and its products— unstable bacterial polyynes—as the principal antifungal agents of *Massilia* sp. YMA4. We succeeded in identifying two new compounds, massilin A **3** and B **4**, and two known compounds, collimonins C **1** and D **2**. Massilin B **4**, the potential precursor of collimonin C **1** or D **2**, did not contain the terminal alkyne and was more stable than other terminal alkyne-containing polyynes but lost its antifungal activity. This chemical property had been mentioned in the study of *Pseudomonas protegens*, which protegenins C **12** and D **13** without terminal alkyne are more stable than protegenins A **10** and B **11** but the antioomycete activities against *P. ultimum* are greatly reduced ^21^.

Notably, the critical terminal alkyne of bacterial polyynes had been mentioned in the study of *Pseudomonas protegens*. This implies that the terminal alkyne of bacterial polyynes is a prerequisite for bioactivity and a contributor to their instability, resulting in self-polymerization.

Previous reports of polyyne BGCs discovered through transposon mutagenesis gave partial information for characterizing the complete BGC architecture^9, 11^. During the genome mining of *mas* BGC, we failed to use the rule-based genome mining tools (antiSMASH^16^) to recognize the cluster information due to the lack of defined polyyne BGC information. However, the deep learning genome mining tool (DeepBGC ^17^) classified *mas* BGC as a type II fatty acid/polyketide synthase (FA-PKS) BGC ^36^ with indicative features, such as fatty acyl-AMP ligase (FAAL), an acyl carrier protein (ACP), fatty acid desaturases (FADs) and hydrolases/thioesterase (H/TE). Subsequently, combining the genome mining and transcriptomics analysis results, the putative BGC was correlated to complete the characterized *mas* BGC. Further information about the polyyne BGCs was attained out by blasting multiple homologs using MultiGeneBlast^18^ with a fully transcribed *mas* BGC as a query. The BGC mining results helped us figure out the core biosynthesis architecture in polyyne BGC as an arranged feature of FAAL-2x FAD-ACP-FAD-H/TE. The subsequent phylogenetic analysis for the polyyne BGCs by conserved core genes combined with chemotaxonomy revealed that a potential evolutionary event, substrate-specific functional evolution (palmitate and stearate) occurred prior to spreading inter-species.

Antibiotics encoding BGCs are important as a defensive strategy for microbial survival. Plasmids are common carriers for BGC transformation between bacteria to gain functional genes. In addition, horizontal gene transfer (HGT) ^37^ is another strategy of gene transfer and usually occurs in bacteria to gain function to defeat enemies. Regarding the relationship within a sister group of polyyne BGCs in palmitic-derived monophylum (*ccn* encoding cepacins and *col* encoding collimonins), HGT events hypothetically transmitted polyyne BGC from *ccn* BGC into *mas* and *col* BGCs, independently. Then, a deletion event occurred with the result that the *ccn* BGC independently divided into *mas* BGC and *col* BGC each of which contained a different self-protection mechanism. The *col* BGC preserved the MFS transporter and, in contrast, *mas* BGC kept the acetyl-CoA acetyltransferase (MasL) for detoxification in polyyne production.

In drug-target surveys, the inhibitor target has sometimes been found to serve a protective function to resist the inhibitor^38, 39^. In this study, we identified the acetyl-CoA acetyltransferase MasL as the direct target of the polyynes, and the homolog ERG10 in *C. albicans* could gain resistance by overexpression. This suggests that ERG10 is the antifungal target of polyynes disrupting the mevalonate and downstream ergosterol biosynthesis, and then abolishing the cell membrane integrity. The success in this case reintroduces the notion that drug targets can be discovered from screening the SRG in gain-of-function assay. Moreover, inhibition of human mitochondrial acetyl-CoA acetyltransferase ACAT1 by the bioactive polyynes (compounds **1-3**) suggested that polyynes would be a species-wide inhibitor of acetyl-CoA acetyltransferases/type II biosynthetic thiolases.

The mevalonate pathway metabolites are essential for cancer cell survival and growth, for example, ketogenesis is associated with prostate cancer progression^40^. Likewise, statins, the hypercholesterolemia drugs also showed anticancer effects on stem cell-like primary glioblastoma by inhibiting HMG-CoA reductase in mevalonate biosynthesis^41^. ACAT1, the first enzyme of the mevalonate biosynthesis pathway, was reported to be an important factor for tumor growth in multiple cancer cell lines^28^. As we revealed that bacterial polyynes could inhibit human mitochondrial ACAT1, it would be worth exploring the anticancer potential of bacterial polyynes in the future.

To date, acetyl-CoA acetyltransferase inhibitors have usually been designed as CoA substrate derivatives or analogs (**Supplementary Fig.16**). Notably, the binding affinity (Km) of acetyl-CoA acetyltransferase with CoA-derivate substrates ranges from 3.8 μM to 1.06 mM ^32^. Compared to previous analog inhibitor reports, in which K_I_ ranged from 1.4 μM to 15 mM and *k*_*inact*_ ranged from 0.26 min^-1^ to 4 min^-1^, bioactive polyynes in our study inhibited MasL with an equal level of binding affinity (K_I_ from 42.84 μM to 297.10 μM) but lower reaction rate (*k*_*inact*_ from 0.03 min^-1^ to 0.1 min^-1^)^42, 43, 44^. These data suggest polyynes may be a potential lead structure for drug design.

In covalent drug design, inhibitors with an electrophile moiety, such as nitrile, alkyne, acrylamide, epoxide, or α,β-unsaturated carbonyl, are the major resources for covalent bond formation to the nucleophilic moieties^27^. For example, falcarindiol was reported to have S-alkylation at Cys151 in Keap1 protein^29^; however, it lacks an actual bond formation mechanism. Polyynes are a group of high electron enriched metabolites that usually react to nucleophilic moieties. In the MasL-collimonin C complex model, the terminal alkyne of polyynes was used to elaborate bond formation with the sulfhydryl moiety in MasL Cys90. Furthermore, we revealed structurally detailed substrate/inhibitor binding models of the thiolases. The superimposition of the MasL-collimonin C complex and the other CoA-thiolase complexes showed collimonin C **1** and CoA shared a similar polar interaction to bind to the thiolases. Additionally, regarding the salt bridge/hydrogen bond formation within the binding site, the induced-fit arginine/lysine residue was conserved in procaryotic species but not in eukaryotic homologous thiolases. This supposedly causes a different affinity in homologous thiolases and could highlight ligand-based drug design with species selectivity.

In summary, we used an integrated strategy to unveil the biosynthesis and antifungal mechanism of bacterial polyynes. A well-characterized core architecture of bacterial polyyne BGC was attained which allowed the exploration of new bacterial polyynes further using genome mining. We illustrated the antifungal mechanism of collimonin C **1**, collimonin D **2**, and massilin A **3** through inhibiting the acetyl-CoA acetyltransferase ERG10 in *C. albicans*. The crystallographic analysis provided detailed structural insight into the MasL-collimonin C complex, which will provide useful information for designing new inhibitors of acetyl-CoA acetyltransferase. These results will help future research in bacterial polyyne mining, biosynthesis, and the structure-activity relationship to develop new antifungal or anticancer drugs.

## Methods

### Genome mining and phylogenetic analysis of polyyne biosynthesis gene clusters

The biosynthesis gene clusters (BGCs) in the genome of *Massilia* sp. YMA4 were characterized via command-line program DeepBGC^17^ and online software antiSMASH^16^ with default settings, and integrated with the criteria: antiSMASH score > 1500, DeepBGC score > 0.7. Then, the *mas* BGC of *Massilia* sp. YMA4 was used to discover the homologous gene clusters in bacteria species using MultiGeneBlast^18^. The database was built with a bacterial sequences database (BCT, 2020 December 01) and whole-genome sequences of polyyne-reported bacterial species from NCBI. A total of 56 bacteria with polyyne BGC (Cumulative Blast bit score > 1500) were found. The homologous protein sequences of each bacterial polyyne BGC were respectively concatenated (total of five amino sequences, starting from MasD homolog to MasI homolog). The concatenated protein sequences were used for alignment (MUSCLE) and the distance (UPGMA, bootstrap 5000 times) between 56 bacteria with *Massilia* sp. YMA4 was identified for phylogenetic tree construction. The analysis was completed by using MEGA 10 with default parameters^45^. iTOL was used to present the results of phylogenetic analysis^46^.

### Mass spectrometry analysis and peptide mapping of polyyne-labeled peptides in MasL, ERG10_L127S,_ and ACAT1

Incubation mixture (20 μL) containing 2 μM protein in 50 mM Tris pH 8.5, 100 mM NaCl, was incubated with 40 μM collimonin C **1**, collimonin D **2**, or massilin A **3** at 25°C for 3 h. The reaction was quenched by adding 4x Laemmli sample buffer (Bio-Rad, USA) with 5 mM DL-dithiothreitol and the protein was separated using SDS-PAGE. The in-gel trypsin digestion was performed with a substrate-to-enzyme ratio of 25:1 (w/w), and the mixture was incubated at 37°C for 20 h^47^. The resultant peptide mixtures were dried and frozen at -20°C until separation by reverse-phase nanoUPLC-ESI-MS. The tryptic peptides were re-dissolved in 10 μL of 0.1% formic acid. An LC-nESI-Q Exactive mass spectrometer (Thermo Scientific, USA) coupled with an online nanoUPLC (Dionex UltiMate 3000 Binary RSLCnano) was used for analysis. An Acclaim PepMap 100 C18 trap column (75 µm x 2.0 cm, 3 µm, 100 Å, Thermo Scientific, USA) and an Acclaim PepMap RSLC C18 nanoLC column (75 µm x 25 cm, 2 µm, 100 Å) were used with a linear gradient from 5% to 35% of acetonitrile in 0.1% (v/v) formic acid for 40 min at a flow rate of 300 nL/min. The MS data were collected in the data-dependent mode with a full MS scan followed by 10 MS/MS scans of the top 10 precursor ions from the full MS scan. The MS scan was performed with 70,000 resolution over the mass-to-charge (*m/z*) range 350 to 1600, and dynamic exclusion was enabled. The data-dependent MS/MS acquisition was performed with a two *m/z* isolation window, 27% normalized collision energy, and 17,500 resolution.

The data were processed using Proteome Discoverer (version 2.4; Thermo Scientific, USA), and the peptides were identified by searching the MS/MS spectra against the MasL, ERG10_L127S,_ and ACAT1 using the Mascot search engine (version 2.3; Matrix Science, UK) and SEQUEST search engine^48^. Cysteine alkylation was used as a dynamic modification, and the modification *m/z* values were +274.121 (+C_16_H_18_O_4_ for collimonin C/D) and +258.126 (+C_16_H_18_O_3_ for massilin A), respectively. For identification, the false discovery rate was set to 0.01 for peptides, proteins, and sites. The minimum peptide length allowed was four amino acids, precursor mass tolerance for 10 ppm, and fragment mass tolerance for 0.02 Da.

### Enzymatic inhibition assay and inhibition kinetics of polyynes

The enzymatic inhibition assay was initiated by adding 50 μM polyynes (collimonin C **1**, collimonin D **2**, and massilin A **3**) into 10 μM ERG10_L127S_ or ACAT1 at 25°C for 1 h. The residue active enzyme reaction started by adding 10 mM acetyl-CoA for another 1 h at 25°C in a total of 12 μL volume with the following concentrations: 8.33 μM enzymes, 41.65 μM polyynes, and 1.67 mM acetyl-CoA. The reaction was quenched by adding 1 μL of 1% formic acid. The monitor method of releasing CoA using a fluorescent probe (7-diethylamino-3-(4-maleimidophenyl)-4-methylcoumarin, CPM) was modified from previous research^49^. The released CoA was used to represent the residual activity or protein occupancy. After 10 min, the pH value was adjusted by 2 μL 0.1 M Tris pH10 and 100 μM CPM probe was added in a total volume of 105 μL for 30 min reaction at 30°C followed by detection of the fluorescent signal using a BioTek Synergy H1 microplate reader (excitation 355 nm; emission 460 nm). Relative fluorescence intensity was obtained by subtracting the fluorescence intensity of the polyyne-free reaction system.

To measure the inhibition kinetics of the polyynes to MasL, different polyyne concentrations as indicated were reacted with the protein for inhibition reaction and then enzymatic reaction as described above. Protein occupancy and inhibition kinetic calculations were performed using GraphPad Prism8 (GraphPad Software, USA; see details in the **Supplementary Notes**).

### Protein Crystallization, Data Collection, Processing, and Refinement

For MasL-collimonin C complex preparation, 20 µM MasL was incubated with 100 µM collimonin C in 20 mM Tris-HCl pH 8.5, and 100 mM NaCl. The MasL-collimonin C complex was purified with a gel-filtration (Superdex 200 Increase 10/300) column.

A freshly thawed aliquot of MasL and MasL-collimonin C complex was concentrated to 20 mg/ml for an initial crystallization screening of ca. 500 conditions (Academia Sinica Protein Clinic, Academia Sinica). The crystallization conditions were manually refined to the final conditions: for MasL, 2% (v/v) Tacsimate pH 7.0, 16% (w/v) polyethylene glycol 3,350, and 0.1 M HEPES, pH 7.5; for MasL-collimonin C complex, 20% (w/v) polyethylene glycol 3,350 and 0.2 M tri-lithium citrate, pH 8. The crystals were grown at 10°C by mixing the protein aliquot with an equivalent volume of crystallization buffer via the hanging drop vapor-diffusion method. For X-ray data collection, the crystals were immediately flash-frozen in liquid nitrogen after dipping into cryoprotectant composed of crystallization solution supplemented with 10% (v/v) glycerol.

X-ray diffraction experiments were conducted at 100K at the TLS beamline 15A or the TPS beamline 05A of the National Synchrotron Radiation Research Center (Hsinchu, Taiwan) with a wavelength of 1 Å. All diffraction data were processed and scaled with the HKL-2000 package^50^. The data collection statistics are listed in **Table 2**. The resulting MasL crystals had a space group of *P*1 with four MasL molecules in an asymmetric unit and a solvent content of ca. 56%. The MasL-collimonin C complex crystals had a space group of *P*2_1_ with one asymmetric unit containing four MasL molecules and a solvent content of ca. 51%^51^.

**Table 2.** Data collection and refinement statistics.

|  | MasL | MasL-collimonin C complex |
| --- | --- | --- |
| **Data collection** |  |  |
| Space group | *P*1 | *P*2<sub>1</sub> |
| Cell dimensions |  |  |
| *a*, *b*, *c* (Å) | 71.786, 76.995, 98.082 | 59.139, 115.077, 125.789 |
| α, β, γ (°) | 79.77, 79.00, 62.98 | 90.00, 91.08, 90.00 |
| Resolution (Å) | 30.0-1.78 (1.84-1.78) | 30.0-1.66 (1.72-1.66) |
| *R*<sub>merge</sub> | 0.045 (0.233) | 0.082 (0.560) |
| *I* / σ*I* | 23.8 (5.0) | 17.7 (2.0) |
| Completeness (%) | 92.3 (85.2) | 99.4 (94.4) |
| Redundancy | 3.8 (3.9) | 5.6 (4.8) |
| **Refinement** |  |  |
| Resolution (Å) | 29.9-1.78 | 29.2-1.66 |
| No. reflections | 160,857 | 180,941 |
| *R*<sub>work</sub> / *R*<sub>free</sub> | 0.128/0.180 | 0.114/0.162 |
| No. atoms |  |  |
| Protein | 11,522 | 11,544 |
| Ligand/ion | - | 80 |
| Water | 1,663 | 1,720 |
| *B*-factors |  |  |
| Protein | 22.3 | 18.3 |
| Ligand/ion | - | 44.0 |
| Water | 34.9 | 34.1 |
| R.m.s. deviations |  |  |
| Bond lengths (Å) | 0.008 | 0.009 |
| Bond angles (°) | 1.455 | 1.449 |

\* Highest-resolution shell is shown in parentheses

The structures of MasL and MasL-collimonin C complex were solved by the molecular replacement method with the program Molrep^52^ using the structure of thiolase from *Clostridium acetobutylicum* (pdb code **4N44**) as the search model. Computational model building was conducted with ARP/wARP or Buccaneer^53, 54^, and the rest of the models were manually built with Coot.^55^ The resulting models were subjected to computational refinement with Refmac5.^56^

The collimonin C and well-ordered water molecules were located with Coot. The stereochemical quality of the refined models was checked with MolProbity^*57*^. Finally, the MasL and MasL-collimonin C complex’s refinement converged at a final *R* factor/*R*_free_ of 0.128/0.180 and 0.114/0.162, respectively. The final refinement statistics are listed in **Table 2**. The refined models of MasL and MasL-collimonin C complex were deposited in the Protein Data Bank with pdb codes **7EI3** and **7EI4**, respectively. The molecular figures were produced with Maestro (**Schrödinger Release 2021-1**: Maestro, Schrödinger, LLC, USA).

### Minimum inhibitory concentration determination and genetic rescue assay

The minimum inhibitory concentration (MIC) measurement was modified from R J Lambert’s method^58^. Different concentrations (300.00, 150.00, 75.00, 37.50, 18.75, 9.38, 4.69, 2.34, 1.17, 0.59, 0.29 µM) of collimonin C **1**, collimonin D **2**, massilin A **3**, atorvastatin, and amphotericin B were prepared in yeast extract-peptone-dextrose (YPD). The *C. albicans* cell viabilities were seeded with initial O.D. 0.05 (600 nm) and incubated at 37°C. After 24 h incubation, the final O.D. was recorded by Epoch 2 Microplate (BioTek Instruments, USA) for MIC calculation.

For genetic rescue assay, the *ERG10* overexpression and *masL* heterologous expression strains were seeded with O.D. 0.05 at 600 nm in YPD treated with MIC of each polyyne at 37°C and supplied with 40 µg/mL doxycycline for gene expression. The *C. albicans* cell viabilities were recorded at 24 h.

The experimental results include at least three biological replicates, and the cell viabilities were normalized to the mock treatment. The statistical results were analyzed using GraphPad Prism 8 (GraphPad Software, USA) with multiple t-test analyses (FDR < 0.05). The MIC of polyynes was built with cell viability (%) of different concentrations, fitting into the modified Gompertz function^58^.

## Data availability

The genome was deposited into the NCBI BioProject database under accession PRJNA476678. The raw-reads of RNA sequencing were deposited at the Sequencing Read Archive (BioProject: PRJNA706894). All LC-MS data used in this paper are publicly available at the GNPS-MassIVE repository under the accession MSV000087007. The raw data of bottom analysis are publicly available at the GNPS-MassIVE repository under the MassIVE accession MSV000087027. The coordinates and structural factors have been deposited with the Protein Data Bank under accession codes **7EI3** (MasL) and **7EI4** (MasL-Collimonin C complex)

## Acknowledgments

We thank Dr. Chao-Jen Shih (Bioresource Collection and Research Center, Taiwan) for isolating and identifying *Massilia* sp. YMA4. The materials and methods for the construction of biosynthetic gene-null mutant strain were generously provided by Prof. Nai-Chun Lin’s Lab (National Taiwan University, Taiwan). We thank Prof. Ching-Hsuan Lin and Ms. Chih-Chieh Hsu (National Taiwan University, Taiwan) for providing clinical *Candida* strains and the material for the construction of tetracycline-inducible expression system. We thank Dr. Pei-Wen Hsiao (Agricultural Biotechnology Research Center, Academia Sinica, Taiwan) for providing human prostate PC-3 cell line for cloning ACAT1 gene. We thank Mr. Ning Lu, Ms. Ying-Mi Lai, and Ms. Chia-Chi Peng for collecting preliminary data. NMR data were collected in the High Field Nuclear Magnetic Resonance Center, Academia Sinica. LC-MS data were collected in the Metabolomics Core Facility, Agricultural Biotechnology Research Center, Academia Sinica, and the Proteomics Core Laboratory, Institute of Plant and Microbial Biology, Academia Sinica. TEM data were collected in the Biological Electron Microscopy Core Facility, Academia Sinica. The EM core facility is funded by the Academia Sinica Core Facility and Innovative Instrument Project (AS-CFII-108-119). We further thank the Protein Crystallization Facility (Academia Sinica Protein Clinic, Academia Sinica) for assistance in crystallization preparation; and the National Synchrotron Radiation Research Center (Hsinchu, Taiwan) with beamlines TLS 15A and TPS 05A in the for assistance in X-ray data collection and access to the synchrotron radiation centers.

## Author contributions

C.-C.L., S.Y.H., C.L., C.-H.S., H.-J.L., P.-Y.C., L.-J.S., B.-W.W., and W.-C.H. performed the experiments. C.-C.L., S.Y.H., K.-F.H., and Y.-N.H. carried out the data analysis. C.-C.L and S.Y.H. wrote the manuscript. Y.-L.Y. supervised the study. Y.-L.Y acquired funding to support the work.

## Additional information

**Supplementary information** The online version contains supplementary material available at xxxxxxx

The supplementary data descriptions as following:

**Supplementary Data 1:** RNA-seq analysis for different culture media by CLC workbench.

**Supplementary Data 2:** KEGG pathway analysis of DEG from RNA-seq analysis.

**Supplementary Data 3:** *In silico* prediction of biosynthetic gene clusters in *Massilia* sp. YMA4 by DeepBGC and antiSMASH.

**Supplementary Data 4:** MultiGeneBlast results of *mas* BGC query in BCT database.

**Supplementary Data 5:** Phylogenetic tree of bacterial polyyne biosynthetic gene clusters.

**Supplementary Data 6:** Bottom-up proteomics data of polyyne-modification proteins.

## References

1. Sanglard D. Emerging Threats in Antifungal-Resistant Fungal Pathogens. Frontiers in medicine 3, 11–11 (2016).

2. Sardi JCO, Scorzoni L, Bernardi T, Fusco-Almeida AM, Mendes Giannini MJS. Candida species: current epidemiology, pathogenicity, biofilm formation, natural antifungal products and new therapeutic options. Journal of medical microbiology 62, 10–24 (2013).

3. Pappas PG, et al. Clinical Practice Guideline for the Management of Candidiasis: 2016 Update by the Infectious Diseases Society of America. Clinical infectious diseases : an official publication of the Infectious Diseases Society of America 62, e1–50 (2016).

4. Lee Y, Puumala E, Robbins N, Cowen LE. Antifungal Drug Resistance: Molecular Mechanisms in Candida albicans and Beyond. Chemical Reviews 121, 3390–3411 (2021).

5. Yassin MT, Mostafa AA, Al-Askar AA, Bdeer R. In vitro antifungal resistance profile of Candida strains isolated from Saudi women suffering from vulvovaginitis. European Journal of Medical Research 25, 1 (2020).

6. Bhattacharya S, Sae-Tia S, Fries BC. Candidiasis and Mechanisms of Antifungal Resistance. Antibiotics (Basel) 9, 312 (2020).

7. Wall G, Lopez-Ribot JL. Current Antimycotics, New Prospects, and Future Approaches to Antifungal Therapy. Antibiotics (Basel) 9, 445 (2020).

8. Negri R. Polyacetylenes from terrestrial plants and fungi: Recent phytochemical and biological advances. Fitoterapia 106, 92–109 (2015).

9. Mullins AJ, et al. Genome mining identifies cepacin as a plant-protective metabolite of the biopesticidal bacterium Burkholderia ambifaria. Nat Microbiol 4, 996–1005 (2019).

10. W L Parker MLR, V Seiner, W H Trejo, P A Principe, R B Sykes. Cepacin A and cepacin B, two new antibiotics produced by Pseudomonas cepacia. J Antibiot (Tokyo) 37, 431–440 (1984).

11. Kai K, Sogame M, Sakurai F, Nasu N, Fujita M. Collimonins A–D, Unstable Polyynes with Antifungal or Pigmentation Activities from the Fungus-Feeding Bacterium Collimonas fungivorans Ter331. Organic Letters 20, 3536–3540 (2018).

12. Fritsche K, et al. Biosynthetic genes and activity spectrum of antifungal polyynes from Collimonas fungivorans Ter331. Environ Microbiol 16, 1334–1345 (2014).

13. M Patel MC, A Horan, D Loebenberg, J Marquez, R Mierzwa, M S Puar, R Yarborough, J A Waitz. Sch 31828, a novel antibiotic from a Microbispora sp. taxonomy, fermentation, isolation and biological properties. J Antibiot (Tokyo) 41, 794–797 (1988).

14. Chen PY, Lu N, Lai YM, Yang YL. Anti-microbial metabolites from a marine bacterium YMA4. Planta Med 82, P593 (2016).

15. Kanehisa M, Furumichi M, Tanabe M, Sato Y, Morishima K. KEGG: new perspectives on genomes, pathways, diseases and drugs. Nucleic Acids Res 45, D353–D361 (2017).

16. Blin K, Shaw S, Kautsar SA, Medema MH, Weber T. The antiSMASH database version 3: increased taxonomic coverage and new query features for modular enzymes. Nucleic Acids Res 49, D639–D643 (2021).

17. Hannigan GD, et al. A deep learning genome-mining strategy for biosynthetic gene cluster prediction. Nucleic Acids Research 47, e110–e110 (2019).

18. Medema MH, Takano E, Breitling R. Detecting sequence homology at the gene cluster level with MultiGeneBlast. Mol Biol Evol 30, 1218–1223 (2013).

19. Kusumi T, Ohtani I, Nishiyama K, Kakisawa H. Caryoynencins, potent antibiotics from a plant pathogen Pseudomonas caryophylli. Tetrahedron Letters 28, 3981–3984 (1987).

20. Ross C, Scherlach K, Kloss F, Hertweck C. The molecular basis of conjugated polyyne biosynthesis in phytopathogenic bacteria. Angew Chem Int Ed Engl 53, 7794–7798 (2014).

21. Murata K, Suenaga M, Kai K. Genome Mining Discovery of Protegenins A–D, Bacterial Polyynes Involved in the Antioomycete and Biocontrol Activities of Pseudomonas protegens. ACS Chemical Biology, doi.org/10.1021/acschembio.1c00276 (2021).

22. Ueoka R, et al. Genome-Based Identification of a Plant-Associated Marine Bacterium as a Rich Natural Product Source. Angewandte Chemie International Edition 57, 14519–14523 (2018).

23. A KR, Shah AH, Prasad R. MFS transporters of Candida species and their role in clinical drug resistance. FEMS Yeast Res 16, (2016).

24. Yan Y, Liu N, Tang Y. Recent developments in self-resistance gene directed natural product discovery. Natural Product Reports 37, 879–892 (2020).

25. Hobson C, Chan AN, Wright GD. The Antibiotic Resistome: A Guide for the Discovery of Natural Products as Antimicrobial Agents. Chem Rev 121, 3464–3494 (2021).

26. Chopra I. Over-expression of target genes as a mechanism of antibiotic resistance in bacteria. The Journal of antimicrobial chemotherapy 41, 584–588 (1998).

27. Sutanto F, Konstantinidou M, Domling A. Covalent inhibitors: a rational approach to drug discovery. RSC Med Chem 11, 876–884 (2020).

28. Fan J, et al. Tetrameric Acetyl-CoA Acetyltransferase 1 Is Important for Tumor Growth. Mol Cell 64, 859–874 (2016).

29. Ohnuma T, Nakayama S, Anan E, Nishiyama T, Ogura K, Hiratsuka A. Activation of the Nrf2/ARE pathway via S-alkylation of cysteine 151 in the chemopreventive agent-sensor Keap1 protein by falcarindiol, a conjugated diacetylene compound. Toxicol Appl Pharmacol 244, 27–36 (2010).

30. Modis Y, Wierenga RK. A biosynthetic thiolase in complex with a reaction intermediate: the crystal structure provides new insights into the catalytic mechanism. Structure 7, 1279–1290 (1999).

31. Kim S, et al. Redox-switch regulatory mechanism of thiolase from Clostridium acetobutylicum. Nat Commun 6, 8410 (2015).

32. Zhang Y, Wei W, Fan J, Jin C, Lu L, Fang W. Aspergillus fumigatus Mitochondrial Acetyl Coenzyme A Acetyltransferase as an Antifungal Target. Appl Environ Microbiol 86, (2020).

33. Haapalainen AM, Meriläinen G, Pirilä PL, Kondo N, Fukao T, Wierenga RK. Crystallographic and Kinetic Studies of Human Mitochondrial Acetoacetyl-CoA Thiolase: The Importance of Potassium and Chloride Ions for Its Structure and Function. Biochemistry 46, 4305–4321 (2007).

34. Modis Y, Wierenga RK. A biosynthetic thiolase in complex with a reaction intermediate: the crystal structure provides new insights into the catalytic mechanism. Structure 7, 1279–1290 (1999).

35. Lewis K. Platforms for antibiotic discovery. Nat Rev Drug Discov 12, 371–387 (2013).

36. Chen A, Re RN, Burkart MD. Type II fatty acid and polyketide synthases: deciphering protein-protein and protein-substrate interactions. Nat Prod Rep 35, 1029–1045 (2018).

37. Soucy SM, Huang J, Gogarten JP. Horizontal gene transfer: building the web of life. Nat Rev Genet 16, 472–482 (2015).

38. Palmer AC, Kishony R. Opposing effects of target overexpression reveal drug mechanisms. Nature Communications 5, 4296 (2014).

39. Sugden CJ, Roper JR, Williams JG. Engineered gene over-expression as a method of drug target identification. Biochemical and Biophysical Research Communications 334, 555–560 (2005).

40. Saraon P, et al. Evaluation and prognostic significance of ACAT1 as a marker of prostate cancer progression. Prostate 74, 372–380 (2014).

41. Jiang P, et al. In vitro and in vivo anticancer effects of mevalonate pathway modulation on human cancer cells. Br J Cancer 111, 1562–1571 (2014).

42. Bloxham DP, Chalkley RA, Coghlin SJ, Salam W. Synthesis of chloromethyl ketone derivatives of fatty acids. Their use as specific inhibitors of acetoacetyl-coenzyme A thiolase, cholesterol biosynthesis and fatty acid synthesis. Biochemical Journal 175, 999–1011 (1978).

43. Holland PC, Clark MG, Bloxham DP. Inactivation of pig heart thiolase by 3-butynoyl coenzyme A, 3-pentynoyl coenzyme A, and 4-bromocrotonyl coenzyme A. Biochemistry 12, 3309–3315 (1973).

44. Palmer MA, et al. Biosynthetic thiolase from Zoogloea ramigera. Evidence for a mechanism involving Cys-378 as the active site base. The Journal of biological chemistry 266, 8369–8375 (1991).

45. Kumar S, Stecher G, Li M, Knyaz C, Tamura K. MEGA X: Molecular Evolutionary Genetics Analysis across Computing Platforms. Molecular biology and evolution 35, 1547–1549 (2018).

46. Letunic I, Bork P. Interactive Tree Of Life (iTOL) v4: recent updates and new developments. Nucleic Acids Research 47, W256–W259 (2019).

47. Shevchenko A, Tomas H, Havlis J, Olsen JV, Mann M. In-gel digestion for mass spectrometric characterization of proteins and proteomes. Nat Protoc 1, 2856–2860 (2006).

48. Diament BJ, Noble WS. Faster SEQUEST searching for peptide identification from tandem mass spectra. J Proteome Res 10, 3871–3879 (2011).

49. Long T, Sun Y, Hassan A, Qi X, Li X. Structure of nevanimibe-bound tetrameric human ACAT1. Nature 581, 339–343 (2020).

50. Otwinowski Z, Minor W. Processing of X-ray diffraction data collected in oscillation mode. Methods in enzymology 276, 307–326 (1997).

51. Kantardjieff KA, Rupp B. Matthews coefficient probabilities: Improved estimates for unit cell contents of proteins, DNA, and protein–nucleic acid complex crystals. Protein Science 12, 1865–1871 (2003).

52. Vagin A, Teplyakov A. Molecular replacement with MOLREP. Acta crystallographica Section D, Biological crystallography 66, 22–25 (2010).

53. Cowtan K. The Buccaneer software for automated model building. 1. Tracing protein chains. Acta crystallographica Section D, Biological crystallography 62, 1002–1011 (2006).

54. Perrakis A, Morris R, Lamzin VS. Automated protein model building combined with iterative structure refinement. Nature structural biology 6, 458–463 (1999).

55. Emsley P, Cowtan K. Coot: model-building tools for molecular graphics. Acta crystallographica Section D, Biological crystallography 60, 2126–2132 (2004).

56. Murshudov GN, et al. REFMAC5 for the refinement of macromolecular crystal structures. Acta Crystallographica Section D 67, 355–367 (2011).

57. Williams CJ, et al. MolProbity: More and better reference data for improved all-atom structure validation. Protein science : a publication of the Protein Society 27, 293–315 (2018).

58. Lambert RJW, Pearson J. Susceptibility testing: accurate and reproducible minimum inhibitory concentration (MIC) and non-inhibitory concentration (NIC) values. Journal of Applied Microbiology 88, 784–790 (2000).

