## Supplementary Information for "Multi-omics approach to identify bacterial polyynes and unveil their antifungal mechanism against *Candida albicans*"

| Contents | Title | Page |
|---|---|---|
| **Supplementary Figures** | | |
| **Figure 1** | Transcriptomics analysis of *Massilia* sp. YMA4 growth in polyynes production (PDA) versus non-production (YMA) medium | **4** |
| **Figure 2** | *mas* gene null-mutant strain was inactive against *Candida* species | **5** |
| **Figure 3** | UPLC-DAD-HRMS/MS of dominant polyynes in *Massilia* sp. YMA4 and absent in *ΔmasH* strain | **6** |
| **Figure 4** | Tandem mass spectra and fragment annotation of polyynes | **7** |
| **Figure 5** | Minimum inhibitory concentration (MIC) of polyynes and commercial drugs against *C. albicans* ATCC18804 in YPD medium | **8** |
| **Figure 6** | Bottom-up proteomics analysis of MasL treated by polyynes | **9** |
| **Figure 7** | Residual enzyme activity of *C. albicans* ERG10 and human ACAT1 treated by polyynes | **10** |
| **Figure 8** | Bottom-up proteomics analysis of *C. albicans* ERG10 treated by polyynes | **11** |
| **Figure 9** | Bottom-up proteomics analysis of human ACAT1 treated by polyynes | **13** |
| **Figure 10** | Overall structures of MasL and MasL-collimonin C complex | **15** |
| **Figure 11** | Monomeric structure of MasL covalently modified by collimonin C | **16** |
| **Figure 12** | Sequence alignment of acetyl-CoA acetyltransferases from different organisms | **17** |
| **Figure 13** | Electron density map of collimonin C in MasL reactive pocket | **18** |
| **Figure 14** | Magnification view of MasL covalently modified by collimonin C | **19** |
| **Figure 15** | Superimposition of MasL and MasL-collimonin C complex | **20** |
| **Figure 16** | Reported inhibitors of acetyl-CoA acetyltransferase (EC 2.3.1.9) | **21** |
| **Figure 17** | Construction scheme (a) and PCR check result (b) of the null- mutant strain *ΔmasH* | **22** |
| **Figure 18** | Tetracycline-inducible expression system in *C. albicans* ATCC18804 | **23** |
| **Figure 19** | $^1$H NMR and $^1$H-$^1$H COSY (600 MHz) of collimonin C **1** | **24** |
| **Figure 20** | HSQC and HMBC (600 MHz) of collimonin C **1** | **25** |
| **Figure 21** | $^1$H NMR and $^1$H-$^1$H COSY (600 MHz) of collimonin D **2** | **26** |
| **Figure 22** | HSQC and HMBC (600 MHz) of collimonin D **2** | **27** |
| **Figure 23** | $^1$H NMR and $^1$H-$^1$H COSY (600 MHz) of massilin A **3** | **28** |
| **Figure 24** | HSQC and HMBC (600 MHz) of massilin A **3** | **29** |
| **Figure 25** | $^1$H NMR and $^1$H-$^1$H COSY (600 MHz) of massilin B **4** | **30** |
| **Figure 26** | HSQC and HMBC (600 MHz) of massilin B **4** | **31** |
| **Supplementary Tables** | | |
| **Table 1** | Characterization of *mas* biosynthetic gene cluster | **32** |
| **Table 2** | Characterization of core genes in cepacin biosynthetic gene cluster | **34** |

| | | |
|---|---|---|
| **Table 3** | List of primer sets used in this study | **35** |
| **Table 4** | List of strains used in this study | **36** |
| **Table 5** | List of plasmids used in this study | **37** |
| **Table 6** | Whole-genome sequences used in mining bacterial polyyne gene clusters | **38** |
| **Table 7** | 1D NMR of collimonin C **1** in $C_2D_6OS$ | **40** |
| **Table 8** | 1D NMR of collimonin D **2** in $C_2D_6OS$ | **41** |
| **Table 9** | 1D NMR of massilin A **3** in $C_2D_6OS$ | **42** |
| **Table 10** | 1D NMR of massilin B **4** in $C_2D_6OS$ | **43** |
| **Supplementary Methods** | | |
| **Method 1** | Chemicals, strains, plasmids, and culture conditions | **44** |
| **Method 2** | Antagonism assay | **45** |
| **Method 3** | Quantitative RT-PCR | **45** |
| **Method 4** | Construction of polyyne biosynthesis gene-null mutant strain *ΔmasH* | **45** |
| **Method 5** | Construction of inducible *ERG10* overexpression strains and *masL*$_{opt}$ heterologous expression strain in *C. albicans* | **46** |
| **Method 6** | Construction of inducible *masL* and *ACAT1* heterologous expression strains in *E. coli* | **46** |
| **Method 7** | Expression and purification of MasL, ERG10, and ACAT1 | **46** |
| **Method 8** | Genome sequence and annotation | **47** |
| **Method 9** | RNA sequencing and transcriptomic analysis | **48** |
| **Method 10** | Extraction of *Massilia* sp. YMA4 and UPLC-DAD-MS/MS analysis | **49** |
| **Method 11** | Isolation, structure elucidation, and quantitation of compounds 1-4 | **50** |
| **Method 12** | Sequence alignment and structure superimposition | **52** |
| **Method 13** | Transmission Electron Microscope | **52** |
| **Supplementary Notes** | | |
| **Note 1** | Background of kinetic evaluation of irreversible inhibitors and polyynes-MasL experiment detail. | **53** |
| **Supplementary References** | | |

### Supplementary Figures

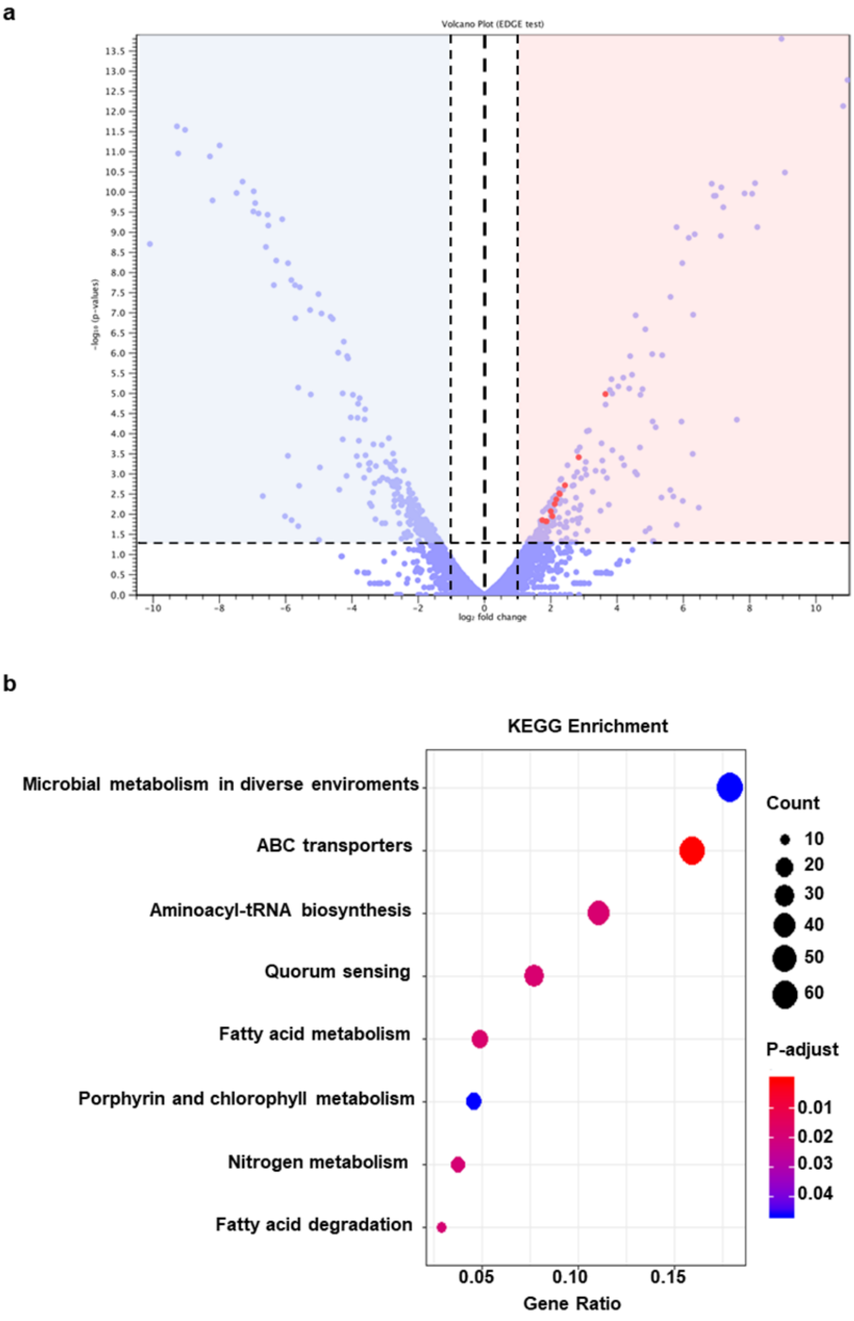


**Supplementary Figure 1 |** Transcriptomics analysis of *Massilia* sp. YMA4 growth in polyynes production (PDA) versus non-production (YMA) medium. (**a**) Volcano plot of differentially expressed genes (DEGs) mapping to 192 upregulated genes (red area) and 226 downregulated genes (blue area). Dot-line represent as statistical criteria for p-value < 0.05 and | fold change (FC) | > 2. Red-colored points indicate the polyyne BGC (*masA* to *masL*). (**b**) KEGG pathway analysis of DEGs. The significant pathway (p-adjust <0.05) shows in the panel.

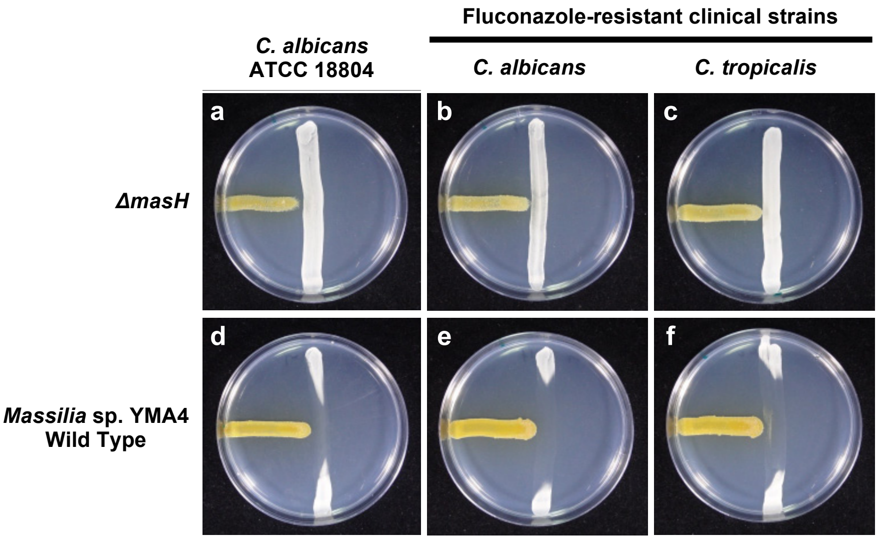


**Supplementary Figure 2 |** *mas* gene null-mutant strain was inactive against *Candida* species. (**a–c**) Antagonistic assay of *ΔmasH* strain against type strain *C. albicans* ATCC 18804 and clinically isolated fluconazole-resistant *C. albicans* and *C. tropicalis*. (**d-f**) Wild type *Massilia* sp. YMA4 was used as a positive control.

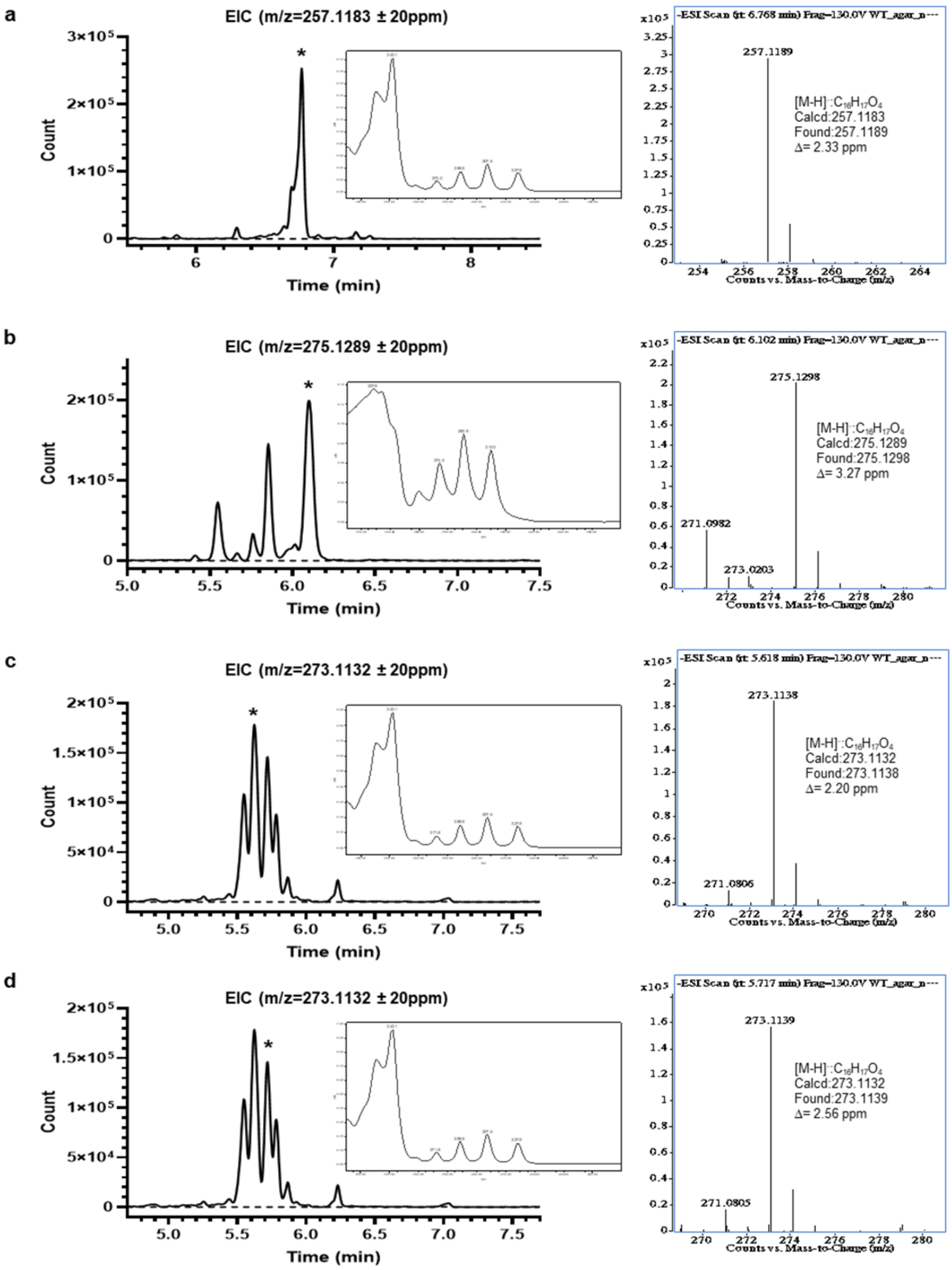


**Supplementary Figure 3 |** UPLC-DAD-HRMS/MS of dominant polyynes in *Massilia* sp. YMA4 and absent in *ΔmasH* strain. (**a**) Massilin A, (**b**) massilin B, (**c**) collimonin C (**d**), and collimonin D. Solid-line represent extracted-ion chromatography (EIC) of wild-type *Massilia* sp. YMA4 and dot-line represent EIC of *ΔmasH* strain. Asterisks in EIC indicate the peak of the polyynes' mass and UV spectra shown in the right panel.

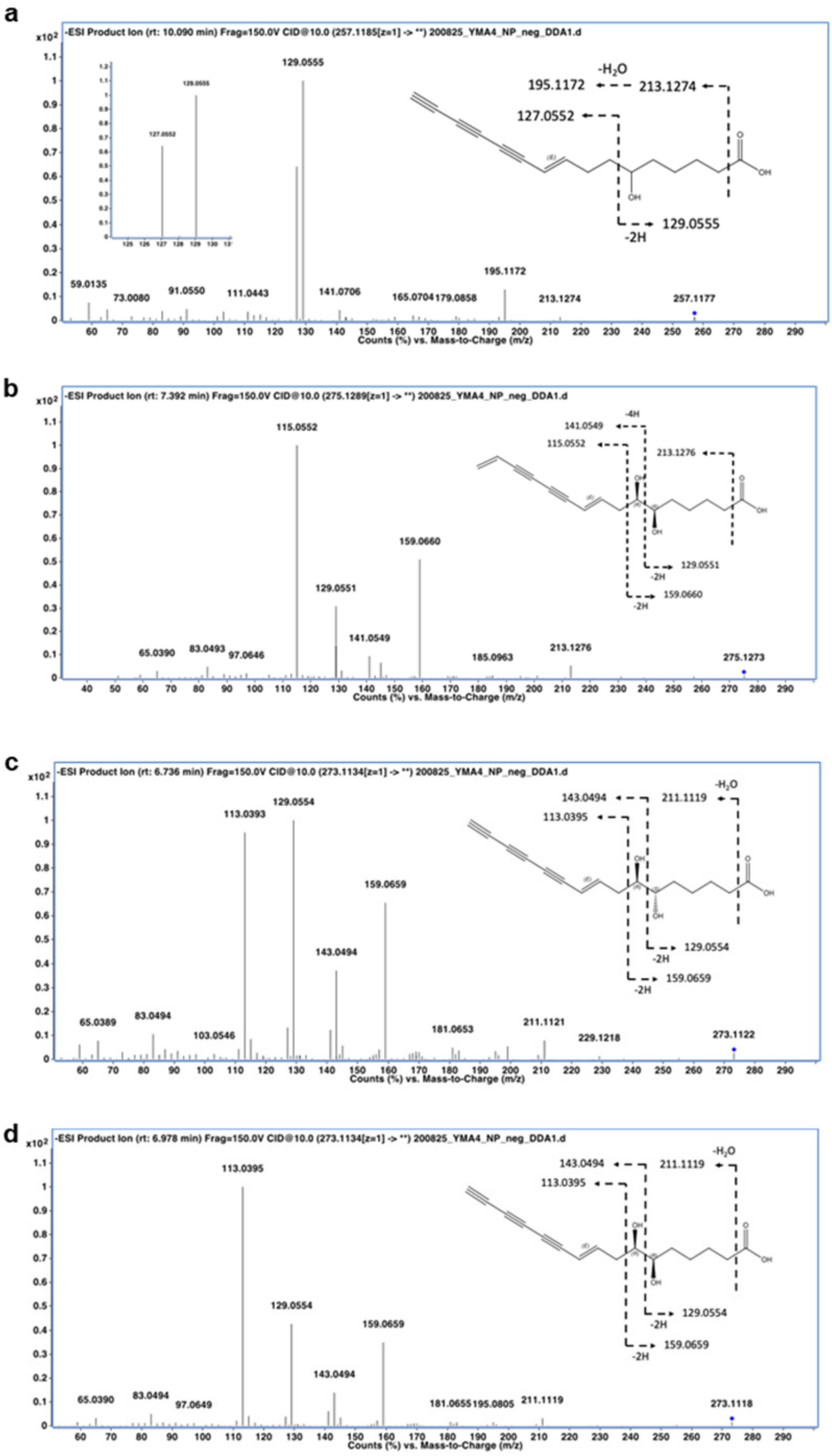


**Supplementary Figure 4 |** Tandem mass spectra and fragment annotation of polyynes. LC-HRMS/MS spectra and fragment annotation of (**a**) massilin A, (**b**) massilin B, (**c**) collimonin C (**d**), and collimonin D.

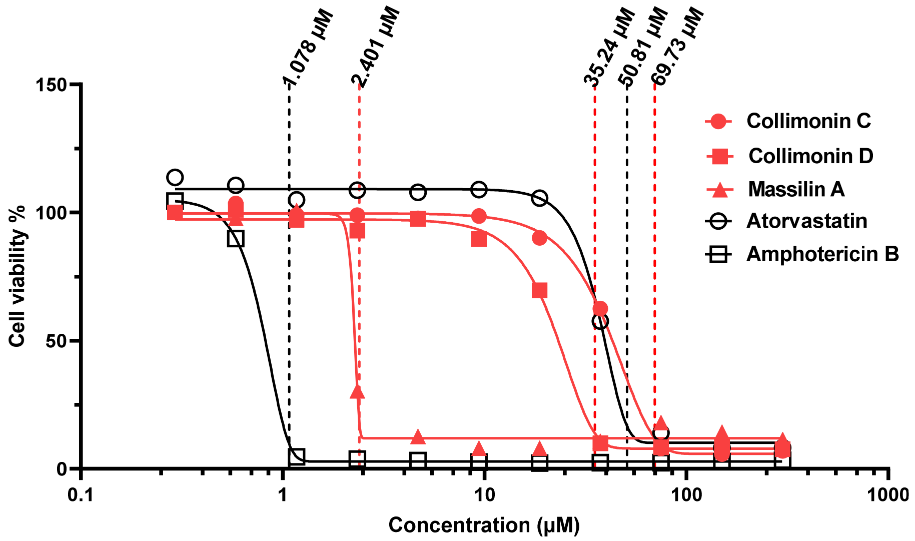


**Supplementary Figure 5 |** Minimum inhibitory concentration (MIC) of polyynes and commercial drugs against *C. albicans* ATCC18804 in YPD medium. Dose-response curves for cell viability treated by collimonin C (red filled circle), collimonin D (red filled square), massilin A (red filled triangle), and commercial drugs atorvastatin (empty black circle) and amphotericin B (empty black square). The MIC value of each compound indicates above the dot-line. Each point was obtained from three biological replicates.

**a**

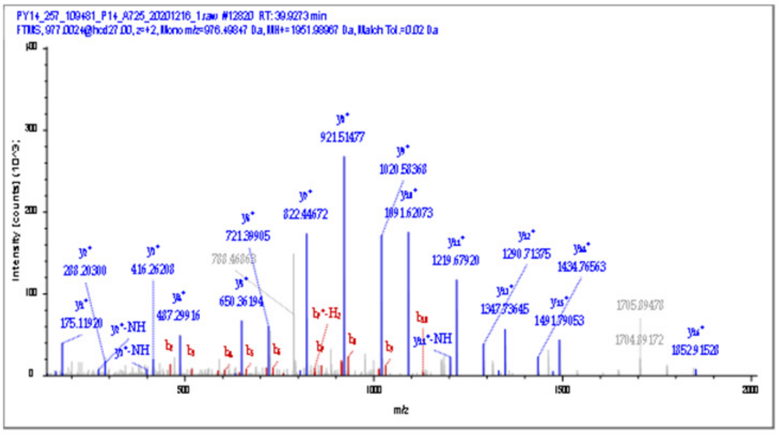


| #b | b⁺ | Seq. | y⁺ | #y |
|---|---|---|---|---|
| 1 | | V | | 17 |
| 2 | 1.70 | C* | 5.89 | 16 |
| 3 | -0.58 | G | 0.59 | 15 |
| 4 | -1.13 | S | 3.01 | 14 |
| 5 | | G | 1.09 | 13 |
| 6 | -2.37 | A | 2.10 | 12 |
| 7 | 0.87 | Q | 0.12 | 11 |
| 8 | 0.81 | A | 0.04 | 10 |
| 9 | -1.58 | V | -0.02 | 9 |
| 10 | -4.74 | V | 0.51 | 8 |
| 11 | | T | 0.13 | 7 |
| 12 | | A | 0.14 | 6 |
| 13 | | Y | 0.15 | 5 |
| 14 | | A | -0.93 | 4 |
| 15 | | Q | -1.17 | 3 |
| 16 | | I | 0.06 | 2 |
| 17 | | R | 4.29 | 1 |

**b**

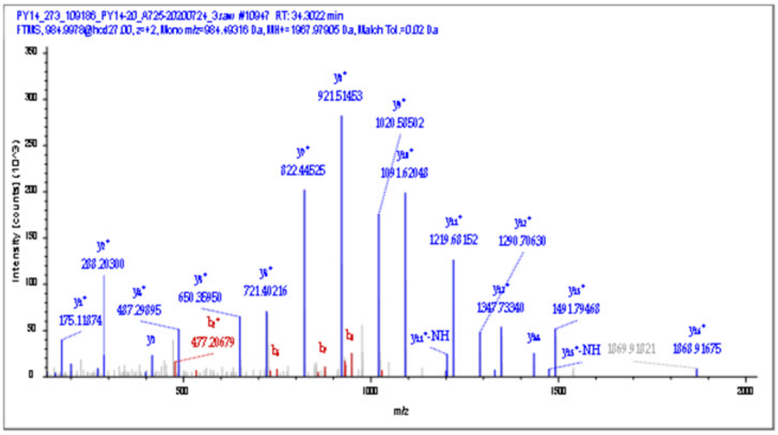


| #b | b⁺ | Seq. | y⁺ | #y |
|---|---|---|---|---|
| 1 | | V | | 17 |
| 2 | -2.97 | C* | 2.33 | 16 |
| 3 | 2.00 | G | -2.19 | 15 |
| 4 | | S | 2.68 | 14 |
| 5 | | G | 3.36 | 13 |
| 6 | -2.69 | A | 7.87 | 12 |
| 7 | | Q | -1.78 | 11 |
| 8 | 5.78 | A | 0.27 | 10 |
| 9 | | V | -1.34 | 9 |
| 10 | | V | 0.77 | 8 |
| 11 | | T | 1.92 | 7 |
| 12 | | A | -4.17 | 6 |
| 13 | | Y | 3.90 | 5 |
| 14 | | A | -0.50 | 4 |
| 15 | | Q | 5.49 | 3 |
| 16 | | I | 0.06 | 2 |
| 17 | | R | 1.21 | 1 |

**Supplementary Figure 6 |** Bottom-up proteomics analysis of MasL treated by polyynes. NanoLC-Q-HCD-orbitrap tandem mass spectra and annotation of massilin A (**a**) and collimonin C/D (**b**) derived covalent modification of trypsin-digested MasL peptides. The annotated ion peaks are colored in blue (y ion) and red (b ion). Mass errors are shown in ppm in the annotation table. Asterisks in the sequence indicate the modified cysteine residue.

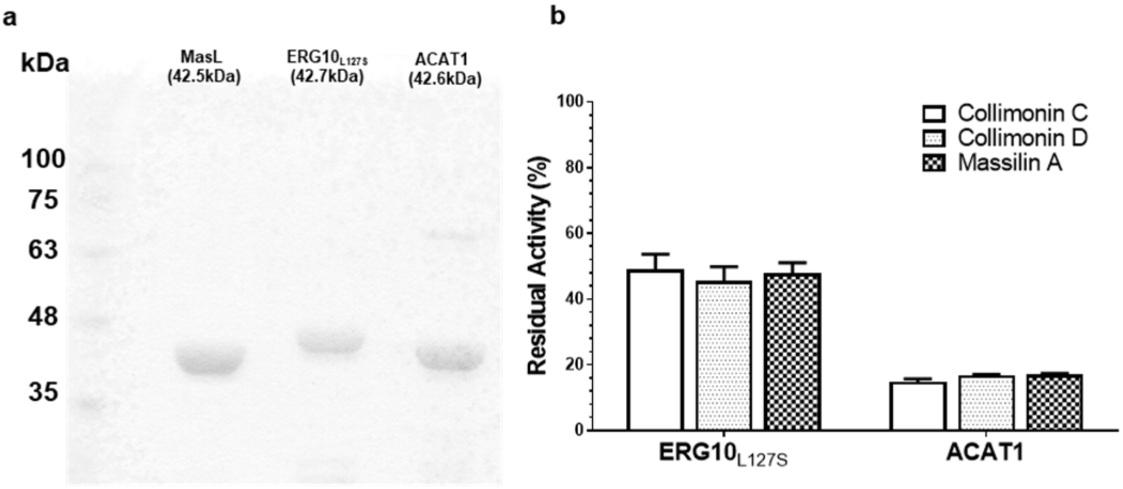


**Supplementary Figure 7 |** Residual enzyme activity of *C. albicans* ERG10 and human ACAT1 treated by polyynes. (**a**) SDS-PAGE of recombinant acetyl-CoA acetyltransferases (MasL from *Massilia* sp. YMA4; $ERG10_{L127S}$ from *C. albicans* ATCC18804; ACAT1 (transition peptide truncated, aa 34-427) from human). (**b**) Residual enzyme activity of $ERG10_{L127S}$ and ACAT1 treated by polyynes. Reaction mixtures contained 20 μM enzyme and 100 μM polyyne for 2 hr reaction and followed by 200 μM acetyl-CoA for 1 hr substrate reaction. Residual activity was detected by the CPM labeling method and normalized with 2% DMSO as a control treatment. Each enzyme-ligand pair with replicates, n=3-4. Error bar shows the standard error of the mean (SEM).

**a**

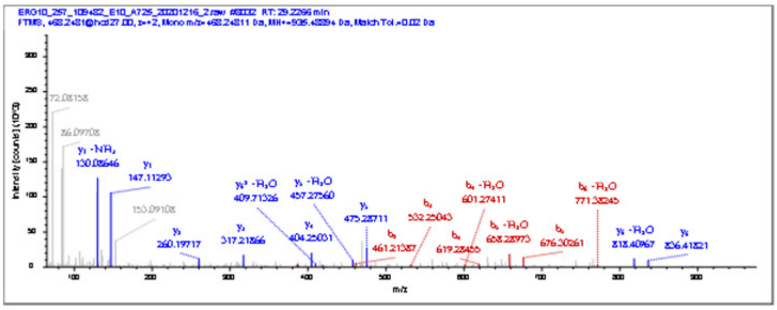

| #b | $b^+$ | Seq. | $y^+$ | #y |
|---|---|---|---|---|
| 1 | | V | | 7 |
| 2 | -7.36 | C* | 4.84 | 6 |
| 3 | -5.34 | A | 0.77 | 5 |
| 4 | -7.97 | S | 0.12 | 4 |
| 5 | -2.26 | G | -1.03 | 3 |
| 6 | | L | -1.16 | 2 |
| 7 | | K | -0.85 | 1 |

**b**

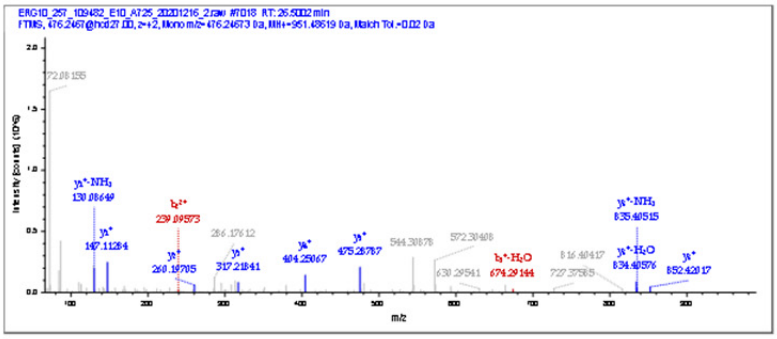

| #b | $b^+$ | Seq. | $y^+$ | #y |
|---|---|---|---|---|
| 1 | | V | | 7 |
| 2 | | C* | -3.53 | 6 |
| 3 | | A | -0.83 | 5 |
| 4 | | S | -0.77 | 4 |
| 5 | | G | -0.25 | 3 |
| 6 | | L | -0.70 | 2 |
| 7 | | K | -0.24 | 1 |

**c**

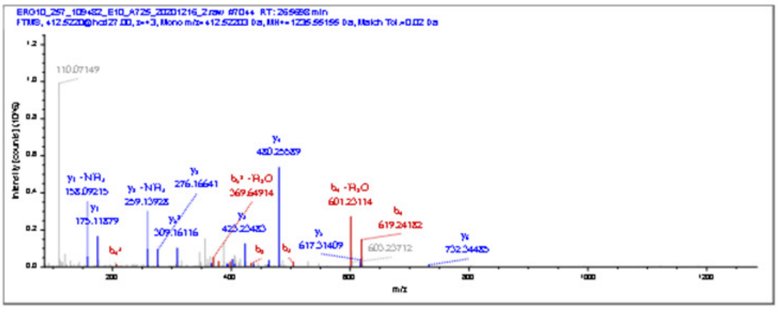

| #b | $b^+$ | Seq. | $y^+$ | #y |
|---|---|---|---|---|
| 1 | | C* | | 9 |
| 2 | -2.09 | A | | 8 |
| 3 | -0.04 | A | 0.56 | 7 |
| 4 | 2.28 | D | -3.40 | 6 |
| 5 | | H | 2.15 | 5 |
| 6 | | G | 1.29 | 4 |
| 7 | | F | 0.51 | 3 |
| 8 | | T | 0.8 | 2 |
| 9 | | R | 0.93 | 1 |

**d**

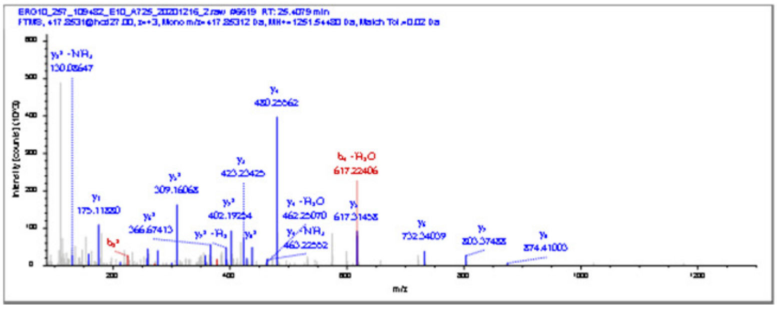

| #b | $b^+$ | Seq. | $y^+$ | #y |
|---|---|---|---|---|
| 1 | | C* | | 9 |
| 2 | | A | 7.5 | 8 |
| 3 | | A | 5.72 | 7 |
| 4 | | D | 2.69 | 6 |
| 5 | | H | 1.36 | 5 |
| 6 | | G | 1.85 | 4 |
| 7 | | F | 1.88 | 3 |
| 8 | | T | -1.99 | 2 |
| 9 | | R | 0.87 | 1 |

**Supplementary Figure 8 |** Bottom-up proteomics analysis of *C. albicans* ERG10 treated by polyynes. NanoLC-Q-HCD-orbitrap tandem mass spectra and annotation of massilin A (**a, c**) and collimonin C/D (**b, d**) derived covalent modification of trypsin-digested ERG10 peptides (Cys90 contained (**a, b**) and Cys166 contained (**c, d**)). The annotated ion peaks are colored in blue (y ion) and red (b ion). Mass errors are shown in ppm in the annotation table. Asterisks in the sequence indicate the modified cysteine residue.

**e**

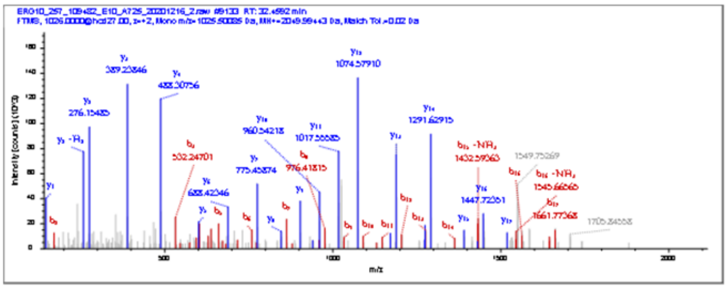

| #b | b⁺ | Seq. | y⁺ | #y |
|---|---|---|---|---|
| 1 | | I | | 20 |
| 2 | 0.08 | G | | 19 |
| 3 | 1.09 | C* | | 18 |
| 4 | | A | 6.19 | 17 |
| 5 | 0.86 | G | -1.77 | 16 |
| 6 | 7.01 | V | 8.36 | 15 |
| 7 | 1.32 | C | 1.49 | 14 |
| 8 | 8.75 | N | 0.26 | 13 |
| 9 | 7.95 | G | -0.13 | 12 |
| 10 | 6.95 | G | 1.62 | 11 |
| 11 | -0.33 | G | -6.40 | 10 |
| 12 | -1.85 | G | -2.24 | 9 |
| 13 | -1.74 | A | 7.08 | 8 |
| 14 | -1.87 | S | -3.54 | 7 |
| 15 | | S | 0.74 | 6 |
| 16 | 6.43 | I | -5.22 | 5 |
| 17 | -4.50 | V | 0.65 | 4 |
| 18 | | I | 2.57 | 3 |
| 19 | | E | 1.98 | 2 |
| 20 | | K | 1.32 | 1 |

| #b | b⁺ | Seq. | y⁺ | #y |
|---|---|---|---|---|
| 1 | | I | | 20 |
| 2 | 0.08 | G | | 19 |
| 3 | 1.93 | C | | 18 |
| 4 | 1.89 | A | | 17 |
| 5 | 5.36 | G | 0.57 | 16 |
| 6 | 1.68 | V | | 15 |
| 7 | 1.32 | C* | 2.57 | 14 |
| 8 | 8.75 | N | 0.26 | 13 |
| 9 | 7.95 | G | -0.13 | 12 |
| 10 | 6.95 | G | 1.62 | 11 |
| 11 | -0.33 | G | -6.40 | 10 |
| 12 | -1.85 | G | -2.24 | 9 |
| 13 | -1.74 | A | 7.08 | 8 |
| 14 | -1.87 | S | -3.54 | 7 |
| 15 | | S | 0.74 | 6 |
| 16 | 6.43 | I | -5.22 | 5 |
| 17 | -4.50 | V | 0.65 | 4 |
| 18 | | I | 2.57 | 3 |
| 19 | | E | 1.98 | 2 |
| 20 | | K | 1.32 | 1 |

**f**

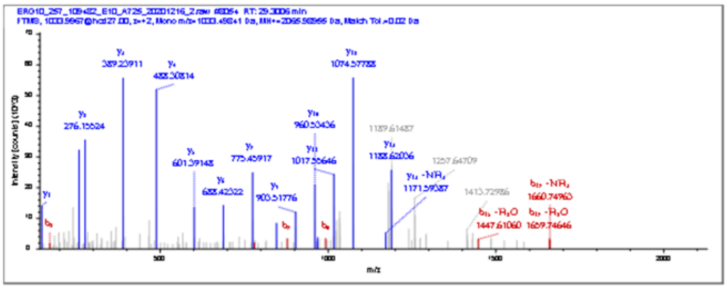

| #b | b⁺ | Seq. | y⁺ | #y |
|---|---|---|---|---|
| 1 | | I | | 20 |
| 2 | -1.26 | G | | 19 |
| 3 | | C* | | 18 |
| 4 | | A | | 17 |
| 5 | | G | | 16 |
| 6 | | V | | 15 |
| 7 | | C | | 14 |
| 8 | 3.90 | N | 1.29 | 13 |
| 9 | | G | 1.01 | 12 |
| 10 | | G | 1.02 | 11 |
| 11 | | G | 1.75 | 10 |
| 12 | | G | -3.53 | 9 |
| 13 | | A | 5.20 | 8 |
| 14 | | S | -4.09 | 7 |
| 15 | | S | 1.09 | 6 |
| 16 | | I | 0.76 | 5 |
| 17 | | V | -0.54 | 4 |
| 18 | | I | 0.90 | 3 |
| 19 | | E | 0.57 | 2 |
| 20 | | K | 0.91 | 1 |

| #b | b⁺ | Seq. | y⁺ | #y |
|---|---|---|---|---|
| 1 | | I | | 20 |
| 2 | -1.26 | G | | 19 |
| 3 | | C | | 18 |
| 4 | | A | | 17 |
| 5 | | G | | 16 |
| 6 | 1.86 | V | | 15 |
| 7 | | C* | | 14 |
| 8 | 3.90 | N | 1.29 | 13 |
| 9 | | G | 1.01 | 12 |
| 10 | | G | 1.02 | 11 |
| 11 | | G | 1.75 | 10 |
| 12 | | G | -3.53 | 9 |
| 13 | | A | 5.20 | 8 |
| 14 | | S | -4.09 | 7 |
| 15 | | S | 1.09 | 6 |
| 16 | | I | 0.76 | 5 |
| 17 | | V | -0.54 | 4 |
| 18 | | I | 0.90 | 3 |
| 19 | | E | 0.57 | 2 |
| 20 | | K | 0.91 | 1 |

**Supplementary Figure 8 (continued) |** Bottom-up proteomics analysis of *C. albicans* ERG10 treated by polyynes. NanoLC-Q-HCD-orbitrap tandem mass spectra and annotation of massilin A (**e**) and collimonin C/D (**f**) derived covalent modification of trypsin-digested ERG10 peptides (Cys378 or Cys382 contained, ambiguous). The annotated ion peaks are colored in blue (y ion) and red (b ion). Mass errors are shown in ppm in the annotation table. Asterisks in the sequence indicate the modified cysteine residue.

**a**

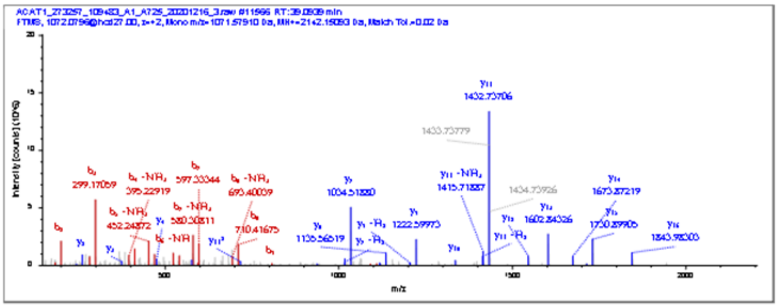

| #b | b⁺ | Seq. | y⁺ | #y |
|---|---|---|---|---|
| 1 | | Q | | 19 |
| 2 | 2.59 | A | | 18 |
| 3 | 2.65 | V | | 17 |
| 4 | 2.10 | L | 2.37 | 16 |
| 5 | 1.09 | G | 2.48 | 15 |
| 6 | 2.60 | A | 5.79 | 14 |
| 7 | 3.43 | G | 0.94 | 13 |
| 8 | 3.94 | L | 4.94 | 12 |
| 9 | 0.79 | P | 1.52 | 11 |
| 10 | | I | -1.05 | 10 |
| 11 | | S | 2.19 | 9 |
| 12 | 7.80 | T | 4.57 | 8 |
| 13 | | P | 3.77 | 7 |
| 14 | | C* | | 6 |
| 15 | | T | 0.65 | 5 |
| 16 | | T | 1.61 | 4 |
| 17 | | I | 3.97 | 3 |
| 18 | | N | 1.54 | 2 |
| 19 | | K | 1.86 | 1 |

**b**

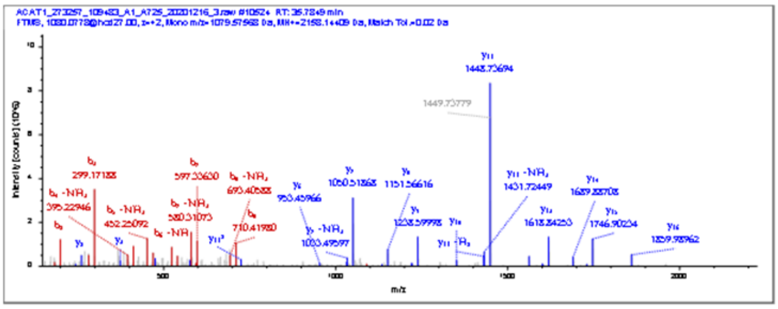

| #b | b⁺ | Seq. | y⁺ | #y |
|---|---|---|---|---|
| 1 | | Q | | 19 |
| 2 | -1.81 | A | | 18 |
| 3 | -1.67 | V | 1.90 | 17 |
| 4 | -2.27 | L | -3.93 | 16 |
| 5 | -1.96 | G | -2.35 | 15 |
| 6 | -1.92 | A | -6.10 | 14 |
| 7 | -1.36 | G | -1.77 | 13 |
| 8 | -0.35 | L | -6.75 | 12 |
| 9 | -1.78 | P | -1.94 | 11 |
| 10 | 6.02 | I | 6.22 | 10 |
| 11 | | S | -2.16 | 9 |
| 12 | | T | -0.77 | 8 |
| 13 | | P | -1.03 | 7 |
| 14 | | C* | 5.43 | 6 |
| 15 | | T | -0.62 | 5 |
| 16 | | T | -2.45 | 4 |
| 17 | | I | -0.44 | 3 |
| 18 | | N | -2.21 | 2 |
| 19 | | K | | 1 |

**c**

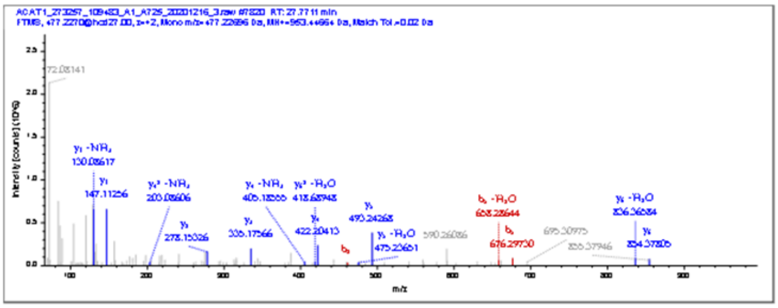

| #b | b⁺ | Seq. | y⁺ | #y |
|---|---|---|---|---|
| 1 | | V | | 7 |
| 2 | -4.91 | C* | 0.74 | 6 |
| 3 | | A | 2.46 | 5 |
| 4 | | S | 6.28 | 4 |
| 5 | 5.59 | G | -2.71 | 3 |
| 6 | | M | 0.10 | 2 |
| 7 | | K | 1.66 | 1 |

**d**

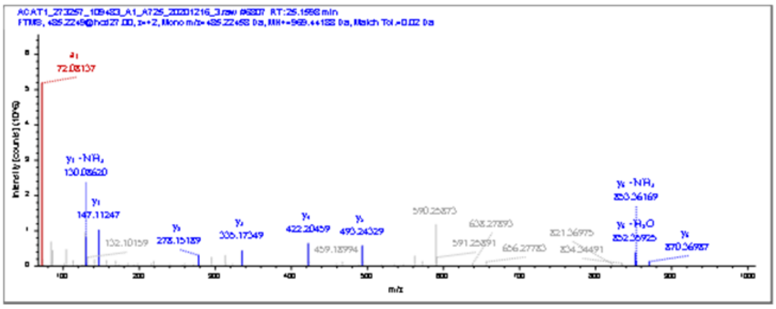

| #b | b⁺ | Seq. | y⁺ | #y |
|---|---|---|---|---|
| 1 | | V | | 7 |
| 2 | | C* | 4.27 | 6 |
| 3 | | A | 1.23 | 5 |
| 4 | | S | 5.19 | 4 |
| 5 | | G | 3.77 | 3 |
| 6 | | M | 5.03 | 2 |
| 7 | | K | 2.27 | 1 |

**Supplementary Figure 9 |** Bottom-up proteomics analysis of human ACAT1 treated by polyynes. NanoLC-Q-HCD-orbitrap tandem mass spectra and annotation of massilin A (**a, c**) and collimonin C/D (**b, d**) derived covalent modification of trypsin-digested ACAT1 peptides (Cys119 contained (**a, b**) and Cys126 contained (**c, d**)). The annotated ion peaks are colored in blue (y ion) and red (b ion). Mass errors are shown in ppm in the annotation table. Asterisks in the sequence indicate the modified cysteine residue.

e

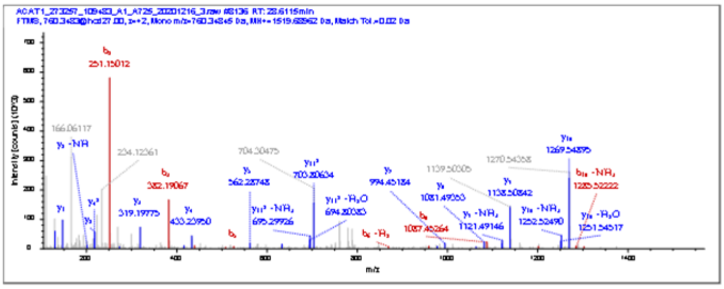

| #b | $b^+$ | Seq. | $y^+$ | #y |
|---|---|---|---|---|
| 1 | | I | | 12 |
| 2 | 0.53 | H | | 11 |
| 3 | 0.18 | M | 0.03 | 10 |
| 4 | 0.30 | G | 0.08 | 9 |
| 5 | -9.81 | S | -6.00 | 8 |
| 6 | | C* | 3.19 | 7 |
| 7 | -0.96 | A | 9.51 | 6 |
| 8 | 5.59 | E | -7.76 | 5 |
| 9 | 6.96 | N | 2.36 | 4 |
| 10 | 0.29 | T | -0.48 | 3 |
| 11 | | A | -0.83 | 2 |
| 12 | | K | -0.24 | 1 |

f

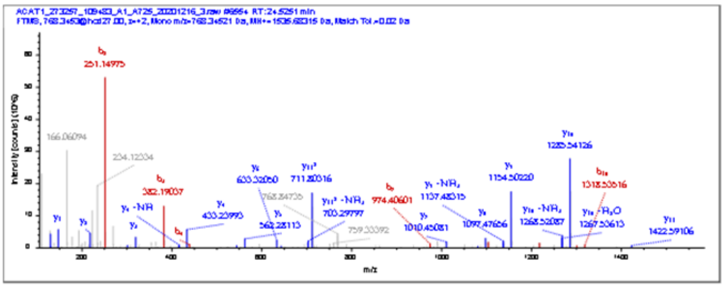

| #b | $b^+$ | Seq. | $y^+$ | #y |
|---|---|---|---|---|
| 1 | | I | | 12 |
| 2 | 2.00 | H | 8.25 | 11 |
| 3 | 0.96 | M | 2.05 | 10 |
| 4 | 0.37 | G | 1.05 | 9 |
| 5 | -0.65 | S | 4.91 | 8 |
| 6 | | C* | -0.89 | 7 |
| 7 | 5.15 | A | -0.43 | 6 |
| 8 | 2.11 | E | 3.53 | 5 |
| 9 | | N | 1.37 | 4 |
| 10 | 6.88 | T | 3.81 | 3 |
| 11 | 2.66 | A | 1.96 | 2 |
| 12 | | K | 1.52 | 1 |

g

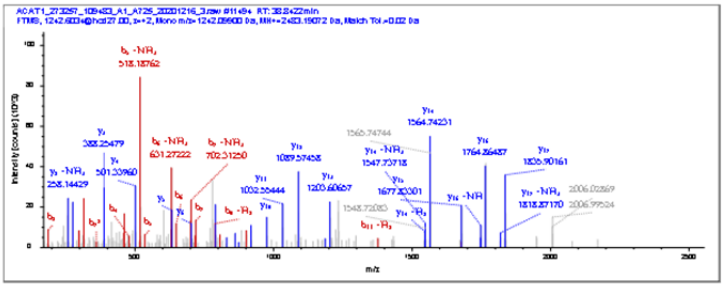

| #b | $b^+$ | Seq. | $y^+$ | #y |
|---|---|---|---|---|
| 1 | | Q | | 23 |
| 2 | -1.03 | G | | 22 |
| 3 | 0.19 | E | | 21 |
| 4 | -0.06 | Y | | 20 |
| 5 | -0.03 | G | | 19 |
| 6 | | L | | 18 |
| 7 | 5.01 | A | 0.77 | 17 |
| 8 | 8.08 | S | 0.59 | 16 |
| 9 | | I | 0.52 | 15 |
| 10 | | C* | 4.80 | 14 |
| 11 | | N | 7.03 | 13 |
| 12 | | G | -2.27 | 12 |
| 13 | | G | -3.68 | 11 |
| 14 | | G | | 10 |
| 15 | | G | 2.36 | 9 |
| 16 | | A | 3.95 | 8 |
| 17 | | S | -5.43 | 7 |
| 18 | | A | -1.40 | 6 |
| 19 | | M | 2.11 | 5 |
| 20 | | L | -0.18 | 4 |
| 21 | | I | 1.69 | 3 |
| 22 | | Q | 1.86 | 2 |
| 23 | | K | | 1 |

h

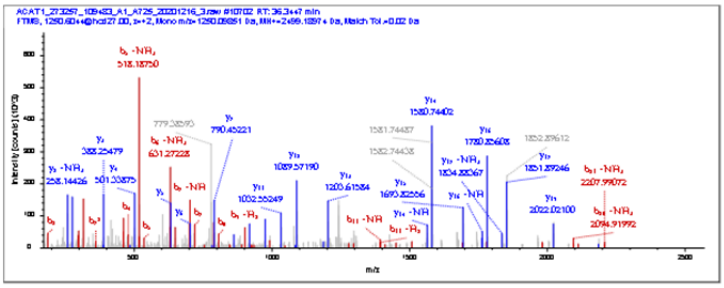

| #b | $b^+$ | Seq. | $y^+$ | #y |
|---|---|---|---|---|
| 1 | | Q | | 23 |
| 2 | 0.26 | G | | 22 |
| 3 | 0.95 | E | | 21 |
| 4 | 1.09 | Y | 1.4 | 20 |
| 5 | 4.08 | G | -8.68 | 19 |
| 6 | 3.67 | L | | 18 |
| 7 | 1.36 | A | 2.95 | 17 |
| 8 | 2.48 | S | 2.66 | 16 |
| 9 | | I | 1.9 | 15 |
| 10 | -5.37 | C* | 0.44 | 14 |
| 11 | -1.77 | N | -0.67 | 13 |
| 12 | | G | 0.19 | 12 |
| 13 | | G | -1.79 | 11 |
| 14 | | G | -1 | 10 |
| 15 | | G | -2.97 | 9 |
| 16 | | A | 4.24 | 8 |
| 17 | | S | -3.89 | 7 |
| 18 | | A | -0.87 | 6 |
| 19 | | M | 1.43 | 5 |
| 20 | -4.59 | L | 1.52 | 4 |
| 21 | | I | 1.69 | 3 |
| 22 | | Q | 1.21 | 2 |
| 23 | | K | | 1 |

**Supplementary Figure 9 (continued) |** Bottom-up proteomics analysis of human ACAT1 treated by polyynes. NanoLC-Q-HCD-orbitrap tandem mass spectra and annotation table of massilin A (**e, g**) and collimonin C/D (**f, h**) derived covalent modification of trypsin-digested ACAT1 peptides (Cys196 contained (**e, f**) and Cys413 contained (**g, h**)). The annotated ion peaks are colored in blue (y ion) and red (b ion). Mass errors are shown in ppm in the annotation table. Asterisks in the sequence indicate the modified cysteine residue.

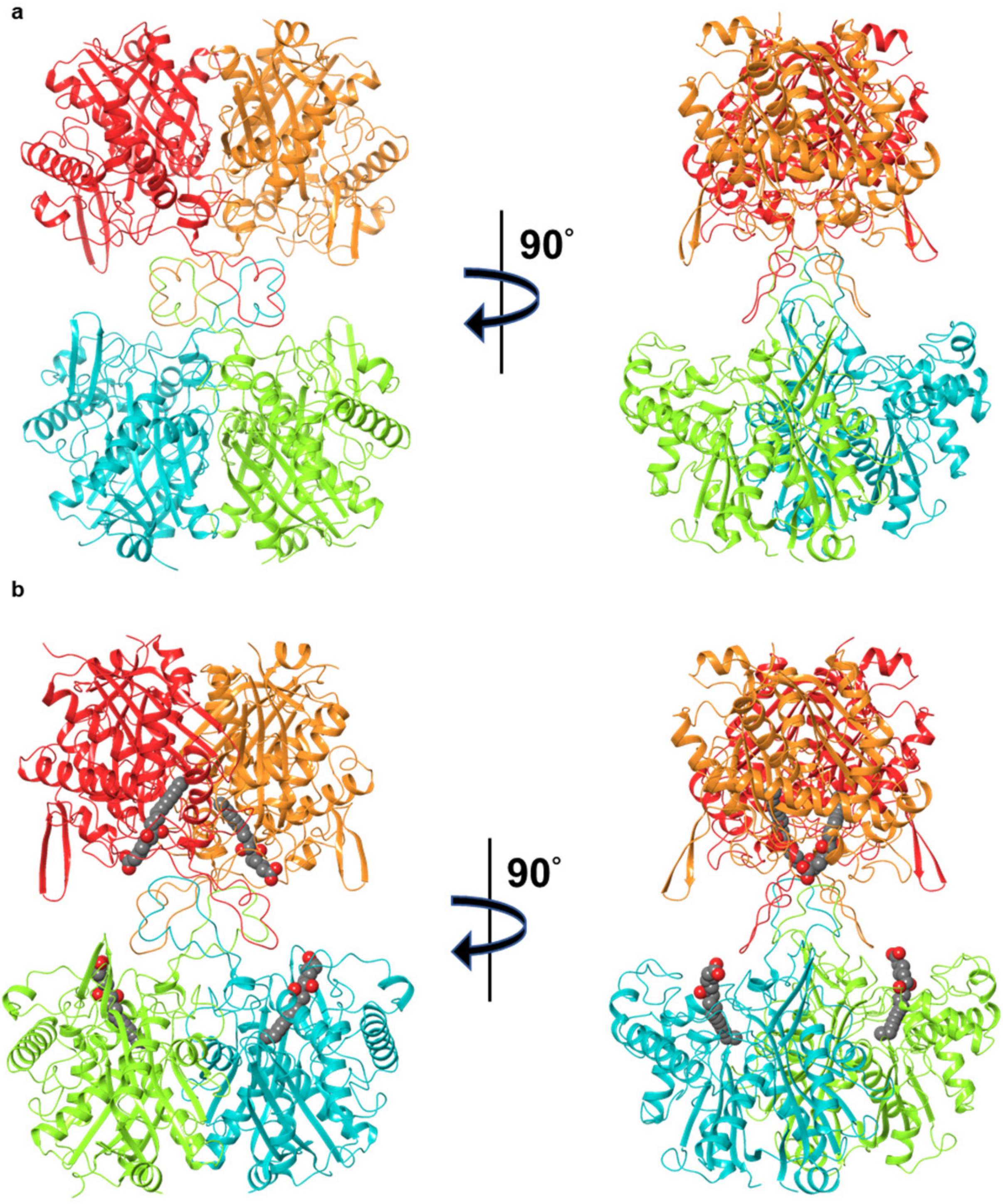


**Supplementary Figure 10 |** Overall structures of MasL and MasL-collimonin C complex. Tetrameric structures are shown as ribbon style for MasL (**a**) and MasL-collimonin C complex (**b**). Four subunits of homogeneous tetramer are distinguished as red, orange, light green, and cyan. Collimonin C is presented as space-filling style in the complex.

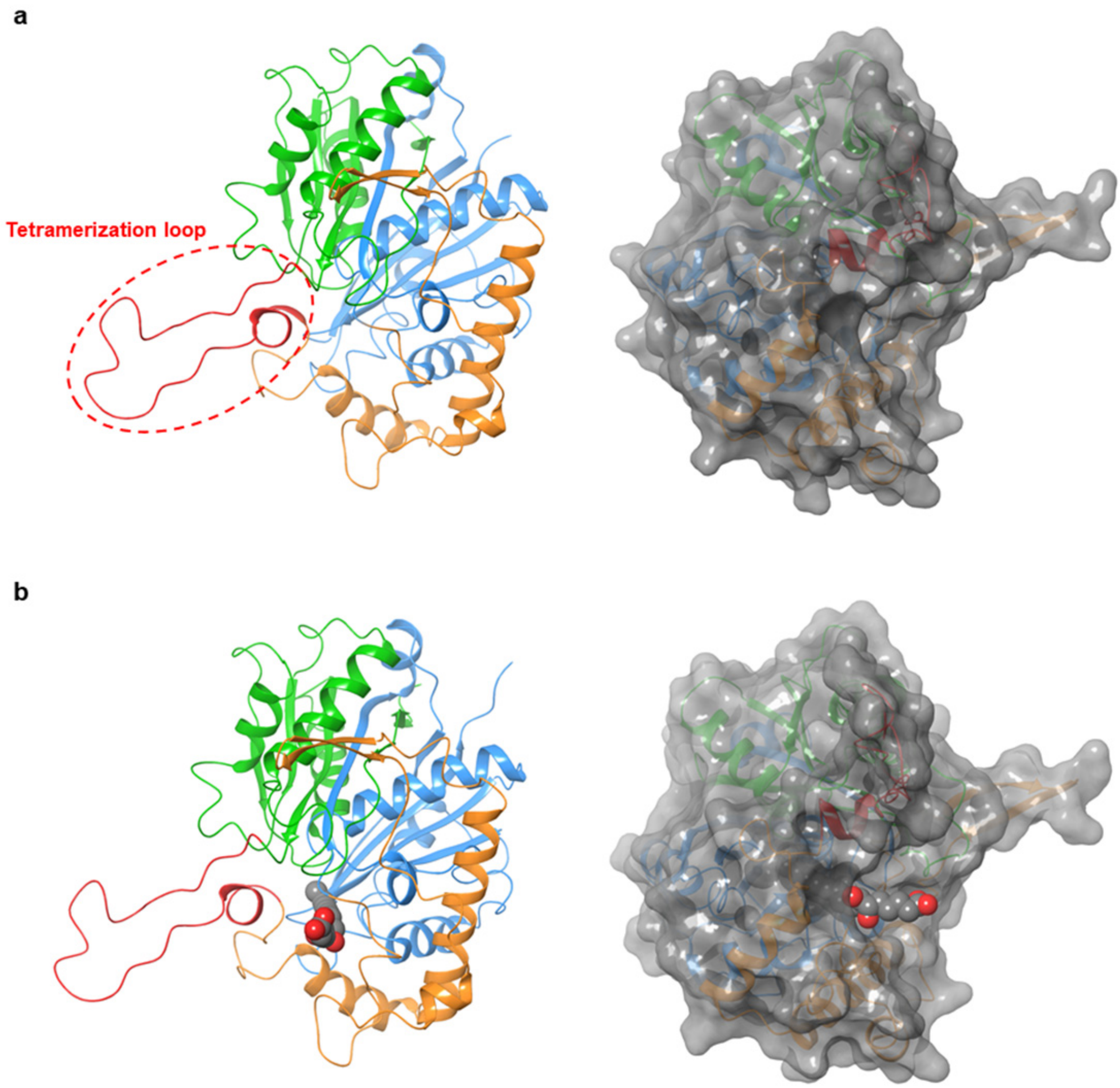


**Supplementary Figure 11 |** Monomeric structure of MasL covalently modified by collimonin C. Monomeric structure of MasL without (**a**) and with collimonin C (**b**) are shown as ribbon representation in which the N-, C- and L-domains are distinguished with green, orange, and blue, respectively. The tetramerization loop is colored red, and the molecular surface is created with gray color. Collimonin C is presented as space-filling style in the complex.

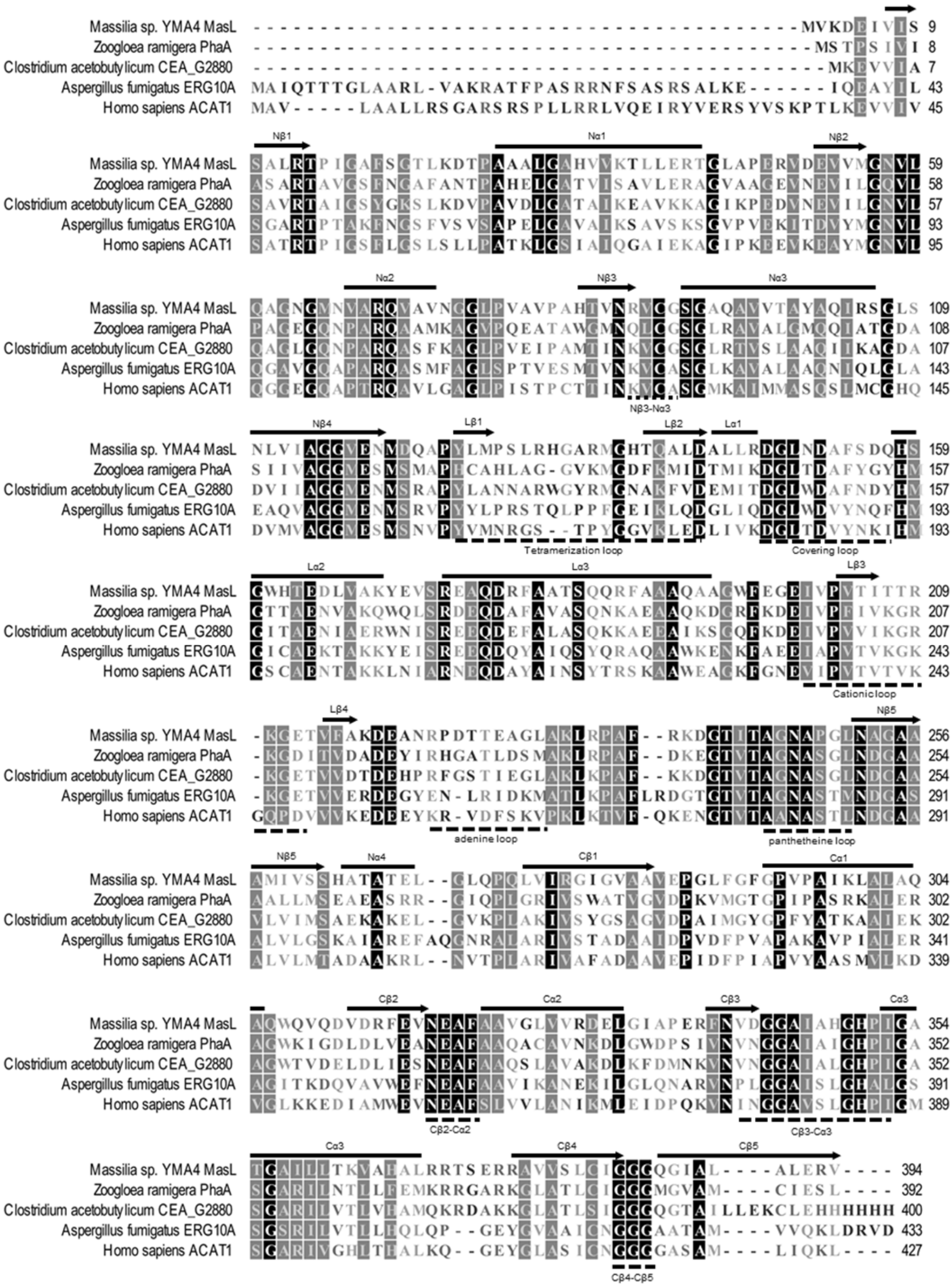

**Supplementary Figure 12 |** Sequence alignment of acetyl-CoA acetyltransferases from different organisms. Amino acid sequence alignment of selected acetyl-CoA acetyltransferases (*Massilia* sp. YMA4 MasL; *Zoogloea ramigera* PhaA, **1QFL**; *Clostridium acetobutylicum* CEA_G2880, **4XL4**; *Aspergillus fumigatus* ERG10A, **6L2C**; *Homo sapiens* ACAT1, **2IBU**). Conserved residue over 80% are colored in gray, and 100% are colored in black.

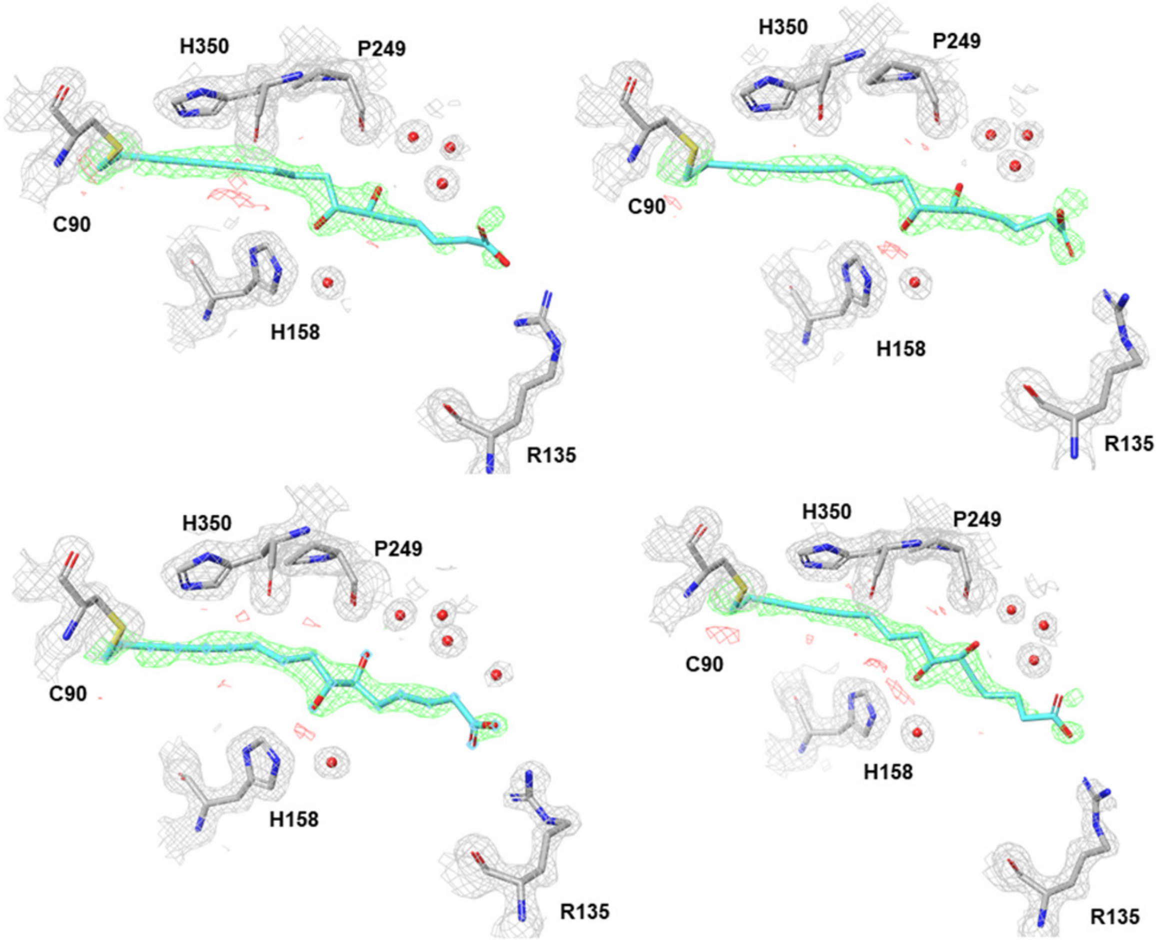


**Supplementary Figure 13 |** Electron density map of collimonin C in MasL reactive pocket. The initial $F_o - F_c$ electron density map contoured at 1.2$\sigma$ around the collimonin C (density in cyan) with refined $2F_o - F_c$ electron density contoured at 1.6$\sigma$ for enzyme residues of MasL-collimonin C complex in four subunits. Collimonin C carbons are shown in cyan, and enzyme carbons are shown in grey, oxygens in red, nitrogens in blue, and sulfurs in yellow.

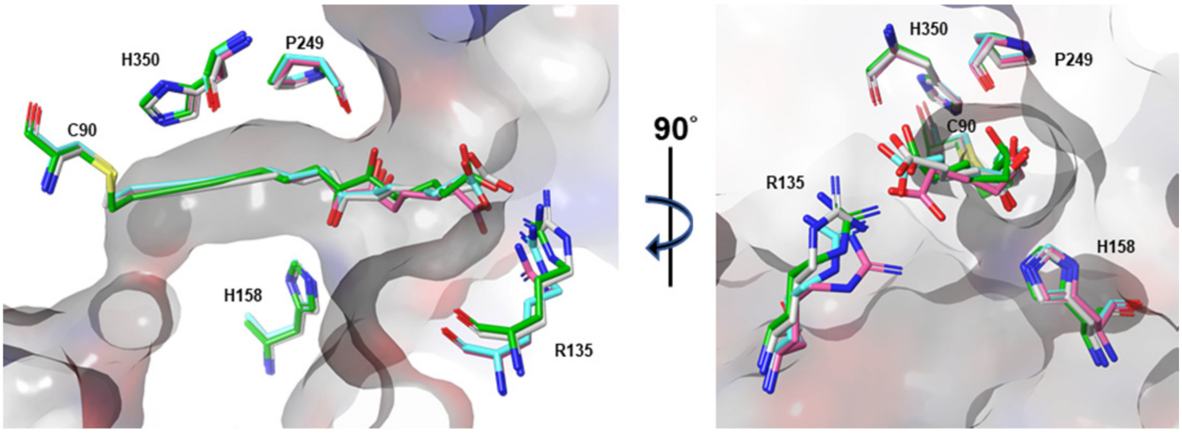


**Supplementary Figure 14 |** Magnification view of MasL covalently modified by collimonin C. Superimposition of collimonin C in four subunits, colored with gray, cyan, green, and pink for each chain, respectively. The residues involved in hydrogen interactions are shown in stick representation with their respective sequence identities indicated. The protein surface of MasL subunit A is colored with the corresponding residual charge for positive (blue) and negative (red).

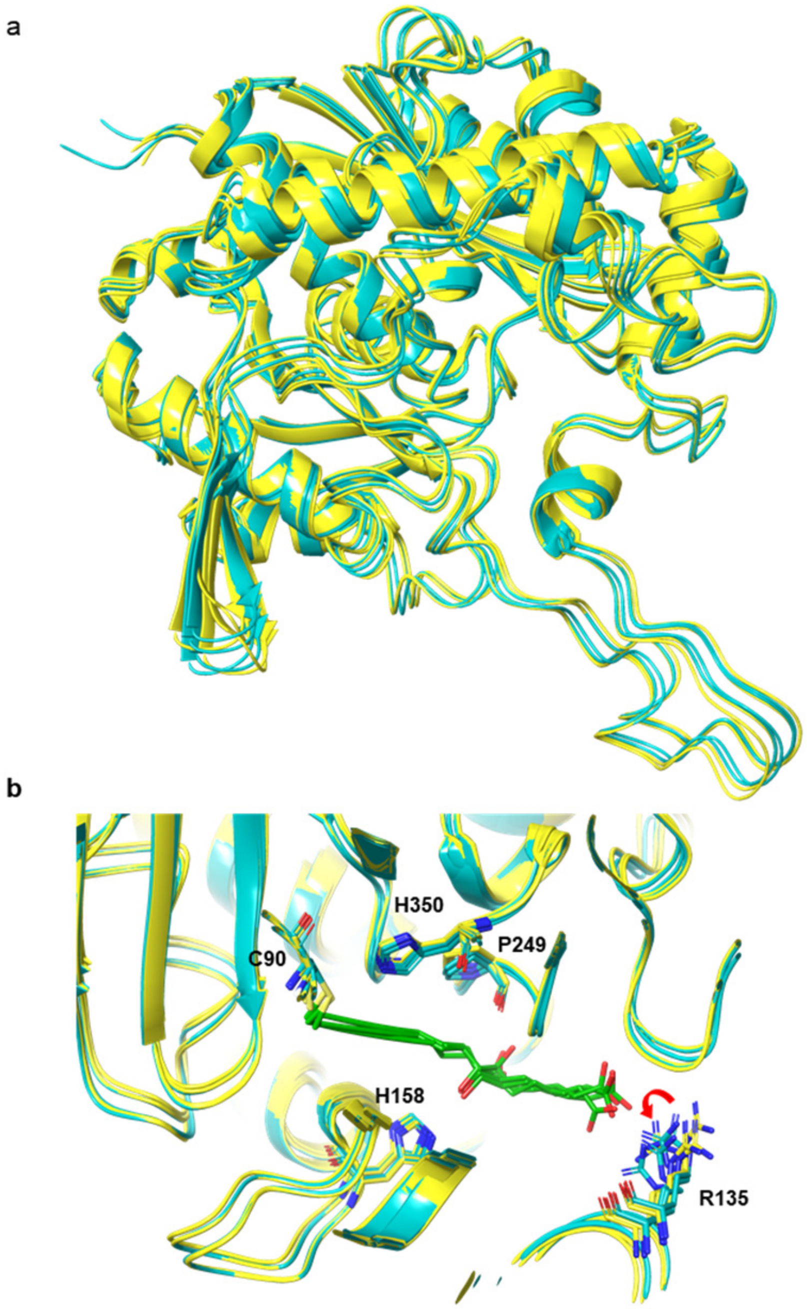


**Supplementary Figure 15 |** Superimposition of MasL and MasL-collimonin C complex. (**a**) Collimonin C binding site of MasL (yellow) and MasL-collimonin C complex (cyan) were superimposed. The monomeric proteins are shown as ribbon representation. (**b**) Magnification view of the binding pocket. The residues involved in polar interactions are shown in stick style with their respective sequence identities indicated and colored in corresponding backbone color. Collimonin C is colored in green and presented as stick style in the complex. The red curve arrow indicates induced-fit movement.

| Inhibitors | $K_I$ (μM) | $k_{inact}$ ($min^{-1}$) | Ref. |
|---|---|---|---|
| 5-Chloro-4-oxopentanoic acid **(Aa-1)** | 15000 | 0.40 ± 0.18 | |
| 7-Chloro-6-oxoheptanoic acid **(Aa-2)** | 11040 ± 1200 | 0.53 ± 0.30 | |
| 9-Chloro-8-oxononanoic acid **(Aa-3)** | 490 ± 100 | 0.52 ± 0.09 | |
| 11-Chloro-10-oxoundecanoic acid **(Aa-4)** | 28 ± 5.8 | 4.0 ± 0.7 | 1 |
| 5-Chloro-4-oxopentanoyl-CoA **(Ab-1)** | 15 ± 1.5 | 2.5 ± 0.2 | |
| 7-Chloro-6-oxoheptanoyl-CoA **(Ab-2)** | 2 ± 0.2 | 2.7 ± 0.4 | |
| 9-Chloro-8-oxononanoyl-CoA **(Ab-3)** | 1.4 ± 0.35 | 2.7 ± 0.3 | |
| 11-Chloro-10-oxoundecanoyl-CoA **(Ab-4)** | 2.5 ± 0.4 | 3.0 ± 0.5 | |
| 3-Pentynoyl-CoA **(B-1)** | 25 | 0.33 | 2 |
| 4-Bromocrotonyl-CoA **(B-2)** | 12.5 | 0.5 | |
| 3-Pentenoyl-S-pantetheine 11-pivalate **(C-1)** | 1250 | 0.26 | 3 |
| 2,3-Pentadienoyl-S-pantetheine 11-pivalate **(C-2)** | 1540 | 1.9 | |

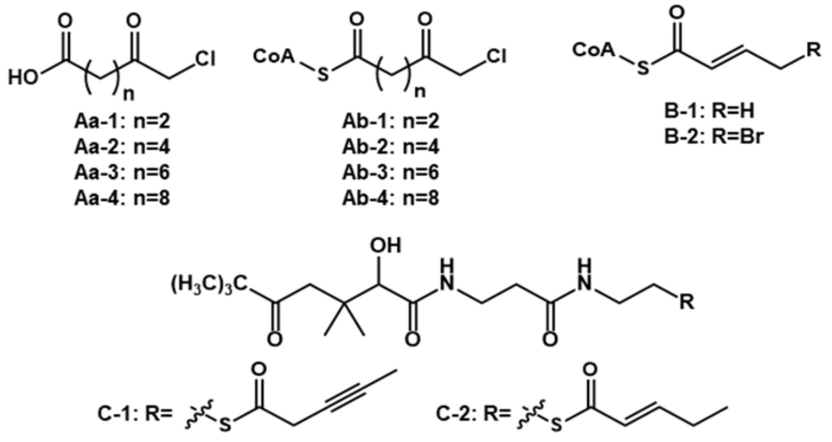


**Supplementary Figure 16 |** Reported inhibitors of acetyl-CoA acetyltransferase (EC 2.3.1.9).

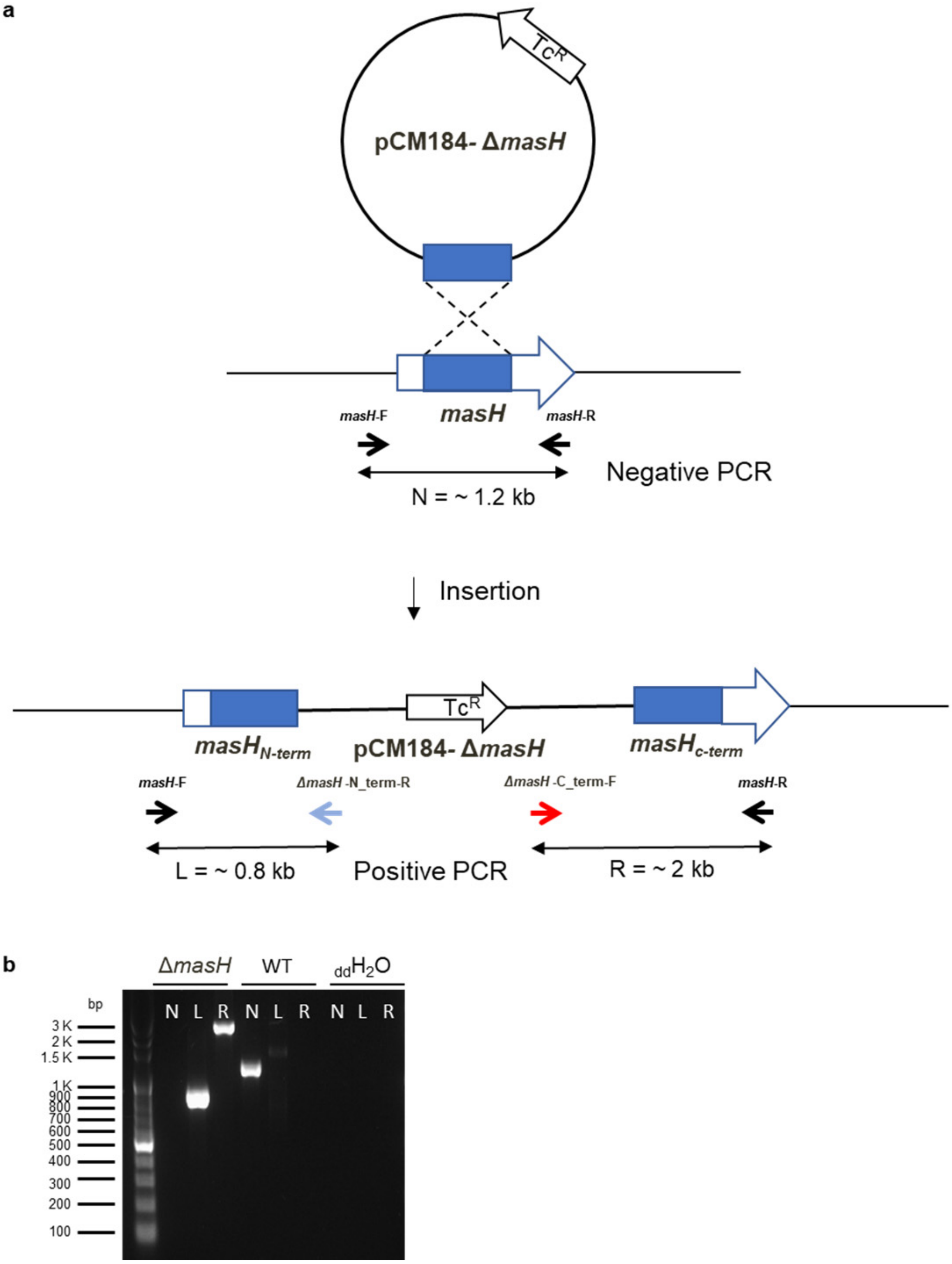


**Supplementary Figure 17 |** Construction scheme (**a**) and PCR check result (**b**) of the null-mutant strain Δ*masH*. The primer sets of PCR check for Δ*masH*: N: *masH*-F + *masH*-R; L: *masH*-F + Δ*masH*-N_term-R; R: Δ*masH*-C_term-F + *masH*-R. The positions of the primer sets are shown in (**a**).

a

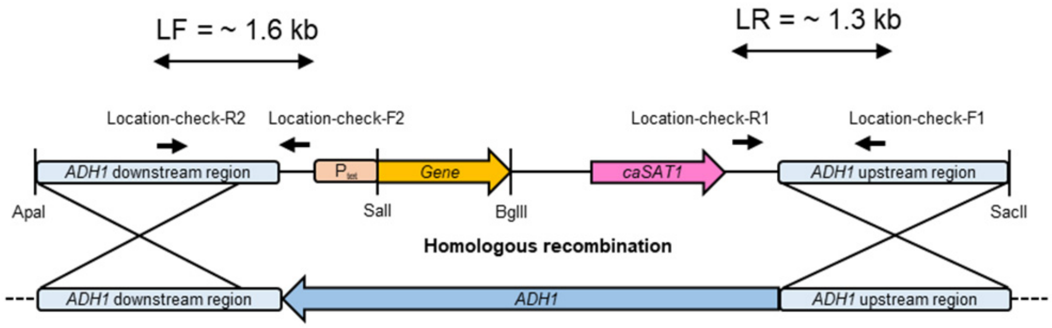


b

>$masL_{opt}$ [*Massilia sp.* YMA4]

ATGAAGGATGAAATTGTTATTTCCTCCGCTTTAAGAACTCCAATTGGTGCTTTTTCAGGTACTTTGAAAGATACT
CCAGCTGCTGCTTTGGGTGCTCATGTTGTTAAAACTTTGTTGGAAAGAACTGGTTTAGCTCCAGAAAGAGTTGA
TGAAGTTGTTATGGGTAATGTTTTGCAAGCTGGTAATGGTATGAATGTTGCTAGACAAGTTGCTGTTAATGGTG
GTTTGCCAGTTGCTGTTCCAGCTCATACTGTTAATAGAGTTTGTGGTTCAGGTGCTCAAGCTGTTGTTACTGCT
TATGCTCAAATTAGATCAGGTTTGTCTAATTTGGTTATTGCTGGTGGTGTTGAAAATATGGATCAAGCTCCATAT
TTGATGCCATCATTAAGACATGGTGCTAGAATGGGTCATACTCAAGCTTTGGATGCTTTGTTGAGAGATGGTTT
AAATGATGCTTTTTCTGATCAACATTCCGGTTGGCATACTGAAGATTTGGTTGCTAAATATGAAGTTTCAAGAGA
AGCTCAAGATAGATTTGCTGCTACTTCTCAACAAAGATTTGCTGCAGCTCAAGCTGCTGGTTGGTTTGAAGGTG
AAATTGTTCCAGTTACTATTACTACTAGAAAGGGTGAAACTGTTTTTGCTAAAGATGAAGCTAATAGACCAGATA
CTACTGAAGCTGGTTTAGCTAAATTGAGACCAGCTTTTAGAAAAGATGGTACTATTACTGCTGGTAATGCTCCA
GGTTTGAATGCTGGTGCTGCTGCTATGATTGTTTCATCACATGCTACTGCTACTGAATTAGGTTTGCAACCACA
ATTGGTTATTAGAGGTATTGGTGTTGCTGCTGTTGAACCAGGTTTGTTTGGTTTTGGTCCAGTTCCAGCTATTA
AGTTAGCTTTAGCTCAAGCTCAATGGCAAGTTCAAGATGTTGATAGATTTGAAGTTAACGAAGCTTTTGCTGCT
GTTGGTTTAGTTGTTAGAGATGAATTGGGTATTGCTCCAGAAAGATTCAATGTTGATGGTGGTGCTATTGCTCA
TGGTCATCCAATTGGTGCAACTGGTGCTATTTTATTAACTAAAGTTGCTCATGCTTTGAGAAGAACTTCAGAAA
GAAGAGCTGTTGTTTCATTGTGTATTGGTGGTGGTCAAGGTATTGCTTTAGCTTTAGAAAGAGTTTAA

c

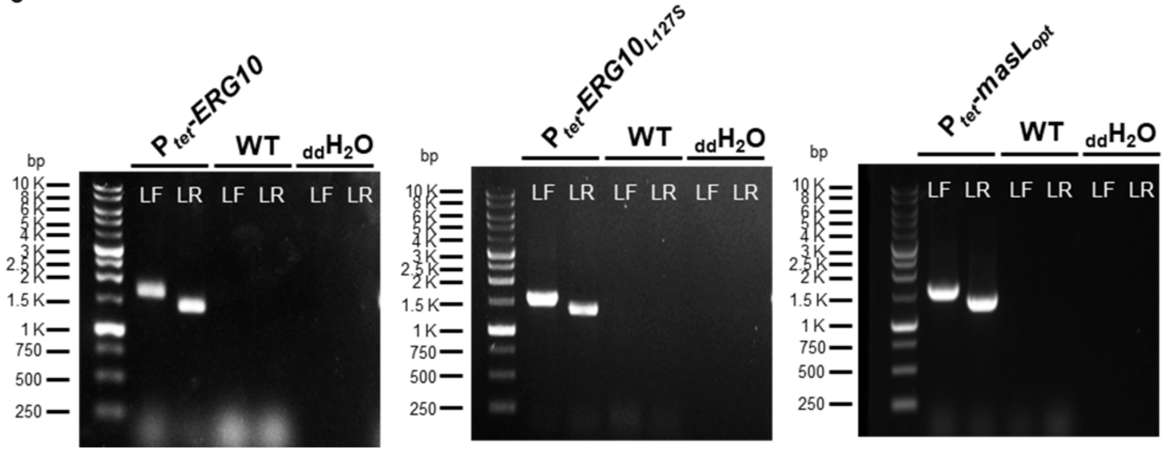


**Supplementary Figure 18 |** Tetracycline-inducible expression system in *C. albicans* ATCC18804. (**a**) Scheme of the inducible overexpression of *CaERG10* and Codon optimized *masL* ($masL_{opt}$) constructed by homologous recombination replacing with *ADH1* gene in the chromosome. (**b**) Codon optimized *masL* sequence for overexpression in *C. albicans* ATCC18804. (**c**) PCR check results of tetracycline-inducible *ERG10*, $ERG10_{L127S}$, and $masL_{opt}$ strains. The positions of the primer sets are shown in (**a**).

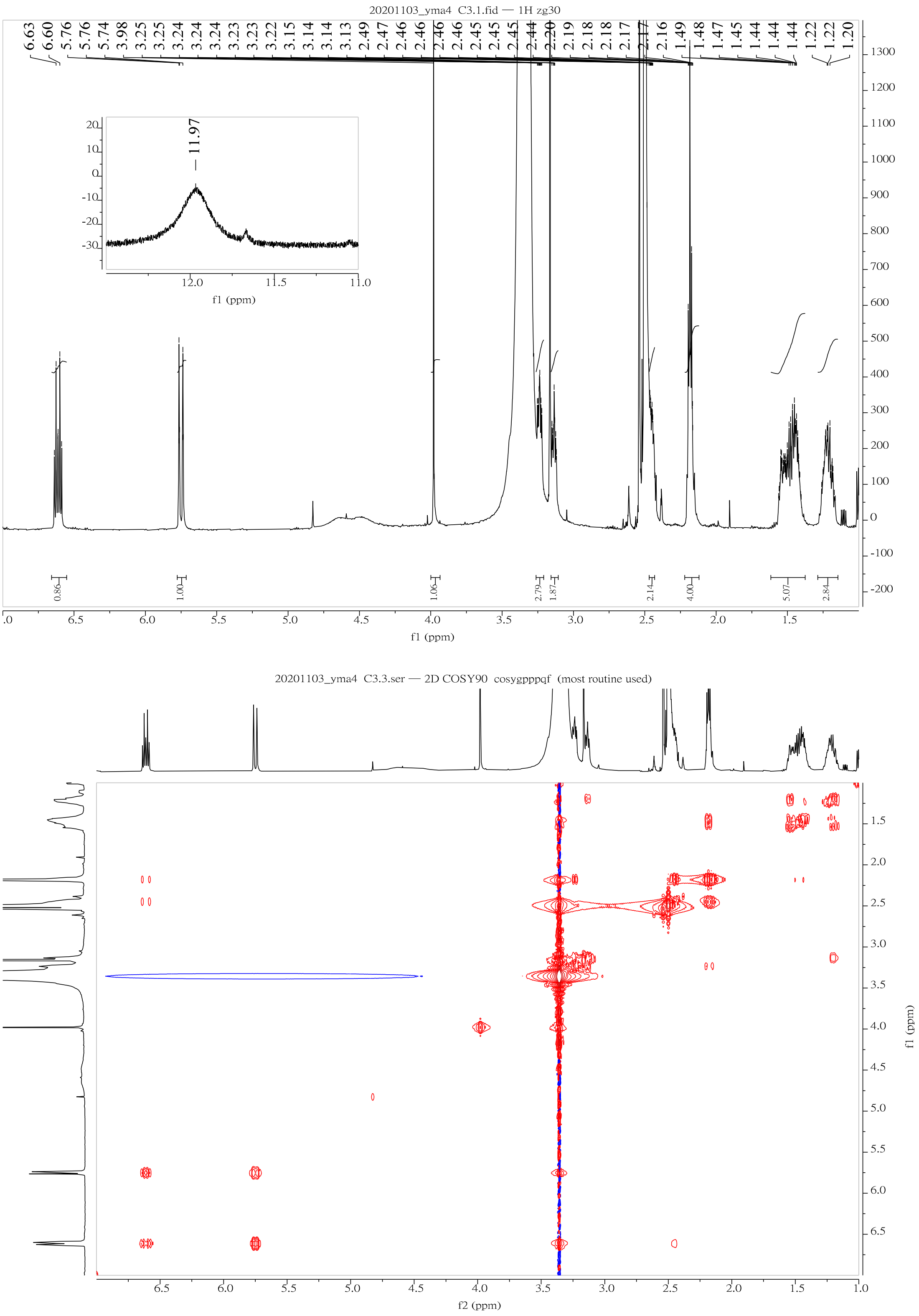


**Supplementary Figure 19** | $^{1}$H NMR and $^{1}$H-$^{1}$H COSY (600 MHz) of collimonin C **1**

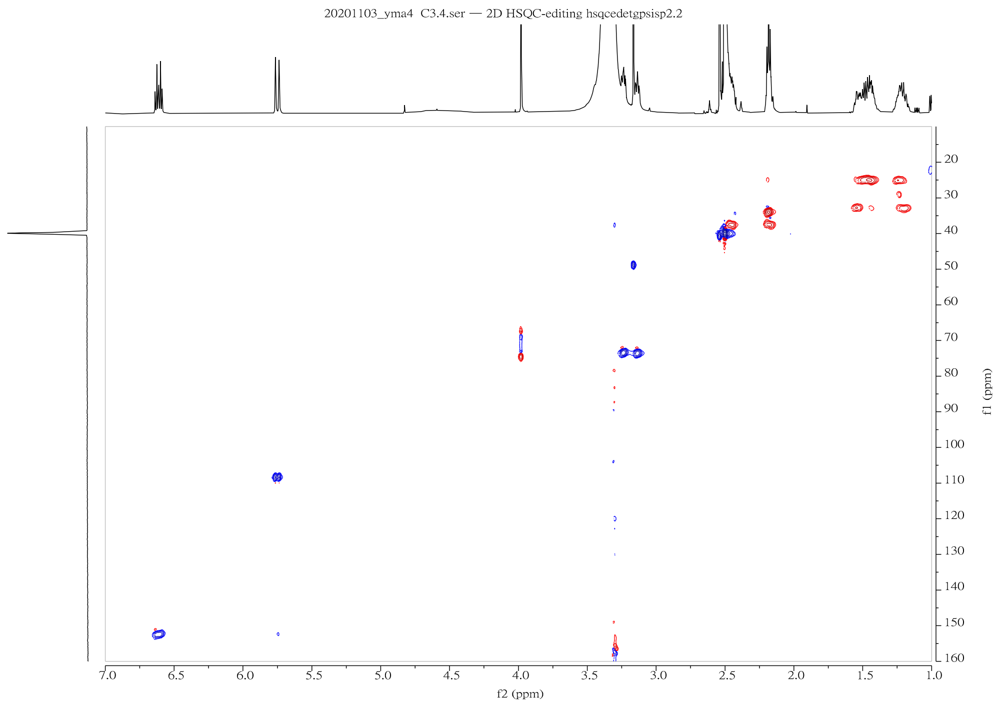


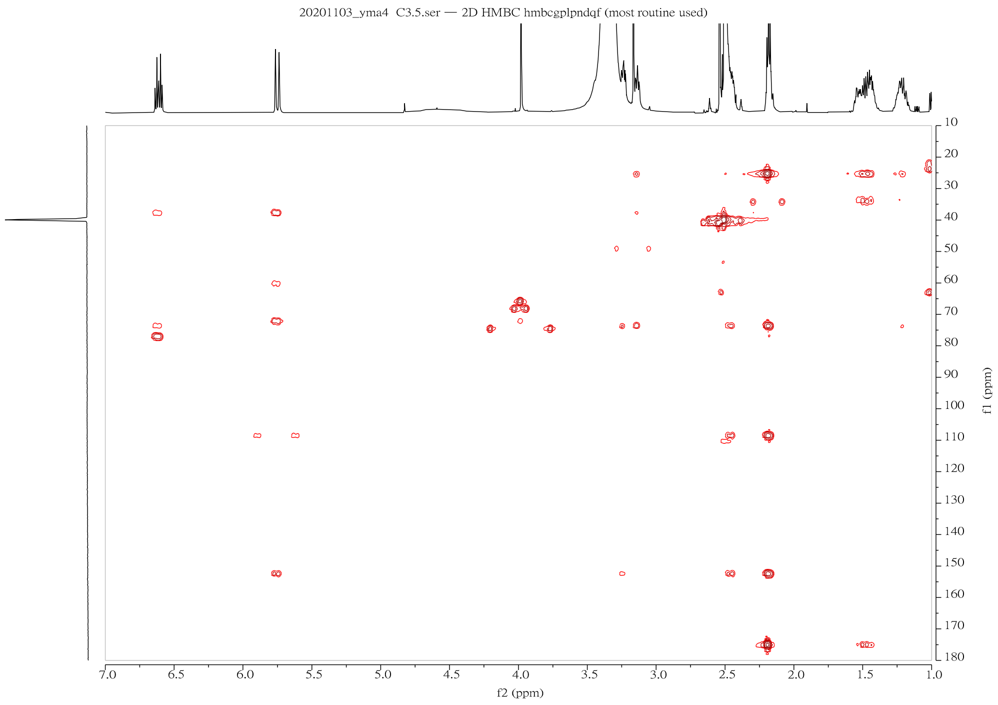


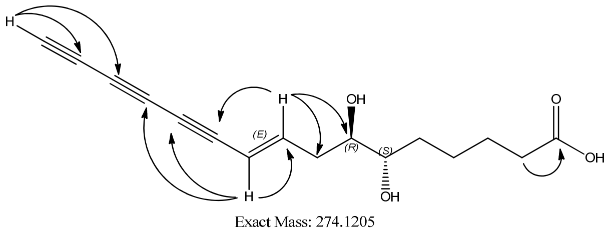


→ Key HMBC correlation of Collimonin C **1**

**Supplementary Figure 20 |** HSQC and HMBC (600 MHz) of collimonin C **1**

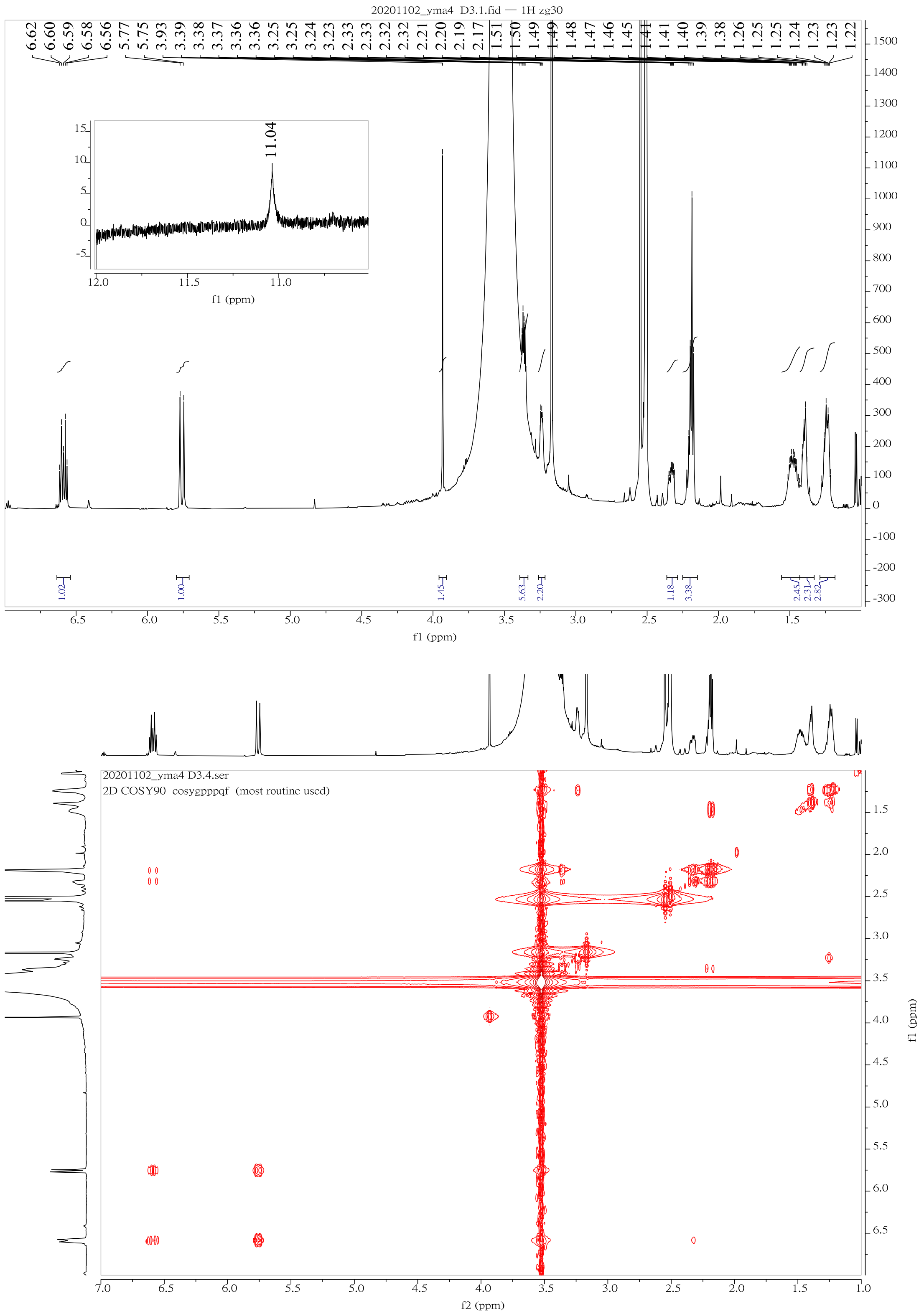


**Supplementary Figure 21 |** $^{1}$H NMR and $^{1}$H-$^{1}$H COSY (600 MHz) of collimonin D **2**

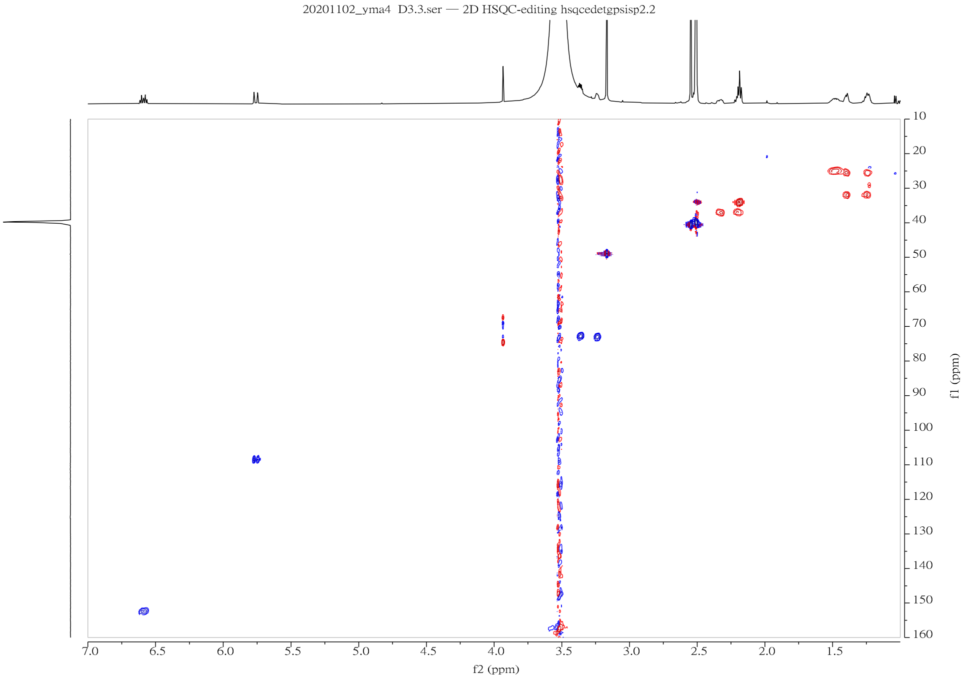


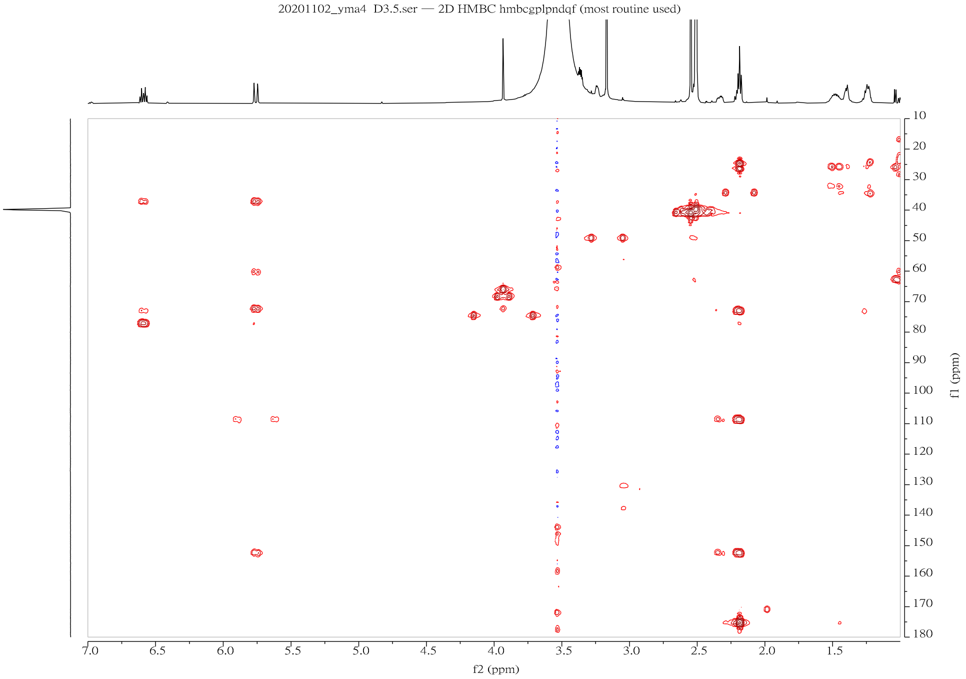


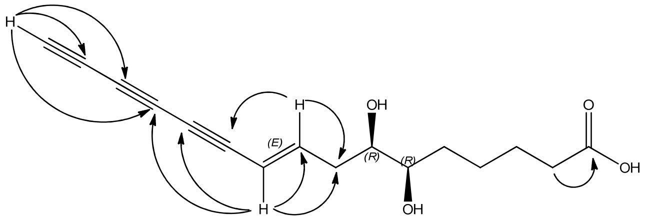


→ Key HMBC correlation of Collimonin D **2**

**Supplementary Figure 22 |** HSQC and HMBC (600 MHz) of collimonin D **2**

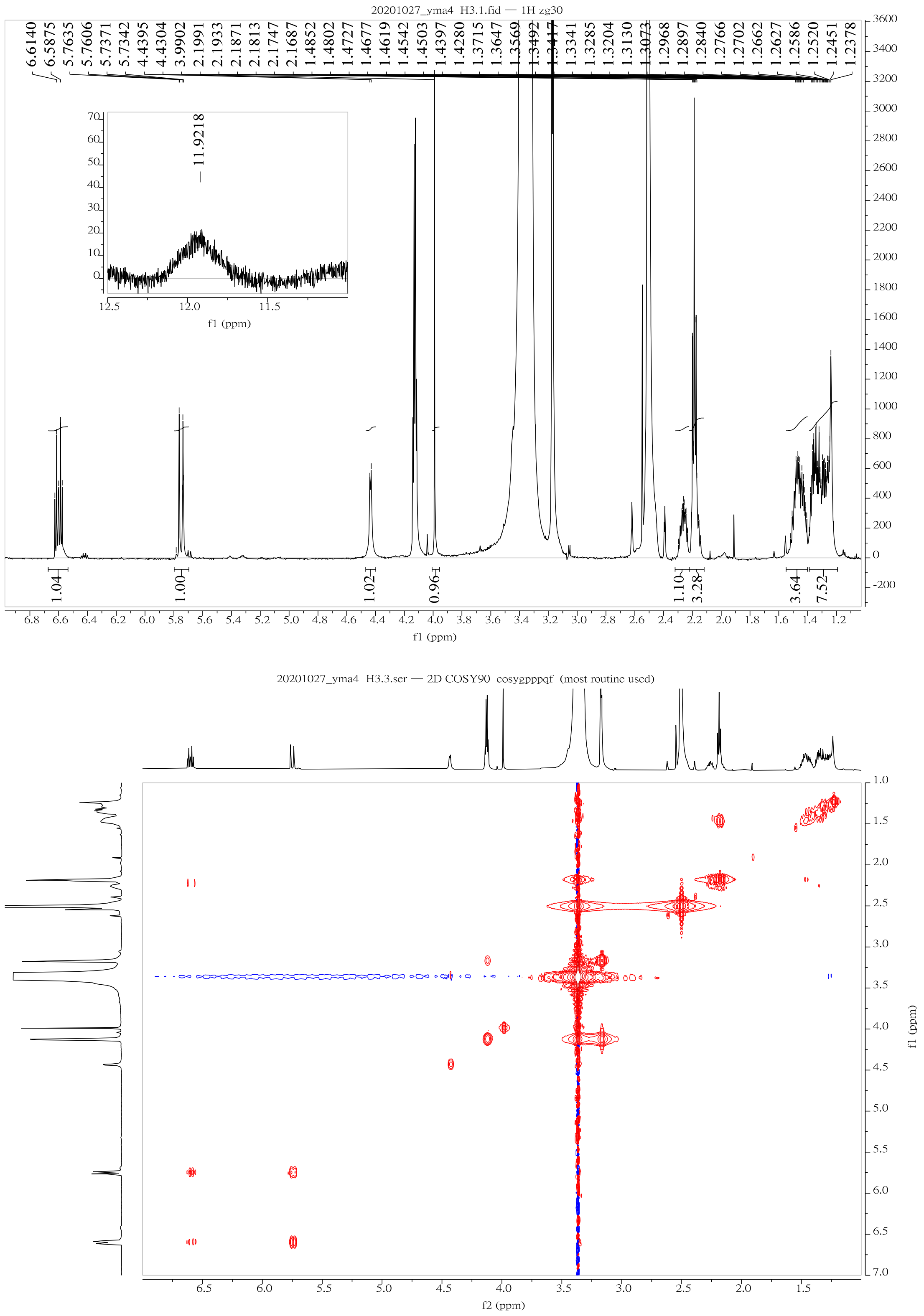

**Supplementary Figure 23 |** $^{1}H$ NMR and $^{1}H$-$^{1}H$ COSY (600 MHz) of massilin A **3**

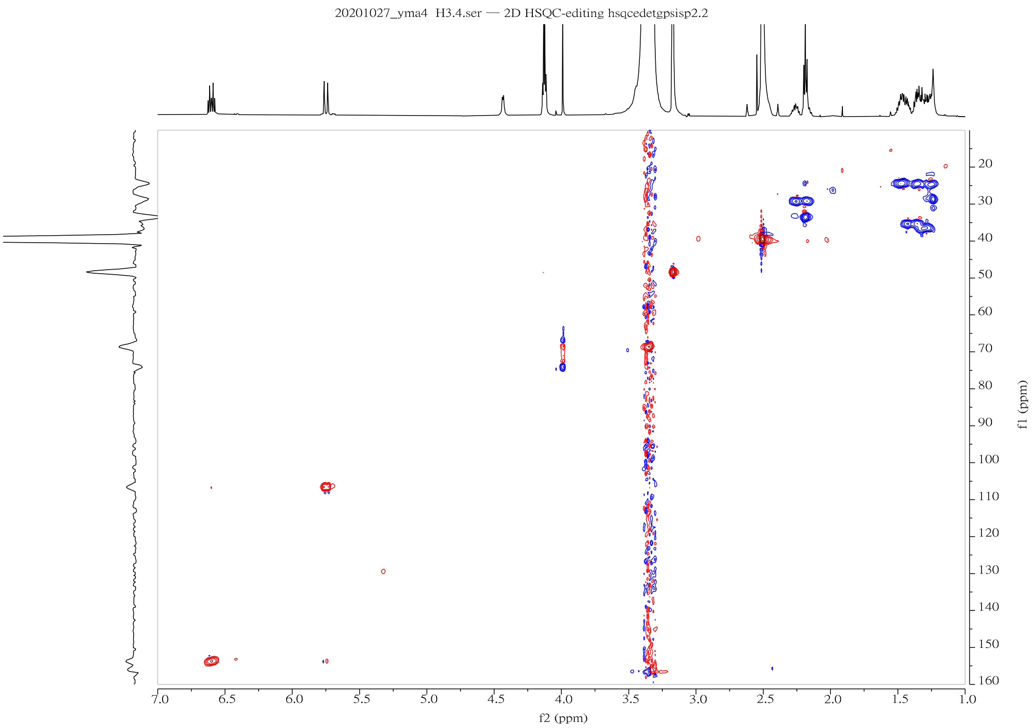


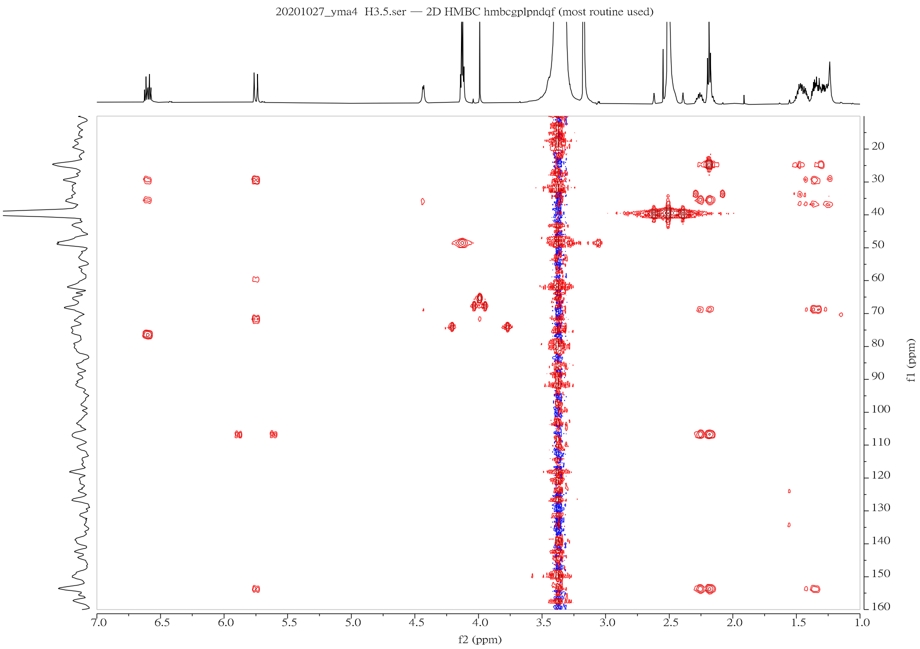


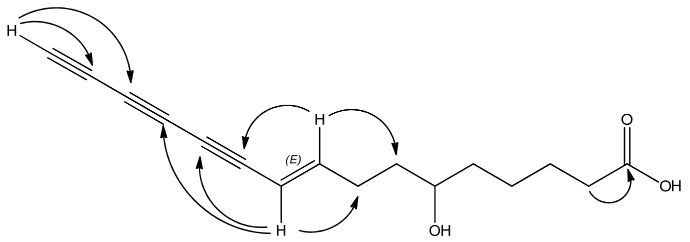


→ Key HMBC correlation of Massilin A **3**

**Supplementary Figure 24 |** HSQC and HMBC (600 MHz) of massilin A **3**

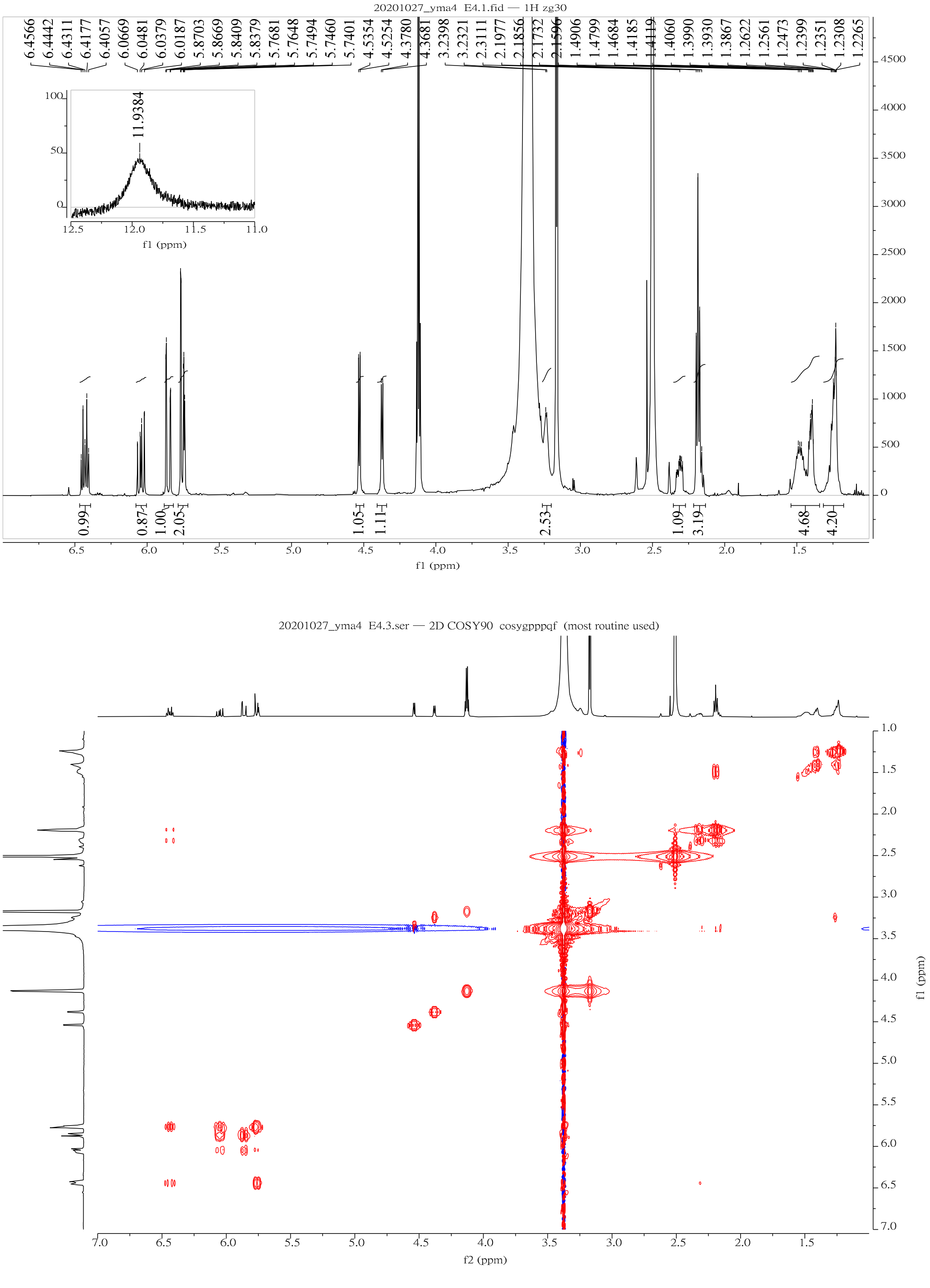


**Supplementary Figure 25 |** $^1$H NMR and $^1$H-$^1$H COSY (600 MHz) of massilin B **4**

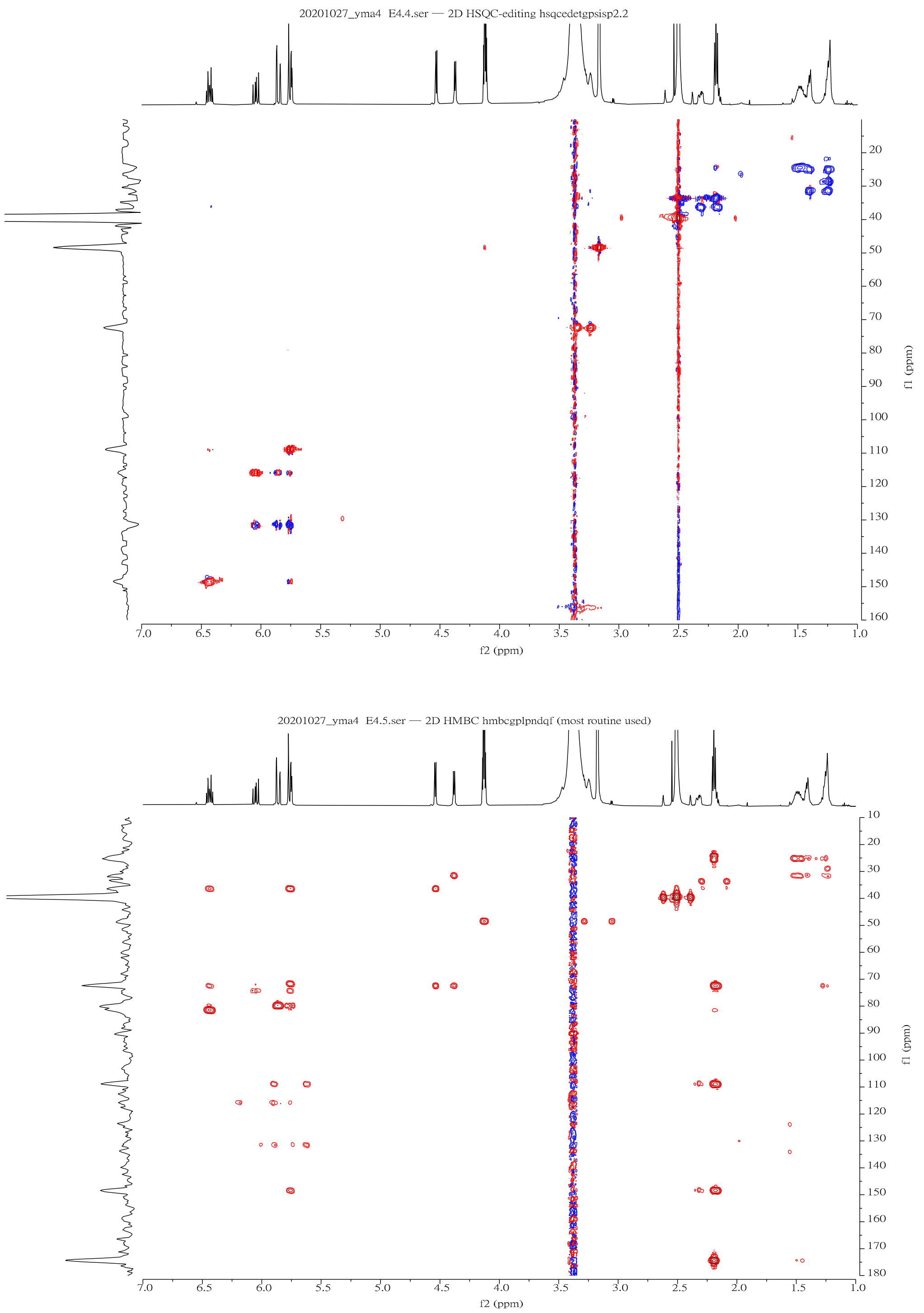


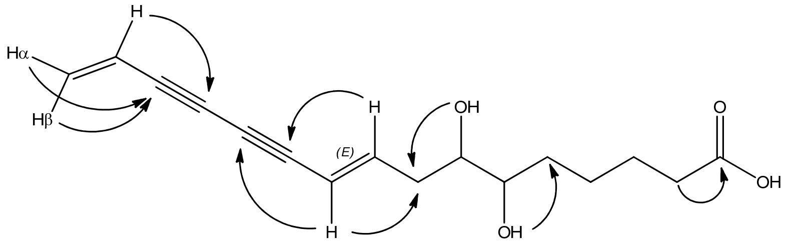


→ Key HMBC correlation of massilin B **4**

**Supplementary Figure 26 |** HSQC and HMBC (600 MHz) of massilin B **4**

#### Supplementary Tables

**Supplementary Table 1 |** Characterization of *mas* biosynthetic gene cluster

| **Gene Name** | **Length bp / aa** | **Proposed Function** | **Similar Proteins (*C. fungivorans* Ter331 and *B. ambifaria* BCC0191)** | **Sequence Identity (%aa)** |
|---|---|---|---|---|
| ***masA*** | 1080 / 359 | Fatty acid desaturase | fatty_acid_desaturase [*C. fungivorans* Ter331]<br>fatty_acid_desaturase [*B. ambifaria* BCC0191], *ccnF* | 70%<br>73% |
| ***masB*** | 1005 / 334 | Aromatic ring-hydroxylating dioxygenase subunit alpha | Rieske_(2Fe-2S) domain_protein [*C. fungivorans* Ter331]<br>aromatic_ring-hydroxylating_dioxygenase_subunit alpha [*B. ambifaria* BCC0191], *ccnG* | 86%<br>83% |
| ***masC*** | 822 / 273 | Hypothetical protein | hypothetical_protein [*B. ambifaria* BCC0191], *ccnH* | 72% |
| ***masD*** | 1854 / 617 | Fatty_acyl-AMP_ligase | Long-chain-fatty-acid--CoA_ligase [*C. fungivorans* Ter331], *colA*<br>fatty_acyl-AMP_ligase [*B. ambifaria* BCC0191], *ccnJ* | 77%<br>52% |
| ***masE*** | 903 / 300 | Acyl-CoA desaturase | delta-9_fatty_acid_desaturase [*C. fungivorans* Ter331], *colB*<br>acyl-CoA_desaturase [*B. ambifaria* BCC0191], *ccnK* | 88%<br>58% |
| ***masF*** | 981 / 326 | Acyl-CoA desaturase | delta-9_fatty_acid_desaturase [*C. fungivorans* Ter331], colC<br>acyl-CoA_desaturase [*B. ambifaria* BCC0191], *ccnL* | 89%<br>75% |

**Supplementary Table 1 (Continued) |** Characterization of *mas* biosynthetic gene cluster

| | | | | |
|---|---|---|---|---|
| ***masG*** | 330 / 109 | Acyl_carrier_protein | phosphopantetheine-binding_protein [*C. fungivorans* Ter331], *colD*<br>acyl_carrier_protein [*B. ambifaria* BCC0191], *ccnM* | 94%<br>66% |
| ***masH*** | 1101 / 365 | Fatty acid desaturase | fatty_acid_desaturase [*C. fungivorans* Ter331], *colE*<br>fatty_acid_desaturase [*B. ambifaria* BCC0191], *ccnN* | 88%<br>68% |
| ***masI*** | 945 / 314 | Alpha/beta hydrolase | delta_12_desaturase [*C. fungivorans* Ter331], *colF*<br>alpha/beta_fold_hydrolase [*B. ambifaria* BCC0191], *ccnO* | 82%<br>52% |
| ***masJ*** | 186 / 60 | Rubredoxin | rubredoxin-type_Fe(Cys)4_protein [*C. fungivorans* Ter331]<br>rubredoxin [*B. ambifaria* BCC0191], *ccnP* | 86%<br>68% |
| ***masK*** | 918 / 305 | Alpha/beta hydrolase | none | - |
| ***masL*** | 1182 / 393 | Acetyl-CoA acetyltransferase | acetyl-CoA_C-acyltransferase [*B. ambifaria* BCC0191] | 59% |

**Supplementary Table 2 |** Characterization of core genes in cepacin biosynthetic gene cluster

| Gene Name | Length bp / aa | Proposed Function | Similar Proteins (*C. fungivorans* Ter331) | Sequence Identity (%aa) |
|---|---|---|---|---|
| ***ccnD*** | 1293 / 430 | Beta-ketoacyl synthase | 3-oxoacyl-(acyl-carrier-protein) synthase [*C. fungivorans* Ter331] | 75% |
| ***ccnE*** | 1386 / 461 | Flavin-dependent monooxygenase | monooxygenase, FAD-binding_protein [*C. fungivorans* Ter331] | 88% |
| ***ccnI*** | 1419 / 472 | MFS_transporter | drug_transporter [*C. fungivorans* Ter331] | 84% |

**Supplementary Table 3 |** List of primer sets used in this study

| **Real-time qPCR** | **Sequence (5'->3')** |
| --- | --- |
| *Act1*-qPCR-F | AGGTTTGGAAGCTGCTGGTA |
| *Act1*-qPCR-R | GAAGATGGAGCCAAAGCAGT |
| *ERG10*-qPCR-F | GAAGCCGCCAGAAATCCAAT |
| *ERG10*-qPCR-R | CAACAGCACCACCATAAGCA |
| **Vector construction** | |
| *masH*-fF | GCAAAAAGCTACCCATCATCGC |
| *masH*-fR | TCAGCCGAAGTAGACGATCGAGG |
| *ERG10*-F | GGAGCGGTCGACATCATTATGGTCCCACCTGTTTA |
| *ERG10*-R | GGAGCGAGATCTTTTGTCGTCCCCCTTACAATTTA |
| *ERG10His*-R | GAGCGAGATCTTAATGATGATGATGATGATGCAATTT AAAGTCACTATCGACTTTTTCAA |
| **Constructed strain examination** | |
| *masH*-F | CTCAACGCAACCGCCTAAAC |
| *masH*-R | TCAGGTATTCGGCACGGAGG |
| Δ*masH* -C_term-F | TTACCAATGCTTAATCAGTGAG |
| Δ*masH* -N_term-R | CGAACGACATGGAGCGGCAC |
| Location-check-F1 | CCGAATTATTCCGGAAGCTGGTAGC |
| Location-check-R1 | AAAGGGCAAAAGTGAGTATGGTGCC |
| Location-check-F2 | GCCCATCAGAAACGACAAACATGGA |
| Location-check-R2 | ACAATCAATGCCAGAGATCAAACCA |
| **Recombinant protein production** | |
| MasL-F | GGAATTCCATATGAAAGATGAAATCGTCATCAGTTC |
| MasL-R | CCCAAGCTTCACGCGTTCCAGCGCCAGCGCGATG |
| *hsACAT1*-F | AAGGAGATATACATATGGTATCAAAACCCACTTTGAA GGAAG |
| *hsACAT1*-R | GGTGGTGGTGCTCGAGCAGCTTCTGAATTAGCATGG CAG |

**Supplementary Table 4 |** List of strains used in this study

| **Strain** | **Description** | **Source** |
|---|---|---|
| *Massilia* sp. YMA4 | Wild-type strain; $Km^r$ | This study |
| Δ*masH* | Biosynthesis mutant strain derived from *Massilia* sp. YMA4 by plasmid insertion at *masH* locus; $Km^r$, $Tc^r$ | This study |
| *Candida albicans* ATCC18804 | Wild-type strain | BCRC |
| $P_{tet}$-*ERG10* | *ERG10* overexpression strain derived from *Candida albicans* ATCC18804; $Nc^r$ | This study |
| $P_{tet}$-$ERG10_{L127S}$-His | Recombinant protein production of C. *albicans* ATCC18804 with point mutation L127S and C-terminal 6xHis tag; $Nc^r$ | This study |
| $P_{tet}$-$masL_{opt}$-His | $masL_{opt}$ with 6xHis tag heterologous expression strain derived from *Candida albicans* ATCC18804; $Nc^r$ | This study |
| *Candida albicans* | Clinical isolate from National Taiwan University Hospital | Prof. Ching-Hsuan Lin[4] |
| *Candida tropicalis* | Clinical isolate from National Taiwan University Hospital | Prof. Ching-Hsuan Lin[4] |
| *Escherichia coli* S17-1 λ *pir* | *Escherichia coli* donor strain for biparental conjugation | Prof. Nai-Chun Lin |
| *Escherichia coli* C41(DE3) | Recombinant protein production of *Massilia* sp.YMA4 MasL and human ACAT1 | Yeastern Biotech |
| Human prostate PC-3 cell line | *Homo sapiens* prostate cancer cell line; derived from metastatic site: bone | Dr. Pei-Wen Hsiao |

Note: $Km^r$, $Tc^r$, $Ap^r$, and $Nc^r$ indicate resistance to kanamycin, tetracycline, ampicillin, and nourseothricin.

**Supplementary Table 5 |** List of plasmids used in this study

| Plasmid | Description | Source |
|---|---|---|
| pCM184 | Broad-host-range allelic exchange vector; $Km^r$, $Tc^r$, $Ap^r$ | Addgene[5] |
| pCM184-Δ*masH* | pCM184-Δ*masH* for insertion on *masH* gene; $Tc^r$, $Ap^r$ | This study |
| pNIM1 | Plasmid for tetracycline-regulated gene expression system with GFP expression; $Nc^r$, $Ap^r$ | Yang-Nim Park and Joachim Morschhäuser[6] |
| pET22b | Vector for construction C-terminal His-tagged recombinant protein; $Ap^r$ | Novagen |
| pET28a | Vector for construction N and C-terminal His-tagged recombinant protein; $Km^r$ | Novagen |
| pNIM-ERG10 | Tetracycline-inducible *CaERG10* overexpression plasmid derived from pNIM1 by replacing GFP gene; $Nc^r$, $Ap^r$ | This study |
| pNIM-ERG10$_{L127S}$ | Tetracycline-inducible *CaERG10*$_{L127S}$ overexpression plasmid derived from pNIM1 by replacing GFP gene; $Nc^r$, $Ap^r$ | This study |
| pNIM-MasL$_{opt}$ | Tetracycline-inducible *masL*$_{opt}$ overexpression plasmid derived from pNIM1 by replacing GFP gene; $Nc^r$, $Ap^r$ | This study |
| pET22b-MasL | Recombinant protein production of *Massilia* sp. YMA4 MasL; $Ap^r$ | This study |
| pET28a-ACAT1 | Recombinant protein production of human mitochondrial ACAT1, truncated with N-term transition peptide sequence for aa sequence from V34 to L427; $Km^r$ | This study |

Note: $Km^r$, $Tc^r$, $Ap^r$, and $Nc^r$ indicate resistance to kanamycin, tetracycline, ampicillin, and nourseothricin.

**Supplementary Table 6 |** Whole-genome sequences used in mining bacterial polyyne gene clusters

| Genus | Strains | Accessions |
|---|---|---|
| *Amycolatopsis* | *Amycolatopsis* orientalis strain B-37 | CP016174 |
| *Burkholderia* 1 | *Burkholderia gladioli* BSR3 | CP002599 |
| *Burkholderia* 1 | *Burkholderia gladioli* pv. gladioli strain FDAARGOS_188 | CP022213 |
| *Burkholderia* 1 | *Burkholderia gladioli* pv. gladioli strain FDAARGOS_389 | CP023522 |
| *Burkholderia* 1 | *Burkholderia gladioli* pv. gladioli strain KACC 11889 | CP022005 |
| *Burkholderia* 1 | *Burkholderia gladioli* strain ATCC 10248 | CP009323 |
| *Burkholderia* 1 | *Burkholderia gladioli* strain Co14 | CP033430 |
| *Burkholderia* 1 | *Burkholderia plantarii* strain ATCC 43733 | CP007213 |
| *Burkholderia* 1 | *Burkholderia plantarii* strain PG1 | CP002581 |
| *Burkholderia* 2 | *Burkholderia ambifaria* IOP40-10 | ABLC01000003 |
| *Burkholderia* 2 | *Burkholderia ambifaria* MEX-5 | ABLK01000019 |
| *Burkholderia* 2 | *Burkholderia ambifaria* strain AU11161 | PVFJ01000031 |
| *Burkholderia* 2 | *Burkholderia ambifaria* strain BCC0191 | NZ_CADEQI010000002 |
| *Burkholderia* 2 | *Burkholderia* sp. Bp7605 strain MSMB0175 | CP013448 |
| *Burkholderia* 2 | *Burkholderia* sp. KJ006 | CP003515 |
| *Burkholderia* 2 | *Burkholderia* sp. LA-2-3-30-S1-D2 | CP013383 |
| *Burkholderia* 2 | *Burkholderia* sp. MBR-1 | CP053610 |
| *Burkholderia* 2 | *Burkholderia stagnalis* strain MSMB735WGS | CP013459 |
| *Burkholderia* 2 | *Burkholderia ubonensis* strain MSMB0783 | CP013422 |
| *Burkholderia* 2 | *Burkholderia vietnamiensis* G4 | CP000615 |
| *Burkholderia* 2 | *Burkholderia vietnamiensis* LMG 10929 | CP009630 |
| *Burkholderia* 2 | *Burkholderia vietnamiensis* strain AU1233 | CP013433 |
| *Burkholderia* 2 | *Burkholderia vietnamiensis* strain FDAARGOS_239 | CP020396 |
| *Burkholderia* 2 | *Burkholderia vietnamiensis* strain FL-2-3-30-S1-D0 | CP013394 |
| *Burkholderia* 2 | *Burkholderia vietnamiensis* strain HI2297 | CP013440 |
| *Burkholderia* 2 | *Burkholderia vietnamiensis* strain MSMB608WGS | CP013455 |
| *Collimonas* | *Collimonas fungivorans* strain Ter6 | CP013232 |
| *Collimonas* | *Collimonas fungivorans* Ter331 | CP002745 |
| *Gynuella* | *Gynuella sunshinyii* YC6258 | CP007142 |
| *Massilia* | *Massilia armeniaca* strain ZMN-3 | CP028324 |

**Supplementary Table 6 (continued) |** Whole-genome sequences used in mining bacterial polyyne gene clusters

| Genus | Strains | Accessions |
|---|---|---|
| *Massilia* | *Massilia* sp. YMA4 | CP030092 |
| *Mycobacterium* | *Mycobacterium cookii* JCM 12404 | AP022569 |
| *Mycobacterium* | *Mycobacterium rhodesiae* NBB3 | CP003169 |
| *Nocardia* | *Nocardia brasiliensis* strain AUSMDU00024985 | CP046171 |
| *Pseudomonas* | *Pseudomonas protegens* Cab57 | AP014522 |
| *Pseudomonas* | *Pseudomonas protegens* CHA0 | CP003190 |
| *Pseudomonas* | *Pseudomonas protegens* Pf-5 | CP000076 |
| *Pseudomonas* | *Pseudomonas protegens* strain FD6 | CP031396 |
| *Pseudomonas* | *Pseudomonas protegens* strain FDAARGOS_307 | CP022097 |
| *Pseudomonas* | *Pseudomonas protegens* strain H78 | CP013184 |
| *Pseudomonas* | *Pseudomonas protegens* strain pf5 | CP032358 |
| *Pseudomonas* | *Pseudomonas protegens* strain pf5-k2 | CP032353 |
| *Pseudomonas* | *Pseudomonas protegens* strain pf5-k3 | CP032352 |
| *Pseudomonas* | *Pseudomonas protegens* strain UCT | CP017964 |
| *Pseudomonas* | *Pseudomonas* sp. CMR5c | CP027705 |
| *Streptomyces* | *Streptomyces albireticuli* strain MDJK11 | CP021744 |
| *Streptomyces* | *Streptomyces griseofuscus* strain DSM 40191 | CP051006 |
| *Streptomyces* | *Streptomyces rochei* 7434AN4 | AP018517 |
| *Streptomyces* | *Streptomyces* sp. ADI95-16 | CP033582 |
| *Streptomyces* | *Streptomyces* sp. HF10 | CP047144 |
| *Streptomyces* | *Streptomyces* sp. Mg1 | CP011665 |
| *Streptomyces* | *Streptomyces* sp. NA04227 | CP054918 |
| *Streptomyces* | *Streptomyces* sp. Tu 2975 | CP047140 |
| *Trinickia* | *Paraburkholderia caryophylli* strain Ballard 720 | FXAH01000012 |
| *Trinickia* | *Trinickia caryophylli* strain Ballard 720 | PNXZ01000011 |
| *Trinickia* | *Trinickia caryophylli* strain DSM 50341 | VJZF01000001 |

**Supplementary Table 7 |** 1D NMR of collimonin C **1** in $C_2D_6OS$

| | Collimonin C **1** | |
|---|---|---|
| **No.** | $\delta_H$ (*J* in Hz) | $\delta_C$ |
| **1** | 11.97, brs | 174.4, COOH |
| **2** | 2.18, t, J = 7.22 | 34.0 |
| **3** | 1.47, m | 25.0 |
| **4** | 1.21,1.53, m | 25.3 |
| **5** | 1.18,1.53, m | 32.9 |
| **6** | 3.13, m | 73.6 |
| **7** | 3.24, m | 73.4 |
| **8** | 2.17, m; 2.45, m | 37.5 |
| **9** | 6.61, dt, J = 16.0, 7.38 | 152.1 |
| **10** | 5.75, d, J = 16.0 | 107.9 |
| **11** | | 77.0 |
| **12** | | 60.2 |
| **13** | | 72.2 |
| **14** | | 65.9 |
| **15** | | 68.1 |
| **16** | 3.98, s | 74.2 |

Note1: 600 MHz for $^1H$ NMR

Note2: $^{13}C$ chemical shifts were assigned based on HSQC, and HMBC

**Supplementary Table 8 |** 1D NMR of collimonin D **2** in $C_2D_6OS$

| | Collimonin D **2** | |
|---|---|---|
| **No.** | $\delta_H$ (*J* in Hz) | $\delta_C$ |
| **1** | 11.04, brs | 175.0, COOH |
| **2** | 2.18, t, J = 7.20 | 34.0 |
| **3** | 1.47, m | 25.0 |
| **4** | 1.24,1.40, m | 25.5 |
| **5** | 1.24,1.40, m | 31.9 |
| **6** | 3.24, m | 73.1 |
| **7** | 3.35, m | 72.7 |
| **8** | 2.18, m; 2.31, m | 37.0 |
| **9** | 6.59, dt, J = 15.89, 7.35 | 152.3 |
| **10** | 5.75, d, J = 15.89 | 108.5 |
| **11** | | 77.0 |
| **12** | | 60.3 |
| **13** | | 72.3 |
| **14** | | 66.0 |
| **15** | | 68.3 |
| **16** | 3.93, s | 74.5 |

Note1: 600 MHz for $^1$H NMR

Note2: $^{13}$C chemical shifts were assigned based on HSQC, and HMBC

**Supplementary Table 9 |** 1D NMR of massilin A **3** in $C_2D_6OS$

| | Massilin A **3** | |
|---|---|---|
| **No.** | $\delta_H$ (*J* in Hz) | $\delta_C$ |
| **1** | 11.92, brs | 174.4, COOH |
| **2** | 2.18, t, J = 7.31 | 33.6 |
| **3** | 1.46, m | 24.4 |
| **4** | 1.25, m; 1.35, m | 24.6 |
| **5** | 1.28, m; 1.31, m | 36.6 |
| **6** | 3.34, m | 68.8 |
| **6-OH** | 4.43, d, J= 5.50 | |
| **7** | 1.35, m; 1.42, m | 35.6 |
| **8** | 2.18, m; 2.26, m | 29.3 |
| **9** | 6.60, dt, J = 15.86, 7.13 | 153.8 |
| **10** | 5.75, d, J = 15.86 | 106.7 |
| **11** | | 76.4 |
| **12** | | 59.6 |
| **13** | | 71.7 |
| **14** | | 65.5 |
| **15** | | 67.7 |
| **16** | 3.99, s | 74.0 |

Note1: 600 MHz for $^{1}H$ NMR

Note2: $^{13}C$ chemical shifts were assigned based on HSQC, and HMBC

**Supplementary Table 10 |** 1D NMR of massilin B **4** in $C_2D_6OS$

| | Massilin B **4** | |
|---|---|---|
| **No.** | $\delta_H$ (*J* in Hz) | $\delta_C$ |
| **1** | 11.94, br | 174.4, COOH |
| **2** | 2.19, t, J = 7.26 | 33.5 |
| **3** | 1.48, m | 24.2 |
| **4** | 1.26,1.40, m | 25.0 |
| **5** | 1.26,1.40, m | 31.4 |
| **6** | 3.24, m | 72.5 |
| **6-OH** | 4.37, d, J= 5.92 | |
| **7** | 3.35, m | 72.3 |
| **7-OH** | 4.53, d, J= 6.0 | |
| **8** | 2.18, m; 2.31, m | 36.2 |
| **9** | 6.43, dt, J = 15.8, 7.4 | 148.5 |
| **10** | 5.76, d, J = 15.8 | 108.8 |
| **11** | | 81.5 |
| **12** | | 71.8 |
| **13** | | 74.3 |
| **14** | | 79.8 |
| **15** | 6.04, dd, J= 11.38, 17.55 | 115.8 |
| **16** | $H_\alpha$, 5.76, dd, J= 11.38,1.96<br>$H_\beta$, 5.85, dd, J= 17.55, 1.96 | 131.5 |

Note1: 600 MHz for $^1$H NMR

Note2: $^{13}$C chemical shifts were assigned based on HSQC, and HMBC

### Supplementary Methods

#### Chemicals, strains, plasmids, and culture conditions

ACS grade ethyl acetate, dimethyl sulfoxide, HPLC grade of methanol, isopropanol and acetonitrile, and LCMS grade acetonitrile were purchased from J. T. Baker (Thermo Fisher Scientific, UK). Trifluoroacetic acid and LCMS grade formic acid were purchased from Sigma (Sigma-Aldrich, USA). Doxycycline was purchased from Cyrusbioscience, Inc. (Taiwan), and other antibiotics used in this study were purchased from Sigma (Sigma-Aldrich, USA). Culture media were purchased from BD Difco™. Coenzyme A was purchased from TRC (C636400; Toronto Research Chemicals, Canada). Acetyl-CoA (SI-A2056) and 7-diethylamino-3-(4-maleimidophenyl)-4-methylcoumarin (CPM; SI-C1484) were purchased from Sigma-Aldrich (St. Louis, MO, USA). Atorvastatin (075942) was purchased from Matrix Scientific Inc., USA), and amphotericin B (SI-A4888) was purchased from Sigma-Aldrich (St. Louis, MO, USA). Sequencing Grade Modified Trypsin (V5111) was purchased from Promega (Madison, WI, USA). All reagents used in crystallization were purchased from Hampton Research (Hampton Research, USA).

The strains, plasmids, and primers used in this study were listed in **Supplementary Tables 3-5**. *Massilia* sp. YMA4 was isolated from a marine sediment core collected by research vessel Ocean Researcher NO3 at Lamay island offshore, Pingtung County, Taiwan, on November 9, 2013 (OR3-1727). The voucher specimen (BCRC 81003) was deposited in the Bioresource Collection and Research Center (BCRC), Food Industry Research and Development Institute, Taiwan. *Candida albicans* ATCC 18804 type strain (BCRC 20512) was purchased from BCRC. Clinical *Candida* isolates were collected from National Taiwan University Hospital and provided by Prof. Ching-Hsuan Lin, National Taiwan University[4]. *E. coli* S17-1 λ *pir*. is a generous gift from Prof. Nai-Chun Lin, National Taiwan University. Human prostate PC-3 cell line was provided by Dr. Pei-Wen Hsiao, Agricultural Biotechnology Research Center, Academia Sinica, Taiwan.

For strain maintenance, all *E. coli* strains were cultured in Luria-Bertani (LB) broth at 37 °C with the corresponding antibiotic supplement, *Massilia* sp. YMA4 was cultivated in yeast and malt extract broth (YMB) consisting of 3 g/L yeast extract; 3 g/L malt extract; 10 g/L Dextrose; 5 g/L Peptone, and 20 g/L Bacto agar (for solid medium) at 30 °C. *Candida* strains were cultured in yeast extract-peptone-dextrose medium (YPD) consisting of 10 g/L yeast extract, 20 g/L Peptone, 20 g/L Dextrose, and 20 g/L Bacto agar (for solid medium) at 30 °C.

**Antagonism assay**

The 12 hours YMB broth (30 °C) of *Massilia* sp. YMA4 and Δ*masH* strain were respectively transplanted to PDA and incubated at 30 °C for two days. Then, overnight YPD broth culture of *Candida* strains was smeared perpendicular to the precultured *Massilia* sp. YMA4 and Δ*masH* strain on the PDA plate and continued to incubate for two days at 30 °C. After taking a photo with the camera, *Candida* cells were collected for quantitative RT-PCR.

**Quantitative RT-PCR**

The total RNA of *C. albicans* from the antagonism assay was collected using NautiaZ Bacteria/Fungi RNA Mini Kit (Nautia Gene Co., Ltd., Taiwan). The total RNA concentration was measured using NanoDrop 1000 Spectrophotometer (Thermo Fisher Scientific, USA). Transcriptor First Strand cDNA Synthesis Kit (Roche) was used to synthesis cDNA from the total RNA samples. The TOOLS 2xSYBR qPCR Mix Kit (BIOTOOLS Co., Ltd., Taiwan) and CFX ConnectTM Real-Time System (Bio-Rad Laboratories, USA) were used to perform quantitative RT-PCR. The experiment was independently repeated at least three times, and the means of the replicates are shown. The expression level of the target genes was normalized to the expression of *ACT1* housekeeping gene. Primers for this experiment were listed in (**Supplementary Table 3**).

**Construction of polyyne biosynthesis gene-null mutant strain Δ*masH***

The DNA fragment containing the *masH* fragment was amplified using primers *masH*-fF and *masH*-fR. The resulted PCR product was first cloned into yT&A vector (YC013; Yisheng Biotechnology Development Co., Ltd., Taiwan) and subcloned into pCM184 by restriction-ligation at *Kpn*I and *Pst*I site (New England Biolabs) to generate pCM184-Δ*masH* plasmid. The pCM184-Δ*masH* plasmid was further transformed into *E. coli* S17-1 for conjugation. The overnight culture of *Massilia* sp. YMA4 and *E. coli* S17-1 with pCM184-Δ*masH* were mixed with a 1:1 (v:v) ratio of optical density and further cultured on YMA plate. The conjugants colonies were selected using oxytetracycline and kanamycin and checked by PCR with mutant examination primer sets (**Supplementary Figure 17**).

**Construction of inducible *ERG10* overexpression strains and *masL*$_{opt}$ heterologous expression strain in *C. albicans***

*ERG10* fragment was amplified with *ERG10*-F, *ERG10*-R, and *ERG10His*-R. The *masL*$_{opt}$ fragment was optimized and synthesized by BIOTOOLS Co., Ltd., Taiwan. The fragments were respectively cloned into the pNIM1 vector for creating pNIM-ERG10, pNIM-ERG10L$_{127S,}$ and pNIM-MasL$_{opt}$ plasmids. The expression module was first linearized by *Apa*I/*Sac*II digestion and transformed into *Candida albicans* ATCC18804 using lithium acetate. The inducible *EGR10* overexpression strains and the *masL*$_{opt}$ heterologous expression strain were generated by replacing the single allele of *ADH1* with the linearized fragments (**Supplementary Figure 18**). The completed strains were selected on YPD plates with nourseothricin (Werner Bioagents, Jena, Germany) according to the previous research[6].

**Construction of inducible *masL* and *ACAT1* heterologous expression strains in *E. coli***

The gene *masL* from *Massilia* sp. YMA4 and the gene *ACAT1* (transition peptide truncated, aa 34 - 427) from human prostate PC-3 cell line were cloned into the site between *Nde*I/*Hind*III of vector pET28a(+) or *Nde*I/*Xho*I of vector pET22b(+) by In-Fusion cloning (Takara Bio USA, Inc.). The plasmids pET22b-MasL and pET28a-ACAT1 were transformed into *E. coli* C41(DE3) (Yeastern Biotech Co., Ltd., Taiwan) via heat shock and selected with ampicillin and kanamycin, respectively, for overexpressing recombinant hexahistidine-tagged proteins.

**Expression and purification of MasL, ERG10, and ACAT1**

For the expression of MasL and ACAT1, a glycerol stock culture of *E. coli* C41(DE3) harboring the selected plasmids was used to inoculate an overnight culture (37 °C, 200 rpm) of LB, supplemented with appropriate antibiotics. The culture was then used to inoculate 1 L 2xYT broth consisting of 16 g/L Tryptone; 10 g/L Yeast extract; 5 g/L NaCl and incubated at 37 °C, 200 rpm for ~3 h (OD600 = 0.4 - 0.6). Protein expression was induced upon the addition of 0.5 mM isopropyl-β-D-thiogalactopyranoside (IPTG), and then the cultures were incubated at 16 °C, 200 rpm for a further 16 h. The cells were harvested via centrifugation (4000 × g, 4 °C, 15 min) and resuspended in 30 mL of lysis buffer (50 mM Tris·HCl pH 8.5, 500 mM NaCl, 30 mM imidazole, and 300 μg/mL lysozyme) and lysed for 30min at 37 °C by gently inverting. The lysed cells were further lysed by sonication (Misonix Sonicator

XL2020) in 10-second intervals and were centrifuged (10,000 × g, 4 °C, 30 min).

For the expression of ERG10$_{L127S}$, a glycerol stock culture of P$_{tet}$-*ERG10$_{L127S}$-His* was used to inoculate an overnight culture (37 °C, 200 rpm) of YPD, supplemented with nourseothricin. The culture was then used to inoculate 4 L YPD broth and incubated at 37 °C, 200 rpm for ~3 h (OD600 = 0.4 - 0.6). Protein expression was induced upon the addition of 40 mM doxycycline, and then the cultures were incubated at 30 °C, 200 rpm for a further 16 h. The cells were harvested via centrifugation (4000 × g, 4 °C, 15 min) and resuspended in 30 mL of lysis buffer (50 mM Tris·HCl pH 8.5, 500 mM NaCl and 30 mM imidazole), by following lysed by a French press operated at 4 °C (20 kpsi) (CONSTANT SYSTEM RCB411) and centrifuged (10,000 × g, 4 °C, 30 min).

The protein supernatants were loaded on a 5 mL HisTrap HP column (GE Healthcare Bio-Sciences, USA) pre-equilibrated with lysis buffer and washed with three column volume (CV) of wash buffer (50 mM Tris·HCl pH 8.5, 500 mM NaCl, and 30 mM imidazole), and eluted with 20 CV elution buffer with a gradient concentration of imidazole (50 mM Tris·HCl pH 8.5, 500 mM NaCl, with imidazole from 50 to 500 mM). The eluents were ultra-filtrated and buffer-exchanged to gel-filtration buffer (20 mM Tris·HCl pH8.5, 100 mM NaCl) using Amicon Ultra-15 centrifugal filter units (Millipore) with a 30 kDa cutoff (MWCO). The His-tag purified proteins were further loaded on Superdex 200 Increase 10/300 GL column (GE Healthcare Bio-Sciences, USA) for size-exclusion chromatography with gel-filtration buffer (20 mM Tris·HCl pH8.5, 100 mM NaCl) in isocratic method. The eluents were subjected to SDS-PAGE analysis. Target protein-contained eluents were combined, ultra-filtrated, and then added glycerol to a final 10% for storing and further crystallization and enzyme assay. All the purification processes were performed at 4 °C. All chromatography was performed on ÄKTA pure 25 M1 purification system. The concentration of purified protein was measured using absorbance at 280 nm by NanoDrop 1000 Spectrophotometer (Thermo Fisher Scientific, USA) with calculated Extinction coefficients (ε)[7]: MasL (20970 $M^{-1}$ $cm^{-1}$), ERG10$_{L127S}$ (15930 $M^{-1}$ $cm^{-1}$) and ACAT1 (24410 $M^{-1}$ $cm^{-1}$).

**Genome sequence and annotation**

Genomic DNA of *Massilia* sp. YMA4 was extracted from the cultures grown at 30 °C in YMB using a genomic DNA purification kit (QIAgen,

Germany) following the manufacturer's instructions. The genomic DNA (total 20 µg) was sequenced by the PacBio RS II system (Pacific Biosciences Inc., USA). A 10-kb SMRTbell library was generated using a DNA Template Prep Kit 2.0 (10 Kb - 20 Kb; Pacific Biosciences Inc., USA), following the manufacturer's instructions. Sequencing was performed with a PacBio RS II sequencer (Pacific Biosciences Inc., USA) using one SMRT cell and P6-C4 chemistry at 360 min movie length. Single molecule real-time reads (159,840 filtered subreads and a mean length of 11,850 bp) were *de novo* assembled using the Hierarchical Genome Assembly Process workflow in SMRT analysis software version 3 (Pacific Biosciences Inc., USA)[8]. Genome annotation was automatically performed NCBI Prokaryotic Genome Annotation Pipeline (PGAP)[9]. The circular genome was upload to NCBI (GCA_003293715.1)

**RNA sequencing and transcriptomic analysis**

*Massilia* sp. YMA4 was activated in 4 mL YMB for 24 hours, and 500 µL of activated *Massilia* sp. YMA4 broth was then transferred into 50 mL YMB in a 250 mL flask and cultured for 24 hours. All of the inoculated broth was cultured under 30 $^{o}$C with 150 r.p.m. *Massilia* sp. YMA4 broth was then transplanted to YMA or PDA by sterilized swabs and kept under 30 °C for 48 hours. *Massilia* sp. YMA4 cells were collected in TRIzol® reagent (Thermo Fisher Scientific, USA). The method of RNA extraction was followed the TRIzol® reagent protocol. The RNA extracts were treated with DNAse I for 15 minutes in ambient to clean up the genomic DNA, and then the purified total RNA was acquired by using a MinElute PCR purification kit (Quiagen). TruSeq Stranded Total RNA with Ribo-Zero kits (Illumina) was used for mRNA-seq library preparation, then sequenced with paired-end reads (2 × 250 bp) using the Illumina MiSeq system.

After sequencing, adapters in raw read were trimmed by MiSeq software system. Quality and length trimming was performed by CLC genomics workbench (version 11, CLC bio) with default settings. The reads were mapped into the assembled *Massilia* sp. YMA4 genome (GCA_003293715.1) for following RPKM quantitation.

The differential gene expression analysis was analyzed through CLC software with default pipeline and settings. Identification of differentially expressed genes (DEGs) with the expression (|Fold change (FC)| ≥ 2 with p-value < 0.05) was based on RPKM and analyzed using the Empirical analysis method. Two biologic replicates of each condition were analyzed.

The enrichment analysis of Kyoto Encyclopedia of Genes and Genomes (KEGG)[10] pathway for DEGs was performed using R software with clusterProfiler package[11].

**Extraction of *Massilia* sp. YMA4 and UPLC-DAD-MS/MS analysis**

The *Massilia* sp. YMA4 was cultured as described in the antagonism assay. The agar plates were gridding and extracted with ethyl acetate (EA) 2-3 times. The extracts were concentrated and replaced with dimethyl sulfoxide (DMSO), preventing polyyne degradation while drying or resuspending with DMSO for LC-MS/MS analysis.

All samples were adjusted to 10 mg/mL and analyzed by using an Agilent 1290 Infinity II ultra-performance liquid chromatography (UPLC) system coupled to an Agilent 1260 Infinity II DAD HS system and to the Dual AJS electrospray ionization (ESI) source of 6545XT AdvanceBio LC/Q-TOF (quadrupole time-of-flight) mass spectrometer (Agilent Technologies, USA). The chromatographic separation was performed on an ACQUITY BEH C18 UPLC column (2.1 × 100 mm, 1.7 μm; Waters, USA) with a flow rate of 0.4 mL/min and a column temperature at 40 °C. Mobile phase A was 1‰ formic acid in water, and mobile phase B was 1‰ formic acid in acetonitrile (LCMS grade). The gradient elution condition for polyyne profiling analysis was as follows: Initially, mobile phase B was held at 5% for 1 min, then changed from 5 to 100% linearly in 10 min. The mobile phase B was then held at 95% for 2 min. Finally, it decreased to 5% in 0.2 min and held on for 2.8 min. The gradient elution condition for collecting tandem mass spectra of polyynes was as follows: Initially, mobile phase B was held at 5% for 1min, followed by 5 to 30% linearly in 3 min, subsequent increased to 35% gradually from 4 to 8 min and then increased to 100% from 8 to 15 min. The mobile phase B was then held at 95% for 2 min. Finally, it decreased to 5% in 0.2 min and held on for 2.8 min.

The diode array detector (DAD) was setting as full-spectrum scanning with the UV range from 220 to 500nm and scan rate 0.2 sec/spectrum. The electrospray ionization (ESI) source was used, and the analyses were performed in negative ion mode. Data were collected in centroid mode with *m/z* 100–700 and scan rate 330 ms/spectrum for HRMS acquisition. For HR-MS/MS acquisition, automated data-dependent acquisition (DDA) mode was performed with *m/z* 100–700 and scan rate 330 ms/spectrum for precursor ion scanning. The top three selected precursor ions were fragmented with collision energy set at 10 eV and an isolation window of 1.3

*m/z*. Data processing and peak identification was performed by using Agilent MassHunter software (version B.08.00).

**Isolation, structure elucidation, and quantitation of compounds 1-4**

NMR spectra were recorded on a Bruker Ascend 600 NMR spectrometer with a Prodigy cryoprobe operating at 600 MHz ($^1$H) using dimethyl sulfoxide-$d_6$ (99.9% D; Cambridge Isotope Laboratories, Inc., USA). NMR spectra were processed using Bruker Topspin (version 3.6) and MestReNova (Mestrelab, version 14.0.0). Spectra were referenced to residual solvent signals with resonances at $\delta_H$ 2.50 for dimethyl sulfoxide-$d_6$.

In general, the EA extract of *Massilia* sp. YMA4 on PDA plates was concentrated (not completely dried) and fractionated by Biotage Isolera One flash purification system. An HP silica column (20 μm, 12g, GRACE Inc., Columbia, MD, US) was used with a flow rate of 12 mL/min. The solvent system consisted of A (hexane), B (ethyl acetate) and C (methanol) with gradient solvent elution programmed as fellows: 0-4 min, 100% A; 4- 20 min, 0-60% B; 20-23 min, 60-100% B; 23-30 min, 100% B; 30-35, 0-100% C; 35-42 min, 100% C. The polyyne fractions were monitored by UPLC-DAD-MS/MS and combined as a polyyne-enriched fraction. The enriched fraction was further purified by reversed-phase-high performance liquid chromatography (RP-HPLC) on Hitachi LaChrom Elite HPLC system with a Hitachi L-2130 pump and L-2455 Diode-array Detector (Hitachi, Japan). The RP-HPLC separation was performed on Discovery HS C18 HPLC column (25 cm × 10 mm, 5 μm; SUPELCO Inc., USA) with a flow rate of 4.25 mL/min and a column temperature at RT The mobile phase A was 1‰ trifluoroacetic acid in water. The mobile B was 1‰ trifluoroacetic acid in acetonitrile (HPLC grade). The gradient elution condition for enriched fraction isolation was as follows: Initially, the concentration of B was changed from 20 to 44% linearly in 27 min, next increased to 100% immediately from 27 to 28 min. The ratio of mobile phase B was then held at 100% for 7 min. Finally, it decreased to 20% in 1 min and held on for 9 min to make a balance.

For further purification of collimonin C **1**, collimonin D **2,** and massilin B **4**, each polyyne fraction was purified by RP-HPLC. The RP-HPLC separation was performed on Discovery HS C18 HPLC column (25 cm × 10 mm, 5 μm; SUPELCO Inc., Missouri, US) with a flow rate of 3.25 mL/min at RT. The mobile phase A was 1‰ trifluoroacetic acid in water. The mobile B was 1‰ trifluoroacetic acid in acetonitrile/isopropanol (3:7, HPLC grade). The gradient elution condition for enriched fraction isolation was as follows:

Initially, the concentration of B was changed from 30 to 42.5% linearly in 25 min, following an increase to 100% from 25 to 30 min. The ratio of mobile phase B was then held at 100% for 5 min.

For further purification of massilin A **3**, the RP-HPLC separation was performed on Discovery HS C18 HPLC column (25 cm × 10 mm, 5 μm; SUPELCO Inc., US) with a flow rate of 3.25 mL/min at RT. The mobile phase A was 1‰ trifluoroacetic acid in water. The mobile B was 1‰ trifluoroacetic acid in acetonitrile/isopropanol (3:7, HPLC grade). The gradient elution condition for enriched fraction isolation was as follows: Initially, the concentration of B was changed from 38.5 to 51% linearly in 25 min, subsequently increased to 100% from 25 to 30 min. The ratio of mobile phase B was then held at 100% for 5 min.

All the purified polyynes were stored in methanol. Polyynes for all experiments were prepared in fresh, and the solvent was replaced by dimethyl sulfoxide-$d_6$ for the following experiments.

The physicochemical properties of collimonin C **1** and collimonin D **2** were identical with the report from *Collimonas fungivorans* Ter331[12]. The UV absorption at 272, 288, 307, 329 nm suggests that collimonin C **1**, collimonin D **2**, and massilin A **3** were functionalized with enetriyne moiety. The UV absorption at 274, 290, 310 nm suggests massilin B **4** was functionalized with enediyne -ene moiety.

**Collimonin C (1)**: light yellow dissolved in DMSO; $^1$H NMR (dimethyl sulfoxide-$d_6$, 600 MHz) data **Supplementary Table 7**; HRMS(ESI$^-$) *m/z* 273.1138 [M-H]$^-$ (calcd. for $C_{16}H_{17}O_4$, 273.1132); Double bond equivalents (DBE) = 8.5.

**Collimonin D (2)**: light yellow dissolved in DMSO; $^1$H NMR (dimethyl sulfoxide-$d_6$, 600 MHz) data **Supplementary Table 8**; HRMS(ESI$^-$) *m/z* 273.1139 [M-H]$^-$ (calcd. for $C_{16}H_{17}O_4$, 273.1132); Double bond equivalents (DBE) = 8.5.

**Massilin A (3)**: racemate, light yellow dissolved in DMSO; $^1$H NMR (dimethyl sulfoxide-$d_6$, 600 MHz) data **Supplementary Table 9**; HRMS(ESI$^-$) *m/z* 257.1189 [M-H]$^-$ (calcd. for $C_{16}H_{17}O_3$, 257.1183); Double bond equivalents (DBE) = 8.5.

**Massilin B (4)**: light yellow dissolved in DMSO; $^1$H NMR (dimethyl sulfoxide-$d_6$, 600 MHz) data **Supplementary Table 10**; HRMS(ESI$^-$) *m/z* 275.1298 [M-H]$^-$ (calcd. for $C_{16}H_{19}O_4$, 275.1289); Double bond equivalents (DBE) = 7.5.

The [13]C chemical shifts of all polyynes were assigned based on HSQC and HMBC. The quantitation of polyynes was measured in 1H NMR using the $^{13}C$-coupling satellite peak of dimethyl sulfoxide-$d_6$ (~1.1% of $^{12}C$; 2.39/2.61)[13] with an equation: $C_{polyyne} = \frac{I_{polyyne}}{I_{DMSO-d6}} \times \frac{N_{DMSO-d6}}{N_{polyyne}} \times C_{DMSO-d6}$

*C* is the concentration, *I* is integral of proton signal, and *N* is the number of nuclei giving rise to the signal. The purity was measured using HPLC-DAD with the same gradient condition for purification at 241 nm with over 95% purity (integrated area).

**Sequence alignment and structure superimposition**

The protein sequence alignment was performed in the CLC genomics workbench (version 11, CLC bio) with gap open cost 10 and extension cost 1.0. The protein superimposition was processed and produced figures in Maestro (Schrödinger Release 2021-1: Maestro, Schrödinger, LLC, USA).

**Transmission Electron Microscope**

*C. albicans* was cultured in YPD started with OD600 = 0.05 at 37 °C for 2 hours. The broth cultures were independently treated with 1% ethyl acetate (EA), 1 mg/mL Δ*masH* crude extract (1% EA), and 1 mg/mL *Massilia* sp. YMA4 crude extract (1% EA) and then cultured at 37 °C for 4 hours. All samples were obtained, and a similar fixation process followed the previous study and modified using a 0.1 M phosphate buffer[14]. Finally, the samples were embedded in pure Spurr's resin and processed with ultrathin section, then observed under an FEI Tecnai™ $G^2$ F20 S-TWIN transmission electron microscope (FEI Company, USA).

### Supplementary Note

#### Background of kinetic evaluation of irreversible inhibitors and polyynes-MasL experiment detail

The measurement and calculation of polyynes inhibition kinetic refer to the previous covalent inhibitor model[15]. The covalent inhibitors described in this paper follow a two-step binding. Initial non-covalent binding to the enzyme under rapid equilibrium conditions is followed by slower covalent bond formation (equation 1). The observed rate constant for inhibition ($k_{obs}$) is a pseudo-first-order rate constant obtained from the product formation by fitting the kinetic data to equation 2. For irreversible covalent inhibitors, kinetic parameters $k_{inact}$ (rate of inactivation, maximum $k_{obs}$ at infinite inhibitor concentration) and $K_I$ (concentration inhibitor that yields a half-maximum $k_{obs}$; ½ $k_{inact}$) can be obtained from plotting $k_{obs}$ values as a function of [inhibitor], and fitting to equation 3. The $k_{inact}$ /$K_I$ ratio represents the efficiency of inhibition in Table 1 of the main manuscript.

$$\mathrm{E + I} \underset{}{\overset{K_I}{\rightleftharpoons}} \underset{\text{Initial Non-covalent complex}}{\mathrm{E * I}} \xrightarrow{k_{inact}} \underset{\text{Final covalent complex}}{\mathrm{E\text{-} I}} \quad (1)$$

$$\text{Total Occupancy} = [\mathrm{E\text{-}I}] / [\mathrm{E}]_{int} = 1 - ( [\mathrm{E}]_{free} / [\mathrm{E}]_{int} ) = 1 - e^{-(k_{obs} * t_I)} \quad (2)$$

$$k_{obs} = \frac{k_{inact}\,[\text{Inhibitor}]}{K_I + [\text{Inhibitor}]} \quad (3)$$

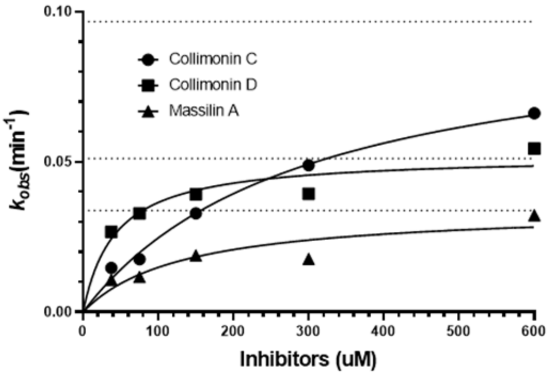


| One site -- Specific binding | Collimonin C | Collimonin D | Massilin A |
|---|---|---|---|
| **Best-fit values** | | | |
| $K_{inact}$ (min$^{-1}$) | 0.09798 | 0.05208 | 0.03449 |
| $K_I$ (uM) | 297.10 | 42.84 | 132.10 |
| **95% CI (profile likelihood)** | | | |
| $K_{inact}$ (min$^{-1}$) | 0.06041 to 0.2239 | 0.03360 to 0.08768 | 0.02291 to 0.06074 |
| $K_I$ (uM) | 103.7 to 1122 | -3.562 to 243.0 | 37.22 to 463.1 |
| **Goodness of Fit** | | | |
| Degrees of Freedom | 38 | 43 | 37 |
| R squared | 0.4723 | 0.06669 | 0.3073 |
| Sum of Squares | 0.01299 | 0.04369 | 0.003650 |
| Sy.x | 0.01849 | 0.03188 | 0.009932 |

CI: Confidence interval.

The progress curves for $k_{inact}$ / $K_I$ relationship between inhibitor concentration were recorded with CPM-labeling for coenzyme A production as acetyl-CoA acetyltransferase's residual activity. The inhibition reaction started from prior incubation of 10 μM MasL with various concentration of polyynes (37.5, 75, 150, 300, 600 μM) for 15 and 30 min ($t_{1/2}^{\infty}$ of 15 and 30 min), followed by enzyme reaction and CPM-labeling for recorded the residual activity to calculate total occupancy. The progress curves used the hyperbolic regression model (Graphpad Prism 8, USA) to calculate the kinetic inhibition parameters. Each reaction point (time and concentration) was recorded at least three replicates for model building.

#### Supplementary References


1. Bloxham DP, Chalkley RA, Coghlin SJ, Salam W. Synthesis of chloromethyl ketone derivatives of fatty acids. Their use as specific inhibitors of acetoacetyl-coenzyme A thiolase, cholesterol biosynthesis and fatty acid synthesis. *Biochemical Journal* **175**, 999-1011 (1978).

2. Holland PC, Clark MG, Bloxham DP. Inactivation of pig heart thiolase by 3-butynoyl coenzyme A, 3-pentynoyl coenzyme A, and 4-bromocrotonyl coenzyme A. *Biochemistry* **12**, 3309-3315 (1973).

3. Palmer MA*, et al.* Biosynthetic thiolase from Zoogloea ramigera. Evidence for a mechanism involving Cys-378 as the active site base. *The Journal of biological chemistry* **266**, 8369-8375 (1991).

4. Lo W-H, Deng F-S, Chang C-J, Lin C-H. Synergistic Antifungal Activity of Chitosan with Fluconazole against Candida albicans, Candida tropicalis, and Fluconazole-Resistant Strains. *Molecules* **25**, 5114 (2020).

5. Marx CJ, Lidstrom ME. Broad-host-range cre-lox system for antibiotic marker recycling in gram-negative bacteria. *BioTechniques* **33**, 1062-1067 (2002).

6. Park Y-N, Morschhäuser J. Tetracycline-inducible gene expression and gene deletion in Candida albicans. *Eukaryot Cell* **4**, 1328-1342 (2005).

7. Wilkins MR*, et al.* Protein Identification and Analysis Tools in the ExPASy Server. In: *2-D Proteome Analysis Protocols* (ed Link AJ). Humana Press (1999).

8. Chin CS*, et al.* Nonhybrid, finished microbial genome assemblies from long-read SMRT sequencing data. *Nat Methods* **10**, 563-569 (2013).

9. Tatusova T*, et al.* NCBI prokaryotic genome annotation pipeline. *Nucleic Acids Research* **44**, 6614-6624 (2016).

10. Kanehisa M, Furumichi M, Tanabe M, Sato Y, Morishima K. KEGG: new perspectives on genomes, pathways, diseases and drugs. *Nucleic Acids Res* **45**, D353-D361 (2017).

11. Yu G, Wang LG, Han Y, He QY. clusterProfiler: an R package for comparing biological themes among gene clusters. *Omics : a journal of integrative biology* **16**, 284-287 (2012).

12. Kai K, Sogame M, Sakurai F, Nasu N, Fujita M. Collimonins A–D, Unstable Polyynes with Antifungal or Pigmentation Activities from the Fungus-Feeding Bacterium Collimonas fungivorans Ter331. *Organic Letters* **20**, 3536-3540 (2018).

13. Moutzouri P, Kiraly P, Phillips AR, Coombes SR, Nilsson M, Morris GA. (13)C Satellite-Free (1)H NMR Spectra. *Analytical chemistry* **89**, 11898-11901 (2017).

14. Vazquez-Muñoz R, Avalos-Borja M, Castro-Longoria E. Ultrastructural Analysis of Candida albicans When Exposed to Silver Nanoparticles. *PLOS ONE* **9**, e108876 (2014).

15. Strelow JM. A Perspective on the Kinetics of Covalent and Irreversible Inhibition. *SLAS discovery : advancing life sciences R & D* **22**, 3-20 (2017).
